# Enhancing the performance of Magnets photosensors through directed evolution

**DOI:** 10.1101/2022.11.14.516313

**Authors:** Armin Baumschlager, Yanik Weber, David Cánovas, Sara Dionisi, Mustafa Khammash

## Abstract

Photosensory protein domains are the basis of optogenetic protein engineering. These domains originate from natural sources where they fulfill specific functions ranging from the protection against photooxidative damage to circadian rhythms. When used in synthetic biology, the features of these photosensory domains can be specifically tailored towards the application of interest, enabling their full exploitation for optogenetic regulation in basic research and applied bioengineering. In this work, we develop and apply a simple, yet powerful, directed evolution and high-throughput screening strategy that allows us to alter the most fundamental property of the widely used nMag/pMag photodimerization system: its light sensitivity. We identify a set of mutations located within the photosensory domains, which either increase or decrease the light sensitivity at sub-saturating light intensities, while also improving the dark-to-light fold change in certain variants. For some of these variants, photosensitivity and expression levels could be changed independently, showing that the shape of the light-activity dose-response curve can be tuned and adjusted. We functionally characterize the variants *in vivo* in bacteria on the single-cell and the population levels. We further show that a subset of these variants can be transferred into the mOptoT7 for gene expression regulation in mammalian cells. We demonstrate increased gene expression levels for low light intensities, resulting in reduced potential phototoxicity in long-term experiments. Our findings expand the applicability of the widely used Magnets photosensors by enabling a tuning towards the needs of specific optogenetic regulation strategies. More generally, our approach will aid optogenetic approaches by making the adaptation of photosensor properties possible to better suit specific experimental or bioprocess needs.

## Introduction

In the field of bioengineering, light has emerged as a versatile input to steer and control various biological processes. It is crucial for photosynthetic processes or as protection against detrimental effects of electromagnetic radiation, such as photooxidative cellular damage. Organisms have evolved a variety of photosensitive proteins called “photoreceptors” or “photosensors”, which allow them to react to a light stimulus. Depending on their function, these domains can register different wavelengths and intensities of light. Light is usually absorbed through an organic chromophore, which induces structural changes within the apoprotein^1^. These structural changes revert back to the ground or “dark” state of the photoreceptor in a thermally driven reaction.

Optogenetics employs such photosensors or photosensory domains for the engineering of genetically encoded light-sensitive proteins^2^. Since the development of the first genetically engineered light-sensitive proteins^3^, light has been used in different optogenetic approaches to interfere with and control a variety of cellular functions^4,5^. The use of light as input is particularly attractive because it enables precise spatial and temporal applications,^6,7^ both with a higher resolution when compared to chemical induction. Furthermore, in nonphotosensitive organisms such as *Escherichia coli*, light is an orthogonal input that does not require uptake and/or conversion as with small molecule inducers.^8–10^ Thus, the use of light allows for minimally invasive spatiotemporal control and predictability.

Photosensitive domains are the key components of optogenetics. In general, optogenetic proteins comprise a photosensory domain as well as an actuating module.^2,4^ For example, many cellular sensors such as kinases initiate signaling and cellular responses through oligomerization.^11,12^ Such proximity-based regulation is also the basis of various natural light-sensing modules in which the interaction, and therefore the distance of the interaction partners, is controlled through light in a mechanism that leads to either homodimerization or oligomerization of the same photoregulator or heterodimerization and oligomerization of two or more different photoregulators. Such proximity control can be used to assemble inactive subcomponents into an active protein, or for recruitment of an active protein to a specific location of action. Light-controlled dimerization domains have been used to implement optogenetic control usually through homodimerization of e.g. receptors^13–19^ or heterodimerization for split proteins ^7,20–22^ as well as subcellular localization^3,23,24^.

A widely used class of photosensors are light, oxygen, and voltage (LOV) domains as they have features that are often especially attractive for optogenetic protein (Opto-protein) engineering, such as a small domain size and tunable kinetic properties. LOV domains employ flavin mononucleotide (FMN) and flavin adenine dinucleotide (FAD), which are present in most organisms, as chromophores. One of the most used photosensory protein heterodimerization pairs is the “Magnets”^25^, which are derived from the homodimerization protein VIVID (VVD). The LOV domain VVD, from the filamentous fungus *Neurospora crassa^26^*, has been used in numerous optogenetic designs (^13,18,27–38^). Blue light induces a conformational change in the LOV domain, which initiates homodimerization of two VVD domains and dimer stabilization.^26,39,40^ Based on a truncated version of this photosensory domain^26^, the Magnets heterodimerization system was created through rational protein engineering.^25^ For this, negatively and positively charged amino acids were introduced into the VVD dimer interface to create complementary pMag and nMag domains. We and others used these Magnets domains for reconstitution of split proteins^7,32–35^ due to their small size (150 amino acids), favorable structure in which N- and C-termini of the two domains come in close proximity in the dimeric state, and dark-state reversion rate tunability.^7^

The Opto-T7RNAP is a light-inducible transcription system that incorporates the VVD-based Magnets photoregulators into the heterologous T7 RNA polymerase (Figure 1A). The fusion of the two split parts to the Magnets domains created a gene expression system that shows high activation with blue light induction and simultaneously low residual expression in the dark.^7^ This difference in expression level with and without blue light is called the light-induced fold change. A crucial aspect for the use of this system as a screening system is that it offers a high dynamic range, which enables screening at sub-saturating induction levels (Figure 1 B, C). We characterized this transcription system via the expression of the red fluorescent protein mCherry (Figure 1A). Several mutations in the Magnets domains and VVD have been previously described using rational structure-guided protein engineering. These approaches enabled changing their kinetic properties^19,25,26,41–43^ or optimized functionality in mammalian cells^44^. Here, we envisioned that the direct phenotypic output of our transcription system in *E.coli* enables a high throughput directed evolution campaign, which allows one to screen for desired photoregulator properties. These properties range from photosensitivity alone to expression level alone, or a combination of both, all of which goes beyond previous protein engineering efforts.

**Figure 1:**
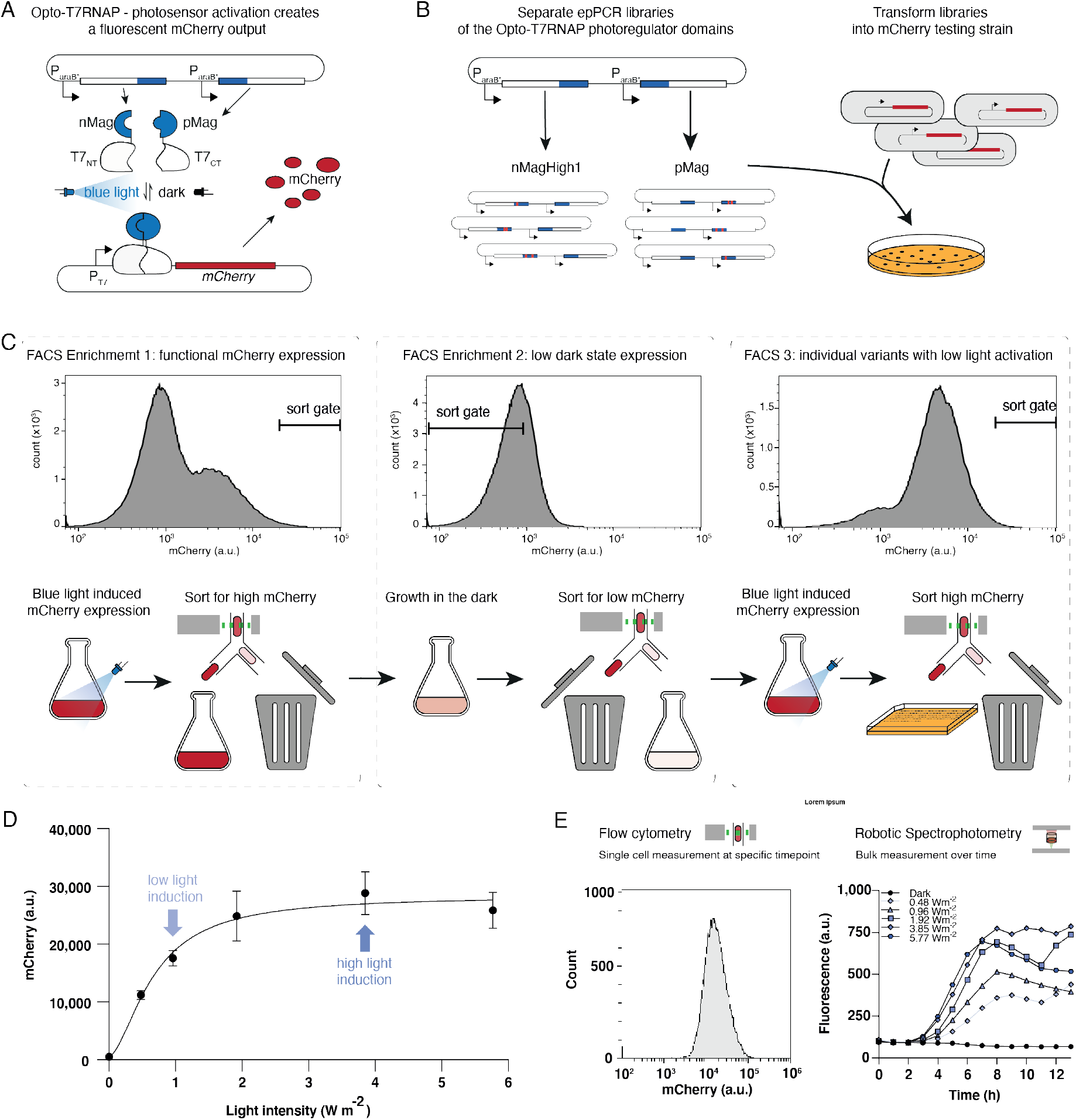
Vivid-based Magnet photosensor engineering using error-prone PCR and the optogenetic transcription system “Opto-T7RNAP” via Fluorescence Activated Cell Sorting (FACS). (A) Opto-T7RNAP^7^ is an engineered T7 RNA Polymerase (T7RNAP) that is blue-light activatable. It consists of two split fragments of the T7RNAP, which are fused to the light-inducible heterodimerizing Magnet domains. nMagHigh1 is fused to the C-terminus of the T7RNAP N-terminus and pMag is fused to the N-terminus of the C-terminal T7RNAP fragment. Light induces a conformational change in the Magnet domains which leads to the binding of the two complementary nMagHigh1 and pMag domains. The resulting spatial proximity of the T7RNAP fragments reconstitutes the function of the enzyme, leading to the transcription of genes from T7 promoters. The screening system consists of two plasmids, one containing the two Opto-T7RNAP genes under the control of arabinose-inducible promoters, and a reporter plasmid containing the red fluorescent protein mCherry expressed from the T7 promoter. mCherry fluorescence serves as a measurable output for Magnet heterodimerization. (B) Individual error-prone PCR libraries of nMagHigh1 and pMag photoregulators were constructed and transformed into an *E. coli* strain that contained the mCherry reporter plasmid. (C) Example FACS strategy to screen for variants with increased light sensitivity, meaning a higher mCherry expression level at non-saturating light-induction conditions. For the first enrichment (left) nMagHigh1 or pMag libraries were induced with nonsaturating light for 4h and then sorted for high mCherry expression, while cells with low or no mCherry fluorescence are discarded. This removes the bulk of the library in which mutations inactivated or reduced the light sensitivity of the photosensory domain. The enriched libraries were regrown in the dark and then sorted for low mCherry expression for the second enrichment (middle). The second step serves to select variants that still show a low dark state and removes constitutively active variants. These enriched libraries were then again induced for 4h with low-intensity 465-nm light blue light and single cells spotted on Omnitrays with LB-agar to isolate individual variants (right). As an example, the histograms of the mCherry expression profile of a nMagHigh1 library are shown with low-intensity light induction of the original mutagenesis library left, this enriched library regrown without light-induction in the middle for the second enrichment, and the second enriched library again induced with light-intensity light induction. All histograms further show exemplary sort gates which select for high mCherry expression with light-induction and low mCherry expression in the dark. (D) Dose-response curve of the original Opto-T7RNAP*(563) of mCherry expression level in response to blue light (465 nm) of different light intensities. To be able to screen for higher light-sensitive photoregulators, 0.96 Wcm^-2^ light intensity was used for sub-saturating induction, and 3.85 Wcm^-2^ for saturating light induction of the libraries. For characterization of identified variants, we used both non-saturating (0.96 Wcm^-2^) and saturating (3.85 Wcm^-2^) light induction. The diagram shows mean mCherry expression values and standard deviation of six (n=6) biological replicates measured after 5h incubation time in all cases. (E) Characterization of variants was performed on the single-cell level through flow cytometry (left) and in batch culture over time through spectrophotometry (right) exemplarily shown with the wild-type Opto-T7RNAP*(563) regulator. Single-cell mCherry expression profile is shown after 5h saturating light induction at 37°C. Spectrophotometric measurements in batch culture over 13h at 37°C. The diagram shows mean mCherry expression values of at least eight (n=8) biological replicates.

Specifically, in this work we aimed to alter the most fundamental property of this photosensory domain: its light sensitivity. Highly light-sensitive photosensors, for example, bear the advantage that lower intensity light can be applied to achieve similar outputs. This reduces potential phototoxic effects (e.g. mammalian cell lines^45^), aids in light-delivery into denser cell cultures^4,46^, and minimizes heat development by the light application. In other scenarios, lower light sensitivity might be preferred, e.g., if a regulator shall not be activated by ambient light or for multiplexing when combined with other, less sensitive, light controllers.

To allow for screening of Magnets variants with increased or decreased light-sensitivities, we used the Opto-T7RNAP*(568), which directly links the activity of the photosensitive domain to a detectable fluorescence output, thus creating a genotype-phenotype linkage that allows for screening of variants with altered photosensitivity. Through a single round of mutagenesis, we identified several mutations in both the nMagHigh1 as well as the pMag domain, some of which led to dramatically increased light sensitivities while maintaining or even improving the dark to light fold-change. While the mutations identified here will expand the applicability of these photosensory domains and enable a wider range of applications, we anticipate that our approach can also be used to identify mutations that change other properties such as dark-state reversion rate or enhancement of fold change, which will further expand the applicability and toolset of VVD-based photoregulators in particular, and optogenetic approaches in general.

## Results

### Library and experimental designs

We used error-prone PCR (epPCR) to diversify both of the photosensory domains (nMagHigh1 and pMag) of our Opto-T7RNAP*(563) individually (Figure 1B). Both Magnets domains are based on a 36-residue N-terminal truncated VVD photoregulator^26^, and differ in several amino acids that were rationally introduced to enable heterodimerization.^25^ nMagHigh1 is C-terminally fused to the N-terminal part of the T7RNAP, while pMag is N-terminally fused to the C-terminal part of the T7RNAP (Figure 1A). Although both photosensory domains are derived from VVD, we chose to create libraries for each individually (Figure 2A, inset; Figure 2B, inset), as they differ in amino acid positions I52 and M55 that confer complementary binding, as well as M135 and M165 for nMagHigh1.^25^ This leaves the possibility for different compensatory mutations for each domain, as well as accounts for their fusion to either the N- or C-terminal part of the T7RNAP.^7^ These libraries were then transformed separately into *E. coli* strain AB360^7^, which contained a plasmid for expression of mCherry under T7RNAP control (Figure 1B). We used two different ribosome binding sites (RBS) strengths, first the previously used RBS from pAB50^7^, as well a version denoted “pAB50-11k” in which we weakened the TIR using the RBS calculator 2.0^47^ to 15% of the original RBS (Supplementary Figure 1). The use of these two different expression strengths in separate libraries should allow one to find higher light-sensitive variants while limiting the effect of metabolic burden induced through mutations that cause higher mCherry expression levels.

**Figure 2:**
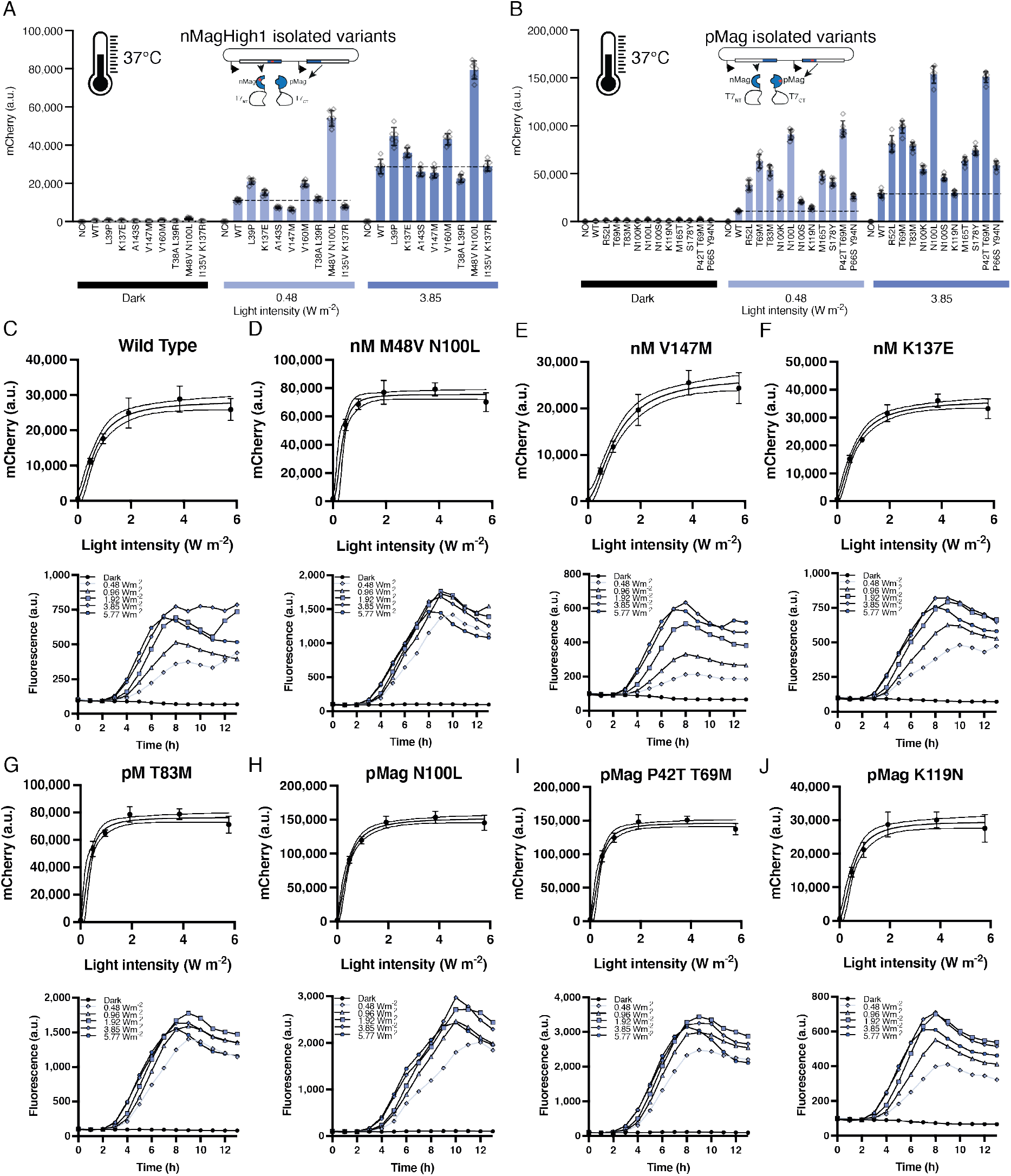
Characterization of identified mutations in nMagHigh1 (A) and pMag (B). Characterization of variants was performed through flow cytometry in comparison to the wild-type Opto-T7RNAP*(563) regulator and a negative control containing the mCherry expression plasmid and a second empty plasmid without the optogenetic regulator. mCherry expression values were acquired after 5h incubation at the indicated light intensity (465-nm light) and 37°C. Shown are the mean fluorescence values and standard deviation as well as individual data points of six (n=6) biological replicates. Fluorescence measurement over time through spectrophotometry at the indicated light intensity and 37°C of the wild type (C) and selected variants M48V N100L (D), V147M (E), K137E (F) of nMagHigh1 and T83M (G), N100L (H), P42T T69M (I) and K119N (J) of pMag. Shown are the mean fluorescence values of at least eight (n=8) biological replicates.

We chose photosensitivity as the target property for tuning for four main reasons: First, as previously mentioned, the ability to sense light effectively is *the* defining feature of a photoregulator, and engineering enzymes for the direct improvement of activities on their natural function is often unsuccessful and generally considered challenging^48^. Second, a potential drawback of optogenetic methods for the control of cells is light-induced cellular damage caused by high-intensity light and/or long illumination durations. While this effect might be less pronounced in bacteria like *E. coli*, it can be problematic in higher organisms such as mammalian cell lines. Third, depending on the light application setting, high light intensity can lead to the development of heat, which in turn might interfere with experiments or applications. Fourth, a set of photoregulators with different sensitivities for the same wavelength will enable the multiplexing of expression levels of different proteins, as well as the activation of genetic circuitry through variation of the light intensity.

### Library induction and screening strategy

We applied a multi-step Fluorescence-Activated Cell Sorting (FACS) for screening and selection of variant libraries with altered light sensitivities (Figure 1C). For this, we used different sequences of induction conditions: No light induction (further denoted “dark”), in which variants with mCherry expression comparable to the uninduced original Opto-T7RNAP*(563) were sorted for, or with light intensities of 0.96 and 3.85 W/cm^2^ (further denoted as low or high light induction respectively), which correspond to intermediate or saturating light induction (Figure 1D).

For FACS sorting, cells were grown for 4h either in the dark or induced with light before growth was arrested through cooling to 4°C on ice and resuspension of the cells in phosphate-buffered saline (PBS), as described in the Materials and Methods section. An example of the FACS protocol is shown in Figure 1C. In this example, the libraries were first enriched by inducing the cells with low-intensity light and sorting for functional mCherry expression. The majority of variants showed reduced-to-non-functional mCherry expression, similar to the negative control (Supplementary Figure 2, left), indicating that most mutations inactivated the function of the photosensor, as to be expected through random mutagenesis. The enriched libraries were then regrown in the dark and sorted for low mCherry expression (Figure 1C, middle panel). This second step excludes constitutively active variants as well as variants with significantly increased dark state expression. In the final step (Figure 1C, right panel), the libraries were again induced with low-intensity light and sorted for functional mCherry expression. Individual cells were spotted on LB-agar plates, which were regrown in M9 medium and screened. Individual mutations were identified by Sanger sequencing and recloned into the original plasmid. For final characterization, cultures were grown for 5h, either in the dark or light-induced, before inhibition and measurement through flow cytometry. In addition, main cultures were inoculated while growth and mCherry fluorescence were monitored periodically through spectrophotometry until stationary growth phase (Figure 1E, right; for WT) as described in the Materials and Methods.

### Identified nMagHigh1 and pMag variants

We compared the mCherry expression of the identified variants to the expression of the original wild-type Opto-T7RNAP*(563)^7^ and a negative control, containing only the mCherry reporter plasmid and an empty backbone with the same antibiotic resistance as the plasmid harboring the Opto-T7RNAP. For this, we took two experimental approaches: First, through single-cell analysis using flow cytometry measurement at a single time point, and second, through spectrophotometry over time.

Single-cell characterization of variants through flow cytometry was performed after 5h expression in log growth phase. This data served to extract changes in the basal dark (denoted *b*), as well as maximal light-induced (denoted *t*) mCherry expression. To assess changes in light sensitivities, we compared the light-intensities that lead to half-maximal gene expression (denoted *I50*), which were obtained from fits of the light-induced dose-response of mCherry expression to a Hill-type equation, as described in the Material and Methods section. Second, we also characterized the variants on the population level through spectrophotometry over time. The data were analyzed both using hierarchical clustering as well as through fits of the mCherry expression data of a single time point to a Hill-type equation to obtain the light-induced dose-response curve parameters.

After just one round of mutagenesis, we selected 19 different variants displaying a variety of interesting properties (Figure 2 at 37°C, Figure 3A-B at 30°C, and Figure 3C-D at 40°C; shown in more detail in Supplementary Figure 3 and Supplementary Figure 4). From these variants, 5 single and 3 double mutations were found in nMagHigh1, and 9 single and 2 double mutations were located in pMag (Supplementary Table 1). All calculated properties for *b, t* and *I50* as well as percentage changes compared to the WT are summarized in Supplementary Tables 2-12 with the underlying data shown in Supplementary Figures 5-20. Apart from these variants, we also isolated a double mutant in the pMag library (M135I, M165I), which was previously described as pMagHigh1^25^, and showed a higher mCherry expression level compared to pMag in the screening (data not shown), thereby confirming the ability of this setup and approach to identify variants with improved properties for a given experimental setup.

**Figure 3:**
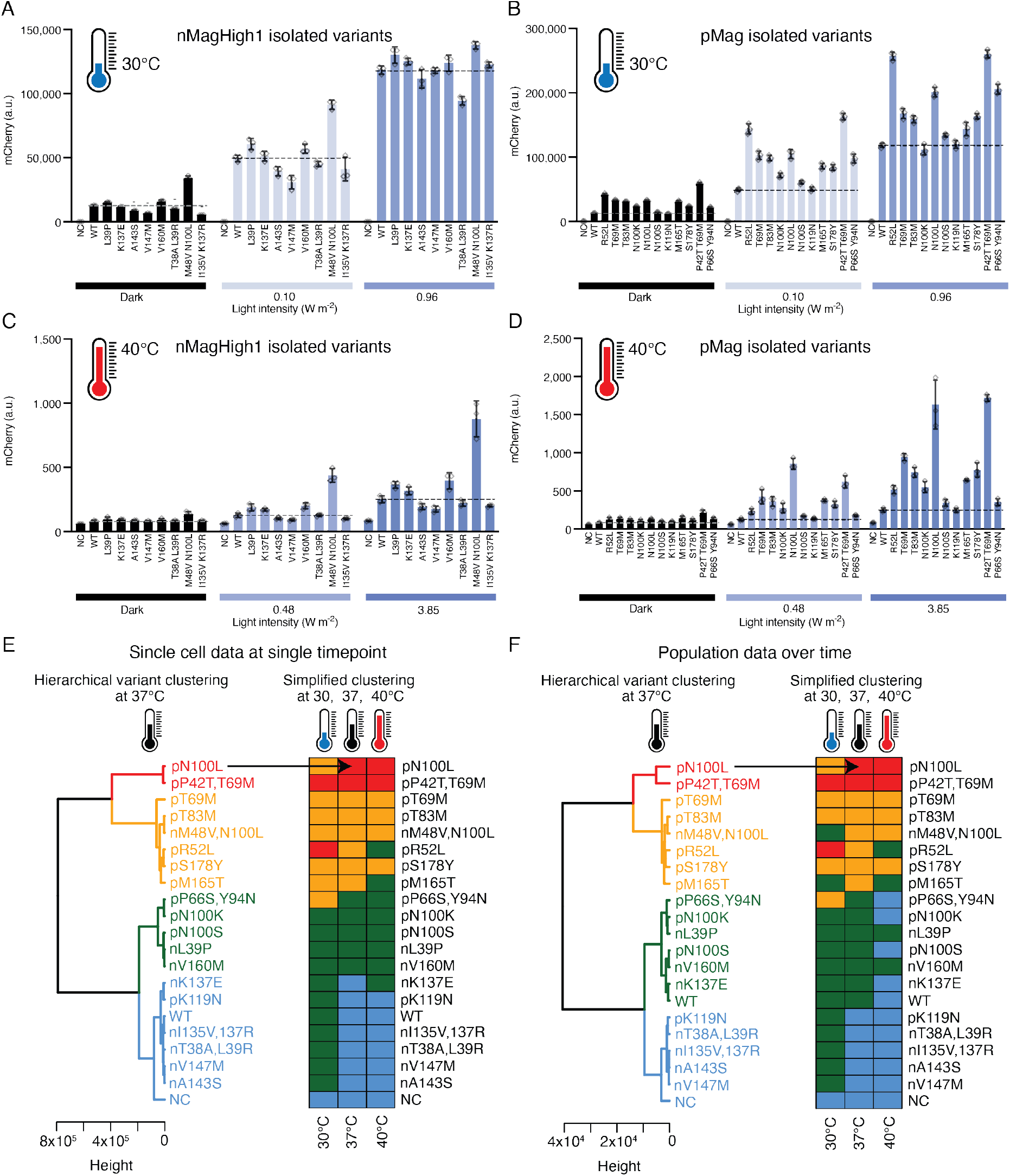
Characterization of identified photoregulator variants found in nMagHigh1(A, C) and pMag (B, D) at 30°C (A, B) and 40°C (C, D). Characterization of variants was performed through flow cytometry in comparison to the wild-type Opto-T7RNAP*(563) regulator and a negative control containing the mCherry expression plasmid and a second empty plasmid that does not contain the optogenetic regulator. mCherry expression values were acquired after 5h incubation at the indicated light intensity (465-nm light) and temperature. Shown are the mean fluorescence values and standard deviation as well as individual data points of three (n=3) independent biological replicates. (E, F) Hierarchical clustering using the Ward method and Euclidean distances of flow cytometry data at one single time point (E) and spectrophotometry data acquired every hour until stationary phase (F). Variants were clustered according to the expression levels into very high (red), high (orange), medium (green), and low (blue) expression levels. Dendrograms represent the clustering of the expression data obtained at 37°C. Lateral bars represent which expression cluster each variant belongs to at the indicated temperature for comparison.

Alongside manual comparison and the extraction of relevant enzyme parameters as mentioned above, we also performed principal component (PC) analysis of the datasets. The first PC could explain over 95% of the variance in the flow cytometry data (Supplementary Figure 21 A-C). In the case of the spectrophotometry data acquired over time, the first PC could explain only 62% of the variance (data not shown). We reasoned that if the data were not scaled before the PC analysis, this would prevent giving excessive weight to the fluorometric data obtained at low ODs values, when the bacterial cultures are starting to grow and gene expression could not be detected yet. Indeed, using non-scaled data the first PC could explain over 95% of the variance in the data (Supplementary Figure 21 D-F). In both sets of data, all the variants were nicely distributed along the PC1 (x-axis) according to their expression level (compare with Figure 2 A-B and Figure 3 A-D). Visual inspection of the PCA plots suggested that the variants can be clustered according to their expression levels. We performed hierarchical clustering (HC) of the complete flow cytometry dataset of the different light intensities and the complete dataset of the spectrometry data over time, and also at the different light intensities using the Ward clustering method with Euclidean distances^49^ (Figure 3 E-F). Four clusters were identified that contained variants with very high (red), high (orange), medium (green), and low (blue) expression profiles (Figure 3 E-F). The negative control (NC) not harboring the Magnets constructs was located in the low expression cluster, accordingly (data shown in Supplementary Figure 22). Visual inspection of the PC and HC cluster analyses showed that they were in agreement. Therefore, this classification of clusters was used for the analysis of the data that was gathered at different incubation temperatures during the characterization of the mutations. Despite the two datasets using different measurement methods (single-cell by flow cytometry and population by fluorometry) and time considerations (single timepoint and expression measured over time), only slight changes were observed when comparing the corresponding dendrograms at the screening temperature (37 °C) (Supplementary Figure 23). The red and orange clusters contained variants with mutations in the pMag, with the exception of M48V N100L, while the green and blue clusters contained variants with mutations both in pMag and nMag.

### nMagHigh1 variants with increased light sensitivity at 37°C

The nMagHigh1 variants L39P and V160M were located in the medium expression (green) cluster (Figure 3E) and showed an increased light sensitivity, described with a decrease in the half-maximal light intensity (*I50*=80 and 83% respectively compared to WT, Figure 2A). We also observed improved dimerization properties in these variants, visible through the increased expression at a saturating light intensity (*t*=151 and 148% compared to WT respectively, Figure 2A), while also showing increased basal expression (*b*=150 and 148% compared to WT, Figure 2A). Interestingly, a double mutant also containing a mutation at position 39 (L39R) with a second mutation T38A located in the low expression cluster (blue) showed an even further increased light sensitivity compared to the single mutant (*I50*=68% compared to the WT). Together with the reduced basal- and light-induced reporter expression (*b*=84%, *t*=77%, Figure 2A), this indicates that expression levels and light sensitivity can be tuned separately. In addition, we identified the double mutant M48V N100L, which shows a dramatically increased light sensitivity (*I50*=50% compared to the WT, light dose-response curves and expression at different intensities over time of WT and M48V N100L, Figure 2 C-D) and an increase in both basal- and light-induced expression (*b*=296%, *t* = 267% compared to the WT, Figure 2A). Accordingly, this variant clustered in the high expression group (orange).

### nMagHigh1 variants with decreased light sensitivity at 37°C

nMagHigh1 variants with decreased light sensitivity all clustered in the low expression group (blue). This includes mutations A143S and V147M (with an *150=143* and 162%, respectively compared to the WT, Figure 2E) that displayed expression levels comparable or slightly reduced compared to the WT (*b*=91 and 89%, *t*=95 and 95% respectively, Figure 2A). Also here, expression level and light sensitivity were tuned separately. In addition, double mutant I135V K137R showed an increase in the half-maximal light intensity (*150*=158% compared to the WT), a reduced dark expression (*b*=88%) and an increase in the maximal light-induced expression (*t* = 111%, Figure 2A).

### nMagHigh1 variant with shifted gene expression at 37°C

Variant K137E showed a light sensitivity that is comparable to the WT (*I50*=98%, Figure 2 F). However, the reporter gene expression was upshifted for both the dark-as well as the light-induced expression (*t*=128%, *b*=133% compared to the WT, Figure 2A). Interestingly, this variant also clustered in the low expression group.

### pMag variants with increased light sensitivity at 37°C

All pMag variants, except for positions N100, were unique (Figure 2B) compared to the ones previously described for nMagHigh1 (Figure 2A). While N100S was found in pMag, a mutation at position N100 was also found in nMagHigh1 double mutant M48V N100L. pMag variants N100L, N100K and N100S as single mutations were all identified during site saturation mutagenesis (see below). All variants R52L, T69M, T83M, N100K, N100L, N100S, K119N, M165T, S178Y, P42T T69M and P66S Y94N all showed an increased light sensitivity of up to 47% compared to the WT. (*I50=79*, 50, 47, 71, 59, 89, 78, 50, 64, 52 and 83% respectively, Supplementary Table 4). The increase in light sensitivity was most pronounced in variants T69M, T83M, N100L, M165T, and P42T T69M, all belonging to the high or very high expression clusters (*I50*=50, 47, 59, 50, and 52% respectively, Figure 2G-I, Supplementary Table 4). In addition, for all variants except for K119N (located in the low expression cluster), the expression levels were increased, which was most pronounced (higher than 300%) in variants T69M, N100L, and P42T T69M up to 540% (*t*=326, 540 and 517% respectively) and comparably less increased for the basal expression (*b*=257, 362 and 422% respectively). In contrast, variant K119N showed expression levels comparable to the WT enzyme (*b*=121%, *t*=105%) while having an increased light sensitivity (*I50*=78%, Figure 2J), again indicating that these properties can be tuned separately. Comparing both T69M variants, the single (high expression cluster, orange) and double mutant with P42T (very high expression cluster, red), light sensitivities are similar (*I50*=50 and 52%), while expression levels are upshifted in the double mutant (*b*=257 and 422%, *t*=326 and 527% respectively), indicating an even further improved dimer assembly through the additional P42T substitution.

### Site saturation mutagenesis at hotspot positions T69 and N100

Random mutagenesis can be used to identify important amino acid residues and positions. We chose the two sites T69 and N100 in the very high expression clusters (red) for further investigation through site saturation mutagenesis (SSM) as they appeared multiple times during our screening both as single and as double mutants and in addition showed highly interesting properties. T69M as single mutant as well as double mutant with P42T was one of the variants with the highest low light sensitivity in the pMag photosensor. N100 was the only residue in which variants were selected in both the nMagHigh1 as well as the pMag libraries, and in both cases increased the light sensitivity of the photosensor. As both mutations were found in the pMag photosensor, we also used this domain to construct SSM libraries either using the “22-codon trick”^50^ or as a “small intelligent”^51^ library. From these libraries, 93 randomly selected colonies were picked per 96-well plate as well as three WT controls in the same plate and incubated either in the dark or induced with low-intensity light as previously described.

SSM of position T69 was performed with a reduced “small intelligent”^51^ library with NDT, VMA and TGG for position T69, but lacking the ATG, as T69M was already identified. As shown in the inset in Supplementary Figure 24 A, T69, shown as red spheres, lies in the interface of the photoactivated VVD dimer. As a result of the SSM, only T69W was identified to perform significantly better at low light induction compared to the WT, while T69L and T69F performed similarly to the WT. However, variant T69M found during the screening outperformed T69W for mCherry expression in low as well as high light induction conditions (Supplementary Figure 24 B). T69M showed a 2.4-fold higher light sensitivity when comparing low light expression levels to the WT expression level (Supplementary Figure 24 C). However, also the sensitivity of T69W was improved 1.7-fold. For both T69M and T69W, but also for the leucine, phenylalanine and tryptophan substitution, the fold change at low light induction was improved 2.8-, 1.5-, 1.6- and 1.3-fold respectively (Supplementary Figure 24 D). In general, all mutations improving mCherry expression are amino acids with hydrophobic side chains, and therefore might play a role in the dimer formation, while substitutions that decreased the function of the photoregulator were mostly hydrophilic or polar e.g. aspartic acid, arginine, or serine (Supplementary Figure 24 ABCD).

SSM of position N100 was performed using the “22-codon trick”^50^. The position is indicated by red spheres, located at the surface of the protein close to the Ncap (Supplementary Figure 24 3E inset). During the screening, we already identified N100L in combination with M48V and N100S. N100L showed the highest expression level at low light induction (Supplementary Figure 24 E,F), which was higher than the levels obtained with the double mutant M48V N100L (Figure 2A-B). The light sensitivity of the single mutant was increased 9.9-fold for the expression at the low light intensity (Supplementary Figure 24 G), compared to 5.7-fold for the double mutation, suggesting that M48V might have a slightly detrimental effect which is overcompensated by N100L in the double mutant compared to the WT. N100L also showed an increase in the fold change at low light intensity of 3.6-fold. Further, methionine, lysine, and arginine, along with the previously identified serine showed increased light sensitivities of 3.2-, 2.9-, 2.9- and 2.5-fold respectively. The dark to light fold change was also improved in all of these variants at low light intensities with the highest improvement of 3.6-fold for N100L and slightly improved or similar to the WT at high light intensities (Supplementary Figure 24 H). The different properties of improving mutations ranging from polar via positively charged to hydrophobic side chains do not allow for clear mechanistic conclusions and require additional assessment in future studies.

### Light-induced gene expression at 30, 37, and 40°C

We performed the same functional characterization as described before (reporter expression at different light intensities) also at lower (30°C) and higher (40°C) temperatures than the standard 37°C cultivation conditions. This should aid in further investigating the performance of the Opto-T7RNAP and the discovered variants in different culture conditions as well as its effect on the mutations itself. As described for the experiments at 37°C, characterization was done by flow cytometry and over time measured through fluorescence spectrophotometry. In general, we observed an approximately 4.5-times higher maximal mCherry fluorescence (*t*=128,601 a.u.) in cells containing the original Opto-T7RNAP*(563) regulator at 30°C compared to 37°C (*t*=28,360 a.u.), which might be caused by a decrease in protein stability and/or in dimerization ability at the higher temperatures. In addition, the light intensity required for half-maximal activation decreased 3.9-fold (from *I50*=0.67 W cm^-2^ at 37°C to *I50*=0.17 W cm^-2^ at 30°C). Therefore, experiments were performed at a lower light intensity range for experiments performed at 30°C (from 0 to 0.96 W cm^-2^) compared to the experiments at 37°C (from 0 to 5.77 W cm^-2^). Accordingly, similar effects were observed for experiments performed at 40°C. At this temperature, the highest used light intensities (5.77 W cm^-2^) did not lead to a saturating reporter expression. To avoid additional heating of the samples caused by the light induction, we did not further increase the light intensities. As increased light intensities would be necessary to reach saturation, which would be required to mathematically fit the dose-response curve, we directly compare the fluorescent signal of the reporter expression in the dark and in the light intensity that led to saturation at 37°C (3.85 W cm^-2^). In comparison to the reporter expression at 37°C, the higher temperature (40°C) led to a 114-fold reduction in reporter expression with 3.85 W cm^-2^ light induction (28,854 a.u. and 252 a.u. respectively), while dark expression was reduced around 6.4-fold (517 a.u. and 82 a.u. respectively). To summarize, this pre-characterization of the wild-type Opto-T7RNAP system showed that light-induced reporter expression and light sensitivity was highest at 30°C and decreased as the temperature was increased to 37°C and 40°C. As described above at 37°C, analogous PC analysis showed that all the variants were nicely distributed along the PC1 (x-axis) according to their expression level also at 30 and 40°C (Supplementary Figure 21 A,C,D,F). Analysis of the number of clusters using the silhouette algorithm^52^ and the data at all temperatures suggested an optimal number of clusters of 4-5 or 4, depending if the dataset was single-cell or population. Four clusters were selected and visual inspection of the HC analyses suggested that they were in agreement.

### Variant characterization at 30°C

As observed for the WT, all variants showed an increased reporter expression at the lower expression temperature (Figure 3 A-B). Interestingly, due to the overall higher expression at 30°C (Figure 3 AB), the NC was the only variant in the low expression cluster, while at 37 and 40 °C the low expression cluster contained 6-7 and 8-11 variants, respectively. From the single-cell expression data, all the members of the medium expression cluster, except V160M, L39P, N100S, and N100K moved to the low expression cluster at 37°C and 40°C. In most cases, the general trends observed for the different variants (Supplementary Table 6) are comparable to 37°C (Supplementary Table 2). Interestingly, nMagHigh1 variant A143S that showed a reduced light sensitivity at 37°C (*I50*=143% compared to the WT) displayed a similar sensitivity as the WT at 30°C (*I50*=98%). Additionally, pMag variant R52L showed a dramatically increased expression level at 30°C, which was similar to pMag double mutant P42T T69M (*t*=206 and 207% respectively compared to the WT) in addition to the increased light sensitivity (*I50*=63 and 56% respectively). In comparison, at 37°C, the expression of R52L was still increased compared to the WT (*t*=275%) but significantly lower than P42T T69M (*t*=517%) or N100L (*t*=540%). Accordingly, variant R52L moved from the very high expression (red) cluster at 30 °C to the high (orange) at 37 °C and medium expression (green) cluster at 40 °C in both datasets, while N100L moved from the high at 30°C to the very high expression cluster at 37°C (Figure 3 E-F).

### Asymmetric changes for dark and light-induced expression

nMagHigh1 variants V147M and I135V K137R both led to a more pronounced decrease in the basal expression at 30°C (*b*=47 and 41% respectively) compared to the expression at 37°C (*b*=89 and 88%), whereas the relative maximal expression was similar in both conditions (*t*=97 and 106% at 30°C and *t*=95 and 111% at 37°C). The contrary was observed for nMagHigh1 variant M48V N100L, where light-induced expression was similar and the basal expression increased compared to the WT at 30°C (*t*=109%, *b*=271%), but increased for both conditions at 37°C (*t*=267%, *b*=296%) whereas the dramatic decrease in light sensitivity was comparable for both temperature conditions (*I50*=50% at 37°C; *I50*=52% at 30°C). For this variant, the extremely low light intensity of 0.09 W cm^-2^ led to half-maximal activation of gene expression. Similarly, pMag variants T69M, T83M, N100K, N100L, and P42T T69M showed a lower increase in the maximal expression at 30°C compared to 37°C (*t*=132, 125, 88, 165 and 207%at 30°C; *t*=326, 270, 190, 540 and 517% at 37°C, respectively), while the change in the basal expression at both temperatures was similar or comparably less (*b*=260, 245, 184, 253 and 468% at 30°C; *b*=257, 232, 145, 362 and 422% at 37°C). For all pMag variants, an increased light sensitivity was observed at 30 as well as 37°C (Supplementary Table 8, Supplementary Table 4).

### Variant characterization at 40°C

Similar to the WT, expression of all variants was dramatically reduced at 40°C compared to 37°C (Figure 3 C-D, Supplementary Tables 10-12). However, all variants with increased light sensitivity and increased reporter expression at 37°C, also showed an increased reporter expression compared to the WT in all light conditions, which might suggest an improved thermostability. The largest increase in expression compared to the WT was observed with nMagHigh1 variant M48V N100L, and pMag variants T69M, N100L, P42T T69M which showed an increase in reporter expression of 412, 407, 782, and 781% respectively at the highest light intensity (Supplementary Tables 10-12), but only variant N100L moved from the high at 37°C to the very high expression group at 40°C to cluster together with P42T T69M. Variant nMagHigh1 T38A L39R, which displayed increased light sensitivity at both 30 and 37°C (*I50*=75% at 30°C; *I50*=68% at 37°C), showed decreased expression levels at all temperatures (compare Supplementary Table 2 and Supplementary Table 10).

### Population data over time supports single-cell data at one single time point

Since the results obtained with the clustered spectrophotrometry data over time was highly similar to the single-cell data as described above and confirmed that the behavior of the variants is not restricted to one single time point at low cell density, but rather consistent up to the early stationary phase of growth. To further analyze the expression at higher cell density, we fitted the fluorescent data to light dose-response curves at one specific time point at which the variants reached the highest fluorescence levels. While in the single-cell analysis, cells were in the exponential growth phase at low cell density, the time point with the highest fluorescence in the population data was reached at the end of the exponential phase - beginning of the stationary phase. Despite this difference in growth stages, results were comparable. For example, nMagHigh1 variants L39P, T38A L39R, and V160M showed increased light sensitivity through a decrease in the half-maximal light intensity at 37°C (*I50=70*, 70, 52% respectively compared to the WT; Supplementary Table 3). Also in these experiments, nMagHigh1 K137E showed a light sensitivity that is comparable to the WT (*I50*=102%). As observed in the single cell data, also here all pMag variants revealed an increased light sensitivity (*I50*=79, 60, 55, 88, 71, 88, 78, 63, 58, 55 and 83% compared to the WT for R52L, T69M, T83M, N100K, N100L, N100S, K119N, M165T, S178Y, P42T T69M and P66S Y94N respectively; Supplementary Table 5). The previously described reduced light sensitivity at 37°C and similar sensitivity as the WT at 30°C for nMagHigh1 variant A143S was also observed in the population dataset (*I50*=160% at 37°C, *I50*=117% at 30°C; Supplementary Table 3, Supplementary Table 7). Also the asymmetric changes for dark and light-induced expression of nMagHigh1 variants V147M and I135V K137R, which led to a more pronounced decrease in the basal expression at 30°C compared to the expression at 37°C and more similar maximal expression in both conditions (see discussion above) could be observed in the population dataset (V147M: *b*=103%, *t*=86%, *I50*=167% at 37°C and *b*=47%, *t*=99%, *I50*=143% at 30°C; I135V K137R: *b*=111%, *t*=82%, *I50*=178% at 37°C and *b*=51%, *t*=98%, *I50*=124% at 30°C; Supplementary Table 3, Supplementary Table 7).

Altogether, the hierarchical clustering, as well as the comparison of parameters, show that the mutations led to similar behaviors both for expression measured at a discrete time point during the early logarithmic growth phase as well as for gene expression over time. This is an important characteristic for future applications of this optogenetic system.

### Variants for enhanced gene expression with mOptoT7 in mammalian cells

After characterization of the different variants in *E. coli*, we set out to test selected mutations also in mammalian cells. For this, we chose variants that showed increased light sensitivity and boosted gene expression. These variants should address two major challenges of optogenetics using the mOptoT7.^53^ First, optogenetic systems with increased light sensitivity require lower intensity light for induction, thus reducing potential phototoxic effects^45^ especially during long-term experiments. Second, although the mOptoT7 has the unique advantage of being a separate transcription system, the overall expression level of the original version is limited. Thus, elevating the light-induced expression level widens the applicability of the mOptoT7.

For this, we incorporated a reduced set of mutations (Figure 4) into version 1 of the mOptoT7 and used it for light-induced expression of mRuby3 from a separate reporter plasmid. This plasmid contains an IRES2 sequence and a polyA tail for enhanced expression in mammalian cells, as described previously.^53^ A constitutively expressed mCitrine plasmid was added as transfection control. HEK293T cells were transfected as described in the Methods section and measured through flow cytometry after 24h of constant light induction at a previously characterized sub-saturating light induction condition (0.01 mW cm^-2^) or incubation in the dark. We observed an increased expression level at the lower light intensity with nMagHigh1 variants L39P, N100L, and N100L M48V compared to the WT. While WT expression only increased 2.7-fold in this condition, L39P, N100L, and N100L M48V reached approximately 6.7- 7,4- and 10,7-fold higher expression than the dark basal expression of the WT. It has to be noted, that also the dark state of these highly expressed variants was increased correspondingly (Figure 4). Thus, the mutations tested here might be especially useful if low-intensity light has to be applied due to phototoxicity/light-application limitations, or if a generally high expression of a gene of interest is required.

**Figure 4:**
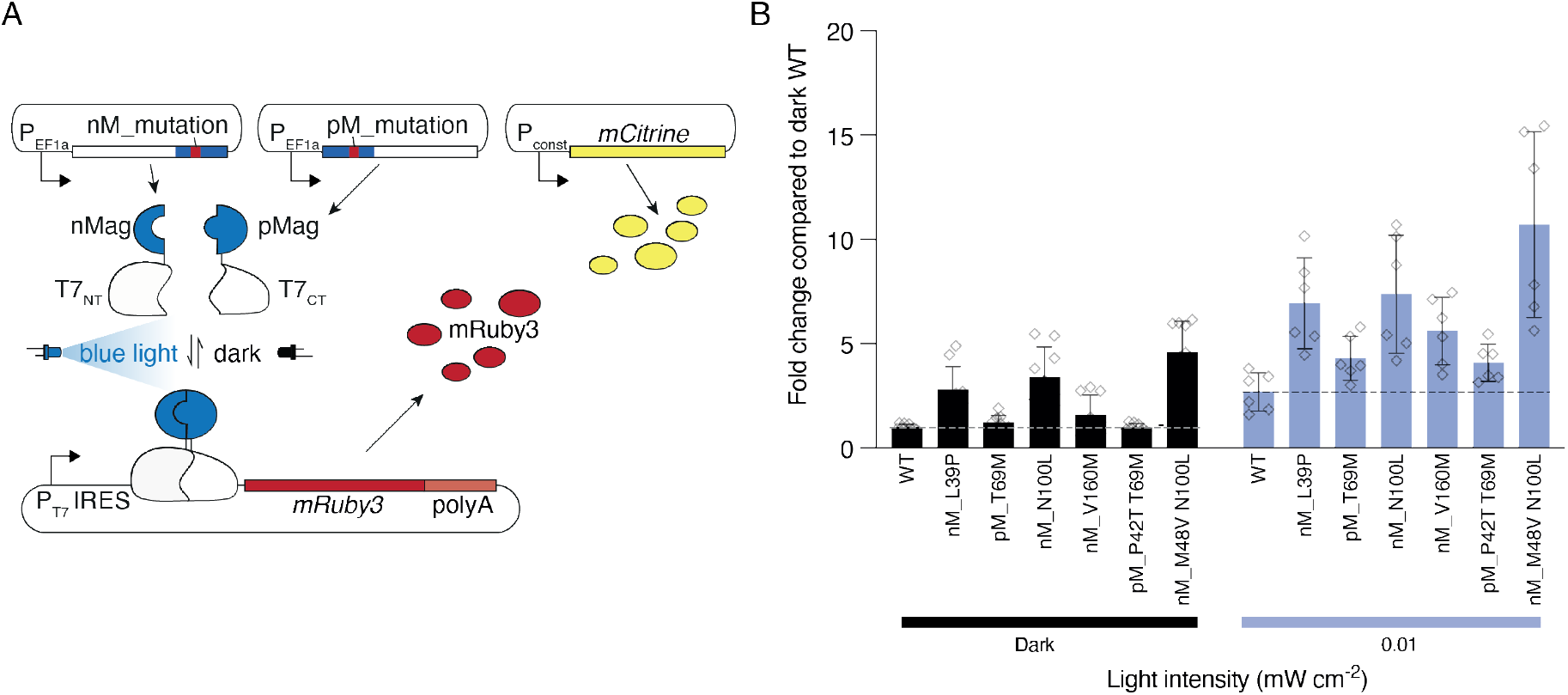
Characterization of selected Magnet variants in mammalian cells. (A) Plasmids used for transfection of HEK293T cells. The mOptoT7 fragments are expressed from EF1a promoters and a separate plasmid for each of the components. The mutations were either located in nMagHigh1 (nM_mutation) or pMag (pM_mutation). mOptoT7 drives the transcription of mRuby3. mCitrine is used to normalize the mRuby3 expression to the transfection efficiency. (B) Fold change of normalized mRuby3 expression of the dark wild type mOptoT7 (WT) to both the light-induced WT and the dark and light-induced expression selected identified variants at a sub-saturating light-intensity (0.01 mW/cm^2^; dose-response curve in Dionisi et al. Figure 1 ^53^). Shown are normalized fold change values as mean with standard deviation as well as individual points of at least six biological replicates (n=6).

## Discussion

### Features of identified mutations

In this work, we have identified a set of 14 single and 5 double amino acid substitutions in the light-inducible Magnets domains after a random mutagenesis screen. All mutations show changes in their light sensitivity and/or their expression levels. Interestingly, we found that properties such as light sensitivity and expression level could be changed independently. For example, pMag N100L shows both increased reporter expression and light sensitivity compared to the WT, while for nMagHigh1 T38A L39R, which was also more light-sensitive, we observed a decreased reporter expression. Variant K137E in nMagHigh1 in turn showed an upshifted dark-as well as the light-induced expression, while maintaining a similar light sensitivity compared to the WT. Increased light sensitivity variants thus require lower light intensity for full induction, which is beneficial when light toxicity or light penetration into a sample is problematic (e.g., in large culture volumes). The increased overall expression level of some mutants is especially interesting for systems in which the setpoint for dark- and light-induced expression cannot easily be altered. Through the use of these mutations, the setpoints can be genetically fixed at different levels.

While the goal of this study was to demonstrate an approach for easy photoreceptor tunability, it would be of interest for further studies to test combinations of mutations, especially if certain variant properties are to be combined. Although mutations might not behave additively, we found several identical amino acid substitutions both in single- as well as in double-mutant variants. An example is double mutant P42T T69M, which shows a similar *I50* to the single mutation T69M, but an increased expression level, suggesting that such a combinatorial screening for additive effects of these mutations might further enhance certain properties.

To better understand how it is possible that expression level or light sensitivity could be so easily changed, the context of the native protein origin must be taken into consideration. The VVD photodimerization system originates from *Neurospora crassa*, where it is involved in the circadian clock. It was previously reported that the VVD protein is temperature regulated, degrading faster at higher temperatures to ensure a stable clock in a wide range of physiological temperatures.^54^ Since the Opto-T7RNAP expression system is mainly used at 37°C in both bacteria and mammalian cell cultures, we performed our screening at this temperature. Thus, improved thermostability might be one feature that was optimized for along with improved dimer assembly or light-mediated allosteric changes within the protein. Regarding thermostability, different phenomena could be observed. For example, pMag variant R52L showed the most pronounced expression level increase at 30°C, where it was the highest expressing variant together with double mutant P42T T69M (Figure 3B). At 37°C its expression level was still 275% higher than the WT, however, several other variants outperformed it (T69M, N100L, P42T T69M with *t*=326, 540, and 527% respectively). The same variants also showed a higher expression than pMag R52L at 40°C (Figure 3D). A similar trend was seen with variant P66S Y94N, which expressed well at 30°C, and the improvement of gene expression in comparison to the WT decreased at 37°C and was comparable to the WT at 40°C (Figure 3E-F), indicating that thermostability might not be the major contribution of the mutation for the changes in properties. The opposite was observed for pMag N100L, which was outperformed by R52L and P42T T69M and was on par with P66S Y94N at 30°C. At 37 and 40°C however, N100L was the highest expression variant, indicating an improvement in temperature stability. In contrast to these increasing or decreasing expression levels with temperature, pMag variant P42T T69M was consistently one of the highest expressing variants at all temperatures, which might indicate improved temperature stability as well as dimerization properties. Comparison with the findings of Vaidya et al.^40^ shows that the same mutation, in this case T69M, could have different effects depending on if the VVD photoregulator is used for homodimerization or through Magnets mutations as a heterodimerization system. Although it leads to constitutive dimerization in the dark and light in the homodimerization system^40^, it led to improved light sensitivity and dimer assembly in our study.

### Structural comparison of mutations

Our goal was to develop a strategy that enables quick and easy parameter tuning of the photodimerization domain. By not restricting the engineering to specific functional sites of the protein, such as the FAD-binding pocket or the dimer interface, we obtained mutations in different compartments of the domain. In general, the VVD photoregulator comprises a LOV Per-Arnt-Sim domain with an N-terminal cap region (Ncap; residues 37-70) and a loop that accommodates the flavin cofactor.^26^ Light induces a covalent cysteine-flavin adduct between the C108 cysteine thiol and the flavin C4a position and induces conformational changes that propagate to the N-terminal helix and the protein surface, which leads to the release of the N-terminal part from the protein core, forming a symmetric dimer via hydrophobic amino acids. Return to the dark state is caused by the scission of this thioether bond. The flavin ring is bound in a pocket formed by two helices (Ea and Fa) and three strands of the central betasheet (Aβ, Hβ, and Iβ) at the end of a water channel, and the structure contains an 11-residue FAD loop at the surface of the protein.^26^

The single mutations we found are located in all functional parts of the two photosensory domains (shown as red spheres in Supplementary Figure 25 A). For visualization purposes, we used the solved structure of the light-induced VVD dimer (PDB code: 3RH8). From the Ncap mutations, T38A, L39P, P42T are located in the N-terminal “latch” (amino acid residues 37 - 44) that wraps around the domain (Supplementary Figure 25 C). Mutations M48V and R52L are located in the interface of the dimer within the subsequent Aa helix, and P66S and T69M are located in the hinge region of the dimer interface. Several single mutations (T38A, L39P, R52L, T69M, and T83M) all show hydrophobic properties, which might aid in dimer formation. Interestingly, T69W was previously described as facilitating intersubunit contact.^40^ In this study, T69W caused VVD homodimerization in both the dark and the light state. With variant T69M, we still observed light inducibility, resulting in even higher mCherry expression levels than the WT. M165T, S178Y, T83M, and M179V are located at the flavin harboring water channel (Supplementary Figure 25 D-E). Specifically, M165T, S178Y and T83M are in proximity to the flavin isoalloxazine ring (<3.8 Å for the original residues in VVD), which might aid in positioning of the chromophore for formation of the cysteine-flavin adduct and subsequent structural rearrangements. T83M at the entrance of the water channel harboring the flavin chromophore and K119N in the FAD loop (Supplementary Figure 25 E) were both characterized as single mutations, and might alter chromophore binding and positioning, again allowing for cysteine-flavin adduct formation at lower light intensities. To gain more definite answers, x-ray crystallography and UV-vis spectroscopy^55^ could shed more light on the underlying structural and mechanistic functions of these amino acid exchanges. This is out of the scope of our engineering-driven approach and should be investigated in future studies.

Zhou et al identified five features that better define the differences between the active, inactive, and transition states. Of those five features, the most important ones were the distances T38-G105 and T38-K119.^56^ Interestingly, we found mutations in two of those residues, both reducing the length of the side chain (T-to-A in 38 and K-to-N in 119), for which we observed reduced light sensitivity.

### Comparison to other VVD mutations

Our findings also revealed interesting connections and additions to previously known mutations. An intriguing example was identified with variant pMag R52L. Previously, VVD variant I52C was reported as having increased homodimer-forming efficiency in both the dark and light state^19^. This position was then used to transform the homodimeric VVD protein into the heterodimeric nMag/pMag Magnets system, in which I52R together with M55R were used for the “positively charged” pMag.^25^ Our mutagenesis revealed that, in combination with nMagHigh1, pMag variants in which the positively charged arginine is changed to leucine, and thus back to an amino acid with a similarly hydrophobic sidechain as with the initial isoleucine, increase both expression and light sensitivity. In our characterization, this change increased both the light-induced and basal expression as well as the light sensitivity at all tested conditions (Supplementary Table 4, Supplementary Table 5, Supplementary Table 8, Supplementary Table 9, Supplementary Table 11).

Residues M135 and M165 are in contact with the flavin ring. Substitution of those residues with isoleucine resulted in variants that remained in the ‘on’ state tenfold longer^41^, slowing the photocycle^57^ of *N. crassa*. During the Magnets development, these mutations M135I M165I were identified to further increase dimerization efficiency. The Opto-T7RNAP version used for screening contained these mutations in the nMag (nMagHigh1), however, we did not include them in the pMag domain, as the combination of nMagHigh1 and pMagHigh1 shows increased dimerization in the dark.^25^ During screening for increased expression level we identified a variant that contained both mutations, and thus it was identical to pMagHigh1.

Another interesting finding was T69M, which appeared both as a single mutation and double mutation together with P42T in pMag. Vaidya et al. found that in VVD, T69W forms dimers in the presence and absence of light. They attributed this to facilitated inter-subunit contacts that overcame light-promoted conformational switching, as previously described. In the Magnets, T69L was found to improve hydrophobic interactions.^44^ T69L, together with S99N, M179I, were identified from comparison with VVD domains from thermophilic fungi.^44^ It was hypothesized that T69L improves hydrophobic interactions, M179I in the hydrophobic core improves packing, and surface-exposed S99N optimized hydrogen bonding and secondary-structure preference. Our site saturation experiments on position T69 identified T69M as the variant which showed the highest expression levels both with high and low light induction, although the expression of T69W was also increased in both cases (Supplementary Figure 24 B). Thus, while T69W might lead to light-independent binding in the homodimeric VVD protein, our findings suggest that the heterodimeric interactions in the Magnets allow for the change of one component, which increases both the expression level as well as light sensitivity.

Our characterization further revealed that multiple different side-chain exchanges of N100 led to both increased expression levels and light sensitivities, with N100L showing the highest changes. N100R was previously reported to allow for stronger dimerization at 37°C due to improved helical preference, which was also rationally identified due to comparison with VVD variants from thermophiles.^44^ In our study, N100L led to the highest expression at both 37 and 40°C through improved thermostability. Our site saturation mutagenesis confirmed that the previously identified N100R variant also shows increased expression compared to the wild type, however it was outperformed by N100L (Supplementary Figure 24).

## Conclusions

Overall, our results show that the Opto-T7RNAP transcription system creates an excellent phenotype-genotype linkage that can be used to adapt various properties of this and possibly other heterodimerization systems towards their actual application in research or biotechnological processes. We have found previously described mutations/residue positions and identified new ones together with hotspots. We challenged this approach by screening for variants with improvements in the most fundamental feature of photoregulators, their light sensitivity. After just one round of mutagenesis, variants with mutations within all the important structural compartments of the photoregulator, the Ncap, the PAS core, as well as the FAD-loop were identified. The mutations which lead to increased light sensitivity were distributed all over the photosensory domain, highlighting the benefit of random mutagenesis of the full photosensor compared to rational selection of residues at important sites. We also showed that selected mutations can also be used in mammalian cells to overcome some of the limitations encountered when using blue light-based optogenetic tools: light toxicity and limited gene expression level. This approach can also be used to screen for other properties such as lower dark-state assembly, improved light-induced dimerization at saturating light induction, increased dark to light fold change, or even dynamic features such as increased or decreased dark-state dissociation rates by varying the light input before sorting. A crucial requirement for successful screening campaigns, especially for features such as induction at sub-saturating light intensities, is the relatively high fold-change of the Opto-T7RNAP. Thus, mCherry expression levels of cells containing the original Opto-T7RNAP*(563) were still sufficiently different from dark controls to screen with sub-saturating light induction, which allowed us to distinguish desired variants from mutations that inactivated the function of the regulator. This also abolished the need for multiple enrichment steps for the different induction conditions, which lead to a broad set of variants. While we screened for changes in the light sensitivity and in expression levels of the photodimerization domains, the same approach can be undertaken to screen for other properties. This is essential for perfectly adjusting photosensors to specific applications. In addition, the variants we identified for the Magnets domains which are based on the VVD photoregulator, will find immediate use in the many existing designs that utilize this photosensor and will significantly expand the applicability of these domains and the optogenetic toolbox.

## Materials and Methods

### Bacterial strains and media

*E. coli* Top10 was used for all cloning. For characterization, we used *E. coli* strain AB360^7^. The strain contains the transcription factor AraC, whereas arabinose metabolizing genes *araBAD* are deleted, and *lacYA177C* which allows for titratable arabinose regulation. Plasmids were transformed using a one-step preparation protocol of competent *E. coli* for transformation of plasmids in testing strains^58^ or made electrocompetent for subsequent electroporation^59^. *E. coli* CloneCatcher™ DH5G Gold Electrocompetent was used for high efficiency transformation of mutagenesis libraries following the manufacturer provided protocol (Genlantis, Inc.).

Autoclaved LB-Miller medium was used for strain propagation. Sterile-filtered M9 medium (M9 Minimal Salts 5X, Sigma Aldrich) supplemented with 0.2% casamino acids, 0.4% glucose, 0.001% thiamine, 0.00006% ferric citrate, 0.1 mM calcium chloride, 1 mM magnesium sulfate was used for all gene expression experiments. Antibiotics (Sigma-Aldrich Chemie GmbH) were used as necessary for plasmid maintenance at concentrations of 100 μg/mL ampicillin, 17 μg/mL chloramphenicol, and 50 μg/mL kanamycin.

### Plasmids and genetic parts

Plasmid pAB150 containing the Opto-T7RNAP*(563) as well as the reporter plasmid pAB50 with mCherry under T7 promoter expression control are taken from our previous study.^7^ All primers used are listed in Supplementary Table 13. The plasmid with lower TIR was constructed by using the RBS Calculator V.2.0^47^ and inserted via PCR amplification of mCherry with primers oAB819 and oAB507 from template pAB50, digested with BamHI and KpnI and ligated into the BamHI and KpnI digested backbone. Primer and plasmid sequences are described in Supplementary Table 13.

Error-prone PCR of Magnets domains was performed through amplification of nMagHigh1 and pMag from pAB150 with primers pairs oAB734/oAB736 and oAB810/oAB744 respectively and error-prone Pfu DNA Polymerase containing D141A, E143A mutations^60^, a gift from Dr. Luzius Pestallozi (DBSSE, ETH Zurich). We used template concentrations of 2.5, 5, 10 ng to reach variable mutation rates. As backbone we used pAB150 amplified with Phusion HF polymerase (Phusion flash HF PCR master mix, Thermo Scientific) and primer pairs oAB589/oAB446 and oAB448/oAB809 for nMagHigh1 and pMag insertion, respectively, and for Gibson assembly^61^. The mutation rate was estimated at 1.3-2.9 per kilobasepair through Sanger sequencing. All identified mutations were recloned into the original pAB150 plasmid and used for final characterization. For nMagHigh, the inserts were amplified using primers oAB409/oAB696 and the backbone amplified from pAB150 with primers oAB589 /oAB446. Before gel purification, the PCR product was digested with DpnI followed by restriction digestion with AvrII and BglII, gel extraction, and ligation. For pMag, inserts were amplified with primer pair oAB49/oAB285 and the backbone was amplified with primers oAB448/oAB286. Before gel purification, the PCR product was digested with DpnI followed by restriction digestion with PacI and KpnI-HF, gel extraction, and ligation.

Single point mutations in all mOptoT7 plasmids were based on previously published plasmids (mOptoT7 Version 1, V1)^53^ and created using CloneAmp HiFi PCR Premix (Takara Bio). All constructs were transformed into Top10 E. coli competent cells and checked through sequencing (Microsynth).

### Screening and isolation of variants

We picked the colonies obtained from the sorting into 96-well plates containing M9 medium and grew them overnight to full cell density. We again inoculated main cultures of the individual variants at low cell concentrations so that the cultures are in log growth phase throughout the experiment for single-cell measurements, before inhibition of transcription and translation for maturation of mCherry as previously described^7^. From every plate, we then inoculated two 96-well plates, and grew them for 5h either in the dark or in the light before inhibition and measurement through flow cytometry. Based on these first characterizations, we chose a reduced subset of variants with differing mCherry expression levels as well as fold changes higher than the Opto-T7RNAP*(563) “wild-type” (WT) control. These variants were then recharacterized and again a subset of these variants was sent for sequencing. We then re-cloned mutations of unique Magnets variants identified through sequencing into the original Opto-T7RNAP*(563) plasmid. We transformed the recloned variants into AB360 strains contains the pAB50 reporter plasmid and grew them in 96-well plates overnight to full cell density, and then stored the plates in 25% glycerol at −80 C. For final characterizations, we inoculated main cultures in 96-well plates at low cell concentrations through pin replication (96 pin replicator, Scinomix, MO, USA, SCI-4010-0S) so that the cultures are in log growth phase throughout the experiment before inhibition of transcription and translation for maturation of mCherry through flow cytometry (Figure 1D, left; for WT).

### Growth and light induction conditions

All experiments were performed in an environmental shaker. The shaking incubator consisted of a Kuhner ES-X shaking module (Adolf Kühner AG, Basel, Switzerland) mounted inside an aluminum housing (Tecan, Maennedorf, Switzerland) and temperature controlled using an “Icecube” (Life Imaging Services, Basel, Switzerland) at 37°C with shaking at 300 rpm and black, clear bottom 96-well plates (Cell Culture Microplates 96 Well μClear^®^ CELLSTAR^®^, Greiner Bio-One GmbH, Product #: 655090), which was sealed with peelable foil (Sealing foil, clear peelable for PlateLoc, No. 16985-001, Agilent) to eliminate liquid evaporation and guarantee sterility, as well as a plastic lid (Greiner Bio-One GmbH, Product #: 656171). The light induction setup was previously described.^30^ For experiments, cultures were pin replicated into fresh M9 medium containing the respective inducer concentrations as described in the methods section for the screening and isolation of variants. This high dilution ensures that the cells are still in logarithmic growth phase after 5 h, at the end of the experiment, as previously described^7^. 200 μl of inoculated culture was incubated per well of the 96-well plates. Cells were grown for 5h before transcription and translation was stopped with rifampicin and tetracycline as previously described.^7^

### Flow cytometry measurement

Cell fluorescence was characterized using a CytoFlex S flow cytometer (Beckman Coulter) equipped with CytExpert 2.1.092 software. mCherry fluorescence was measured with a 561 nm laser and 610/20 nm band pass filter and the following gain settings: forward scatter 100, side scatter 100, mCherry gain 300. Thresholds of 2,500 FSC-H and 1,000 SSC-H were used for all samples. The flow cytometer was calibrated before each experiment with QC beads (CytoFLEX Daily QC Fluorospheres, Beckman Coulter) to ensure comparable fluorescence values across experiments from different days. For the primary high-throughput characterization of the sorted variants, measurements were aborted after 5,000 events or after a period of 30 seconds. At least 15,000 events were recorded in a two-dimensional forward and side scatter gate, which was drawn by eye and corresponded to the experimentally determined size of the testing strain at logarithmic growth and was kept constant for analysis of all experiments and used for calculations of the median and CV using the CytExpert software (Supplementary Figure 26). Cells were inhibited with rifampicin and tetracycline and mCherry was matured as previously described before measurement.^7^

### Spectrophotometric and fluorometric measurements

Cells were grown overnight in a 96-well master plate containing 200 μl of M9 liquid media and the appropriate antibiotics in the dark. This master plate was pin-replicated into 96-well black, clear bottom 96-well plates (Cell Culture Microplates 96 Well μClear^®^ CELLSTAR^®^, Greiner Bio-One GmbH, Product #: 655090), containing 200 μl of fresh M9 liquid media plus antibiotics and sealed with peelable foil (Sealing foil, clear peelable for PlateLoc, No. 16985-001, Agilent). The 96-well plates contained each strain in technical triplicates. Measurements were performed as previously described^62^ with the following changes. Measurements in a Tecan Infinite 200Pro and Firmware v. 3.40 were multiplexed for absorbance at 600 nm (to measure bacterial growth) and fluorescence at 535 nm (ex) / 595 nm (em) to quantify mCherry expression. The reader settings were adjusted to 9 nm bandwidth, 25 flashes and 200 ms settle time for absorbance mode and to 25 nm bandwidth for excitation, 35 nm bandwidth for emission, gain 100, 25 flashes, integration time 2,000 μs, and 200 ms settle time. Plates were incubated in an environmental shaker as described above. Light induction was performed using the 96-LED array described above at intensities ranging from 0 (dark control) to 0.96 W cm-2 at 30 °C and from 0 (dark control) to 5.77 W cm-2 at 37 °C and 40 °C. Measurements were taken at 1 h time intervals with the aid of a robotic arm.^62^. At 30 and 37 °C, we observed overgrowth of the bacterial cultures in some of the wells that were accompanied by higher fluorescence levels out of the range observed within the replicates and in comparison to other variants. These samples were excluded from the analysis using the criterion: absorbance at 600 nm > 0.805. This excluded an average of 31 and 15 samples per experiment, coming up to 5.5% and 2.6% of the total number of samples at 30°C and 37°C, respectively. Background subtraction was only done at 40 °C because at lower temperatures the background levels (fluorescent levels of the negative control not expressing mCherry) did not significantly affect the analysis of the data. At 40 °C background subtraction was performed individually with the background level at the corresponding temperature.

### Fluorescence-Activated Cell Sorting (FACS)

10 mL M9 was inoculated with 50μL of glycerol stocks of the libraries and grown in dark tubes overnight at 37°C and 230 rpm. AB363 controls were prepared accordingly. From these overnight cultures, 2 μL was used for inoculation of 10 mL of M9 (1:5000 v:v), mixed through vortexing and distributed to single wells of a 96-well plate which were immediately incubated and exposed to the respective light conditions for 4 hours at 37°C. The respective libraries or controls were then again pooled and pelleted at 4000 rpm for 15 min and 4°C. The supernatant was discarded and the pellet resuspended in ice-cold PBS. The samples were kept on ice in dark tubes until they were sorted. It was defined before that mCherry fluorescence values of cells in PBS at 4°C remain constant (data not shown). Single-cell sorting was performed using a Sony SH800S Cell Sorter and a 70 μm chip. Fluorescence of mCherry was measured with a 561 nm laser and a 617/30 nm bandpass filter. The sorting chamber was maintained at 4°C and sample chamber pressure was never set higher than 4 arbitrary units on the machine. Gain settings were set individually at the respective control samples. Sheath fluid was used as running buffer, and gates for setting the threshold for sorting were individually set. Their stringency varied and was determined by comparison to the respective control sample. For enrichment, cells were sorted into 8 ml LB medium in black tubes (Black Tubes CELLSTAR^®^ (15 mL), Greiner Bio-One GmbH), while for singlecell sorting, selected events were spotted onto LB agar omnitrays (Nunc™ OmniTray™, Thermo Scientific) supplemented with chloramphenicol and ampicillin. Plates were grown at 37°C overnight. Colonies were picked in 96-well plates (Microplates 96 well polystyrene, Product #: 655161, Greiner Bio-One GmbH) filled with 200 uL LB medium using sterile tooth picks and incubated overnight at 37°C and 230 rpm and sealed with gas-permeable membrane (BREATHseal™, Greiner Bio-One GmbH). The next day, 100 μL was pipetted into a new 96-well plate and 50% glycerol (PanReac Applichem) was added into both plates which were then sealed and frozen to −80°C until use.

### Fitting of light dose-response curves

For fitting of mCherry expression in response to light intensity using least squares regression, we used Prism 9 for MacOS (Version 9.3.1 (#350, GraphPad Software, LLC.).

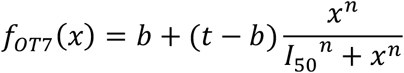

where *f_OT7_*(*x*) describes the gene expression controlled by the respective Opto-T7RNAP variant as a function of light intensity, *x* represents light intensity, *b* corresponds to the basal promoter under not activated conditions, *t* is the maximal promoter expression, *I*_50_ is the light intensity for half-maximal activation, and *n* is the Hill coefficient for Opto-T7RNAP. As weighting method, no weighting was chosen, and each replicate Y value considered as an individual point.

Values for *b, t, I*_50_, *n*, and their lower and upper 95% profile-likelihood confidence limits as well as R squared values, were calculated and confidence bands shown in the respective plots.

Flow cytometry measurements were performed after 5h induction during the exponential growth phase. As the time point for the spectrophotometric measurement, we chose 8h for cells grown at 37°C and 13h for cells grown at 30°C. To compensate for the slight shift in OD600 of variant N100L 30°C, and thus slightly later fluorescence peak, we took the 15h time point only for this variant.

### Cell culture and Transfection

HEK293T cells (ATCC, strain number CRL-3216) were cultured and transfected as described previously^53^ with the following modifications: transfections were carried out in suspension using 24 well plates (either black for light experiments, PerkinElmer, or transparent for dark control, ThermoFisher) at a density of 1.5×10^5^ cells/well; DNA and PEI complexes were incubated for 25 minutes at room temperature prior to addition to the cells.

### Light induction and Flow Cytometry analysis of HEK293T cells

Cells were illuminated with constant light using 470nm LEDs (Super Bright LEDs Inc) in an optimized version of the Light Plate Apparatus (LPA).^53^ HEK293T cells were analyzed 24h after illumination using CytoFLEX S Flow Cytometer (Beckman Coulter) as described previously.^53^ For each sample, light-induced expression was normalized to mCitrine constitutive plasmid (dividing the mRuby3 reporter fluorescence values by the mCitrine fluorescence values) to account for transfection effiency. Data was analyzed using Cytoflow Software and a customized R code.

### Clustering analysis

Principal component analysis and hierarchical clustering was performed in R using the following packages: cluster^63^, factoextra^64^, dendextend^65^ and ggplot2^66^. The optimal number of clusters was determined using the silhouette algorithm^52^ implemented in the factoextra package of R.

The flow cytometry (single cell) data was scaled before PC analysis. We observed that scaling the data before PC analysis for the fluorometric (population) data gave too much weight to the measurements in the lag phase of growth when the mCherry activity was below the detection level of the plate reader due to the low number of cells. Therefore, the fluorometric data was not scaled before PC analysis.

Hierarchical clustering was performed using the Ward method with Euclidean distances.^49^ When we analysed all the data at the three different temperatures together (in both cases: single cell-flow cytometry and population-fluorometric data), we employed the principal components scores to perform the hierarchical clustering rather than the raw data to have the same mean in the data at all temperatures and again to avoid giving special weight to the data at 30 °C, where the mCherry showed the highest values.

### Protein structure visualization

Protein structures were visualized using the PyMOL Molecular Graphics System, Version 2.5.2 Schrödinger, LLC.

## Author Contributions

A.B. conceived, planned, and coordinated the project and wrote the manuscript with contributions from all authors. A.B. and Y.W. generated the libraries and performed the FACS. A.B., Y.W., and D.C. designed and performed bacterial experiments and analyzed the corresponding data. D.C. performed the PC analysis and hierarchical clustering. S.D. performed experiments in mammalian cells and analyzed the corresponding data. M.K. supervised the project and provided funding.

## Acknowledgements

We thank Dr. Tsvetan Kardashliev for helpful discussions and Dr. Luzius Pestalozzi for the testing and the supply of the polymerase and buffer used for error-prone PCR. We further thank Dr. Stephanie Aoki for helpful discussions. We thank the Single Cell and Lab Automation Facility of the DBSSE, ETH Zurich, in particular Dr. Gregor Schmidt, Dr. Aleksandra Gumienny and Dr. Mariangela Di Tacchio for their excellent support throughout the project.

This article is dedicated to the memory of Josep (Pepe) Casadesús.

## Data availability

All relevant data are available from the corresponding author on reasonable request.

## Notes

### Competing Interest Statement

The authors have declared no competing interest.

