## Supplementary Material for "Enhancing the performance of Magnets photosensors through directed evolution"

Supplementary Information
for

### Enhancing the performance of Magnets photosensors through directed evolution

by

Armin Baumschlager, Yanik Weber, David Cánovas, Sara Dionisi, Mustafa Khammash

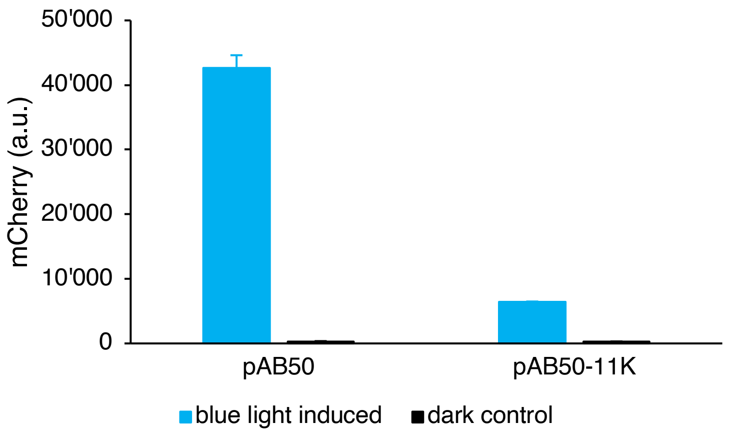


Supplementary Figure 1: Expression level of pAB50 and pAB50-11k with reduced RBS strength using wild-type (WT) Opto-T7RNAP*(563) and saturating light-induction (3.85 $W/m^2$). Diagram shows mean mCherry expression values and standard deviation of three (n=3) biological replicates measured after 5h incubation time measured through flow cytometry.

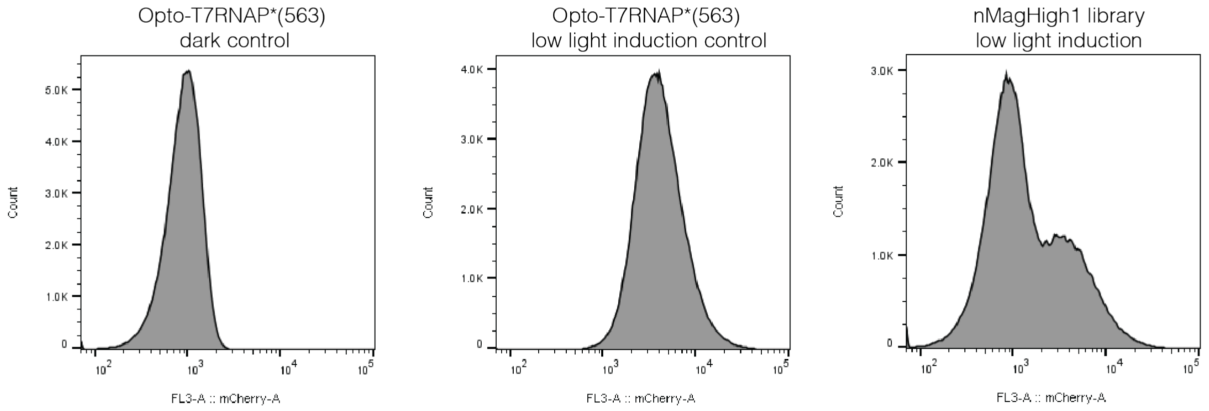


Supplementary Figure 2: mCherry expression level of AB363[1] in the dark (left) and induced with low intensity light (middle) in comparison to the nMagHigh1 library induced with non-saturating (0.96 W $cm^{-2}$) 465-nm light blue light (right). Histograms show mCherry fluorescence obtained during FACS as described in the Materials and Methods section.

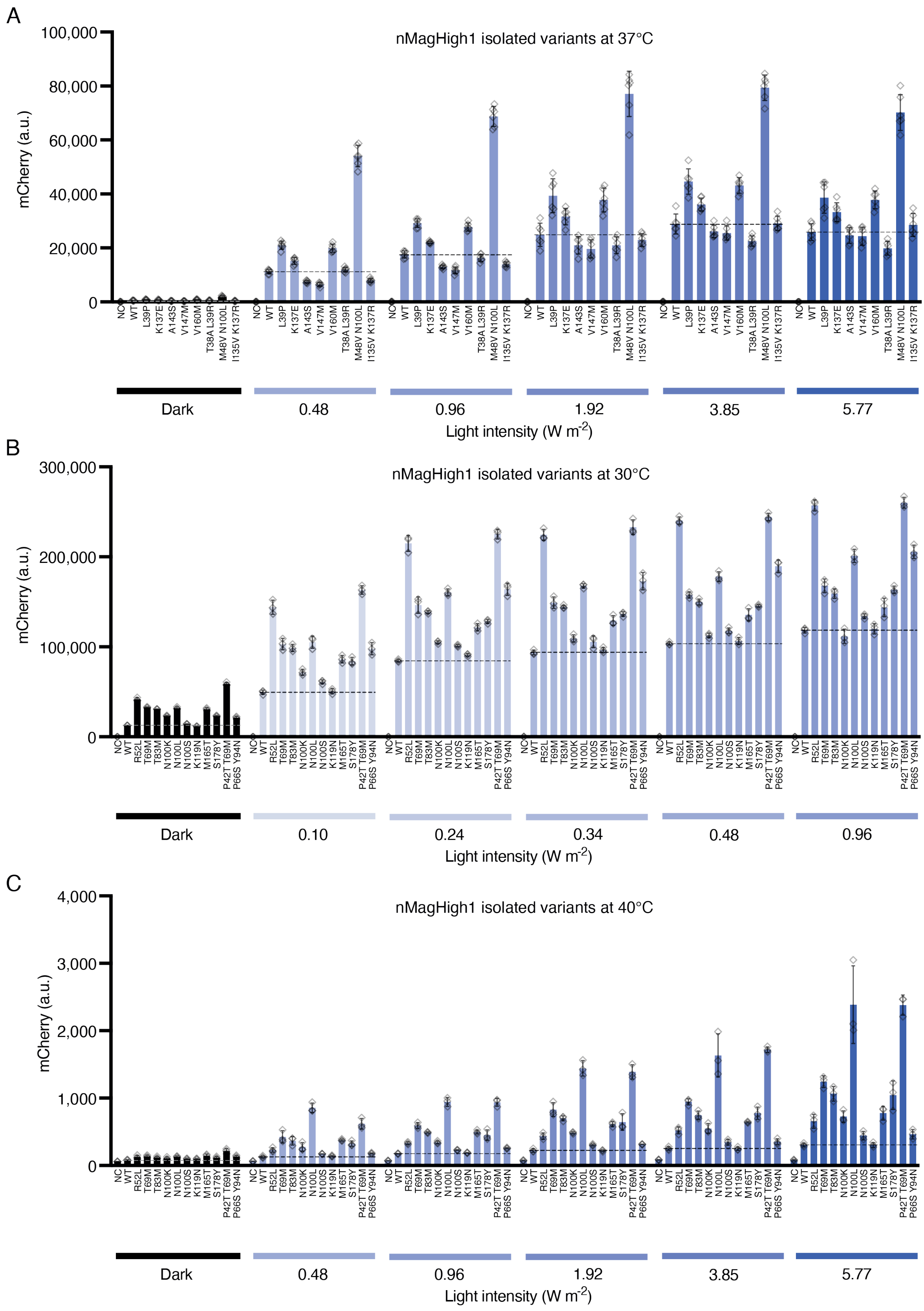


Supplementary Figure 3: Characterization of identified mutations in nMagHigh1 incubated at 37°C (A), 30°C (B) and 40°C (C) through flow cytometry in comparison to the wild-type Opto-T7RNAP*(563) regulator and a negative control containing the mCherry expression plasmid and a second empty plasmid that does not contain the optogenetic regulator. mCherry expression values were acquired after 5h incubation at the indicated light intensity and temperature. Shown are the mean fluorescence values and standard deviation as well as individual data points of at least three (n=3) biological replicates.

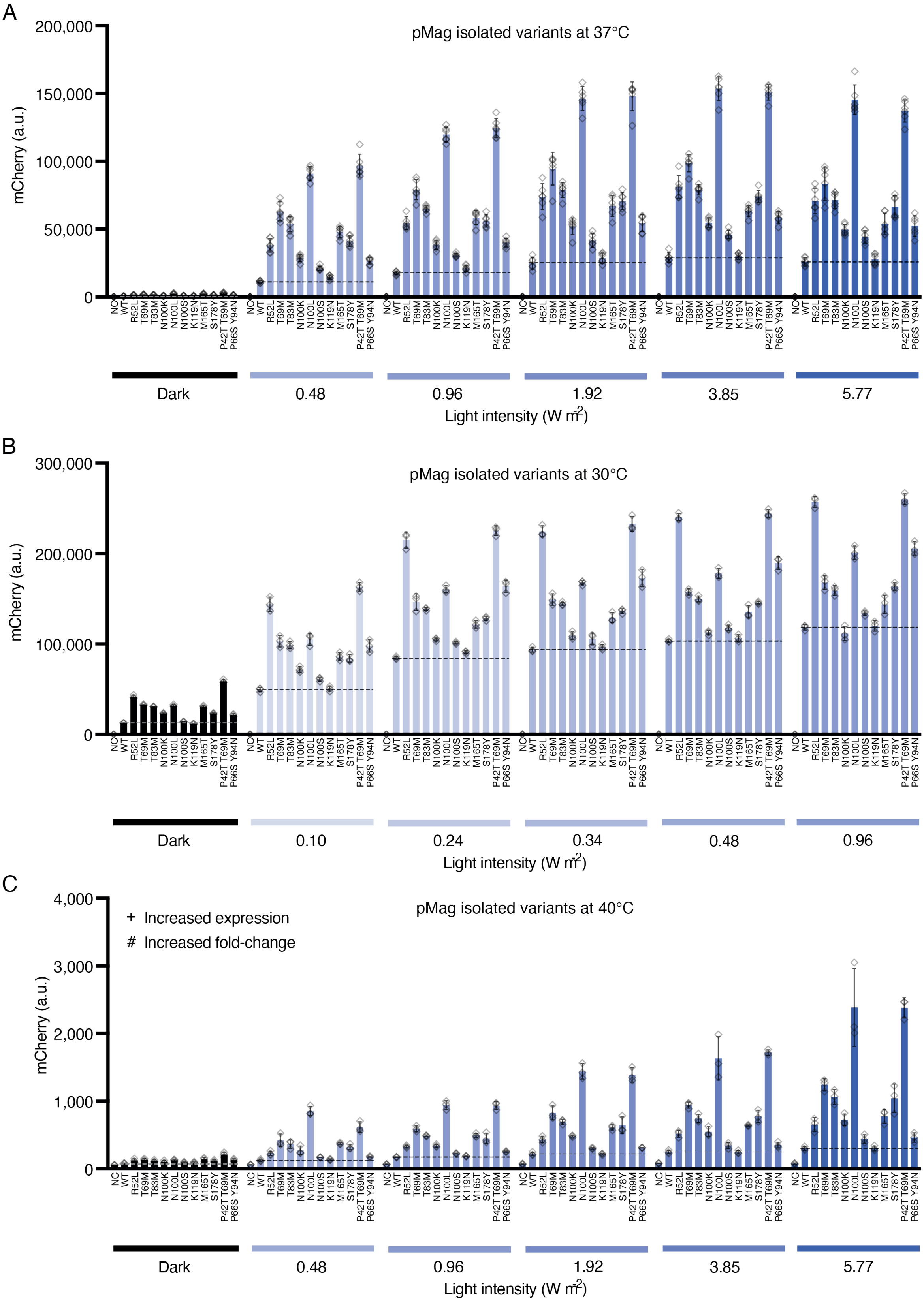


Supplementary Figure 4: Characterization of identified mutations in pMag incubated at 37°C (A), 30°C (B) and 40°C (C) through flow cytometry in comparison to the wild-type Opto-T7RNAP*(563) regulator and a negative control containing the mCherry expression plasmid and a second empty plasmid that does not contain the optogenetic regulator. mCherry expression values were acquired after 5h incubation at the indicated light intensity and temperature. Shown are the mean fluorescence values and standard deviation as well as individual data points of at least three (n=3) biological replicates.

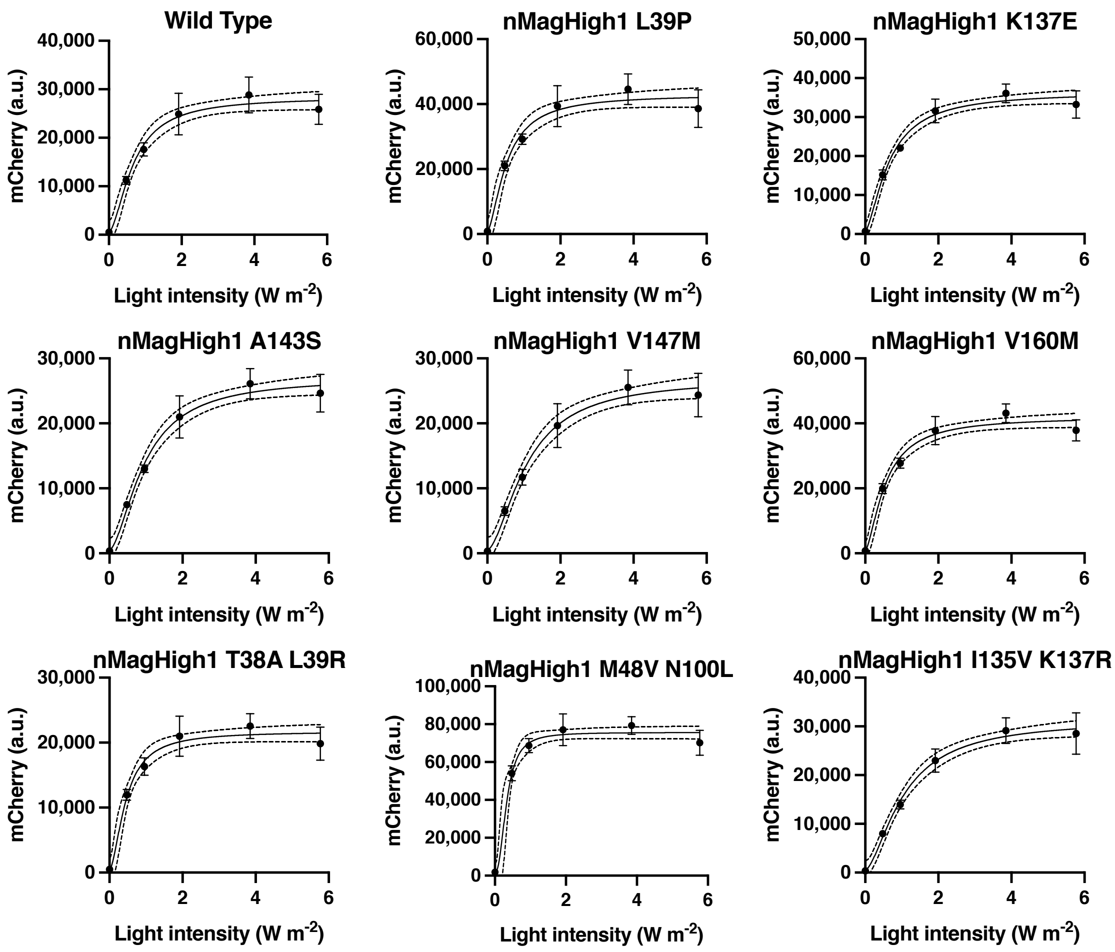


Supplementary Figure 5: Dose-response curves of mCherry fluorescence of Opto-T7RNAP*(563) and different nMagHigh1 variants in response to varying light intensities. Cultures were incubated for 5h at 37°C and endpoints measured through flow cytometry. Shown are mean and standard deviation of six biological replicates (n=6) and mathematical fits (solid lines) with lower and upper 95% profile-likelihood confidence limits (dashed lines).

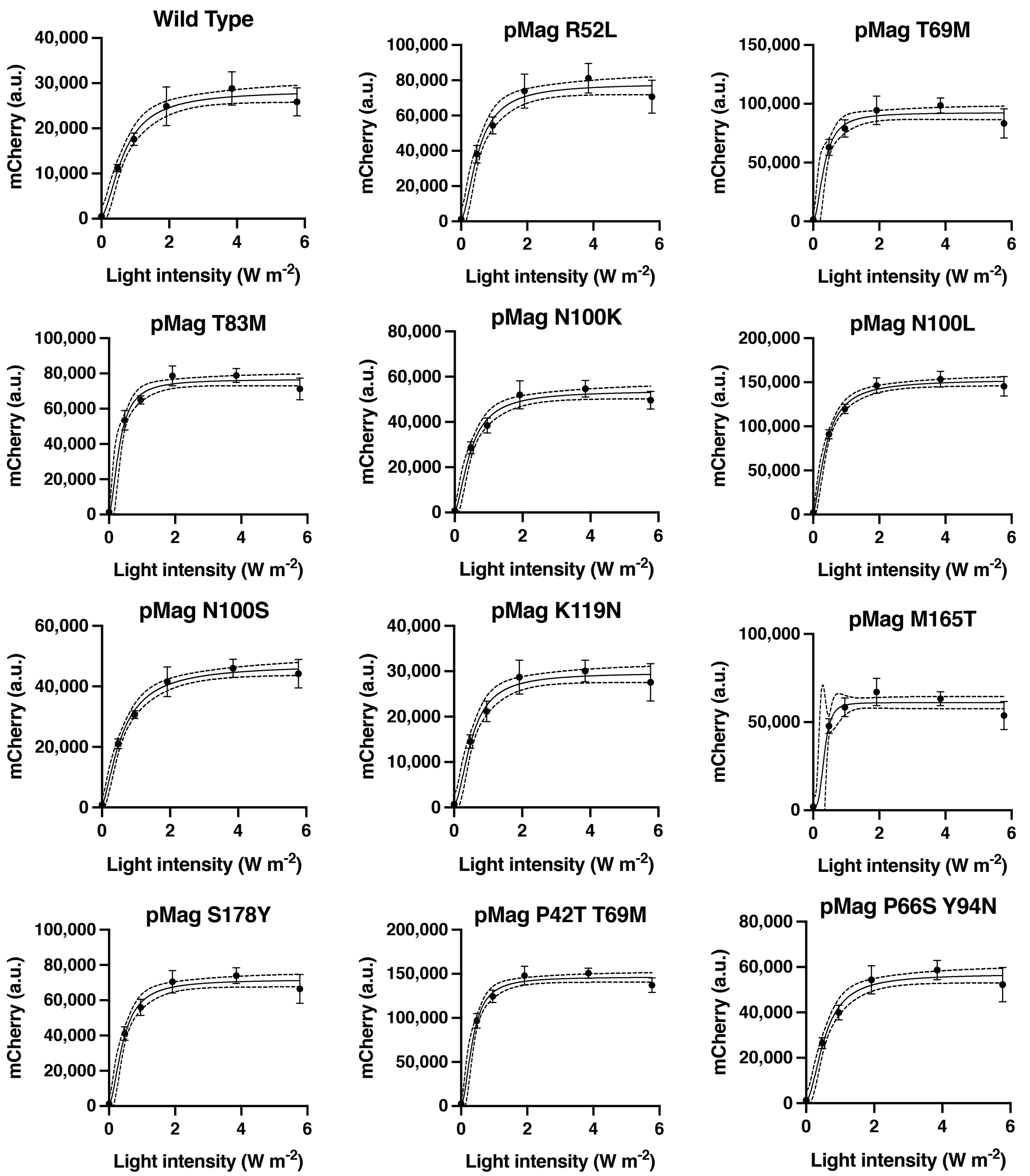


Supplementary Figure 6: Dose-response curves of mCherry fluorescence of Opto-T7RNAP*(563) and different pMag variants in response to varying light intensities. Cultures were incubated for 5h at 37°C and endpoints measured through flow cytometry. Shown are mean and standard deviation of six biological replicates (n=6) and mathematical fits (solid lines) with lower and upper 95% profile-likelihood confidence limits (dashed lines).

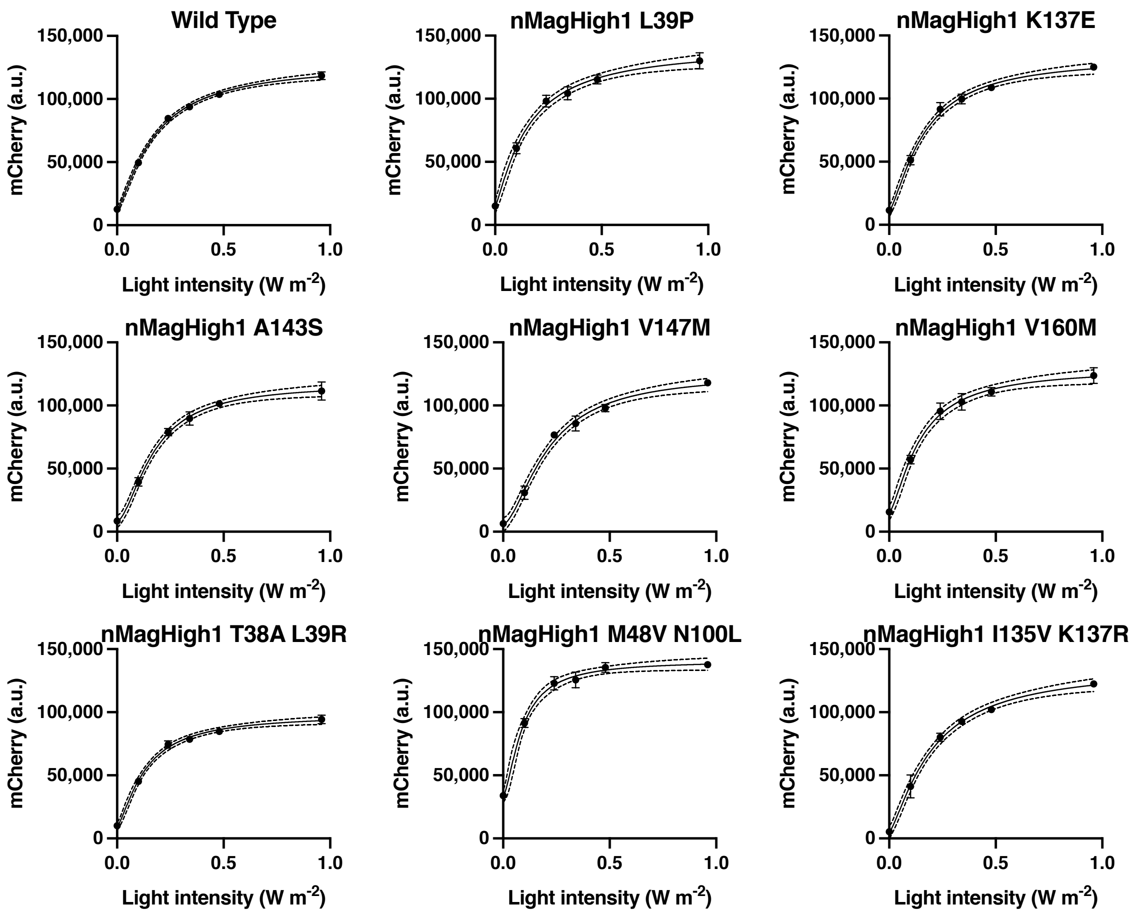


Supplementary Figure 7: Dose-response curves of mCherry fluorescence of Opto-T7RNAP*(563) and different nMagHigh1 variants in response to varying light-intensities. Cultures were incubated for 5h at 30°C and endpoints measures through flow cytometry. Shown are mean and standard deviation of three biological replicates (n=3) and mathematical fits (solid lines) with lower and upper 95% profile-likelihood confidence limits (dashed lines).

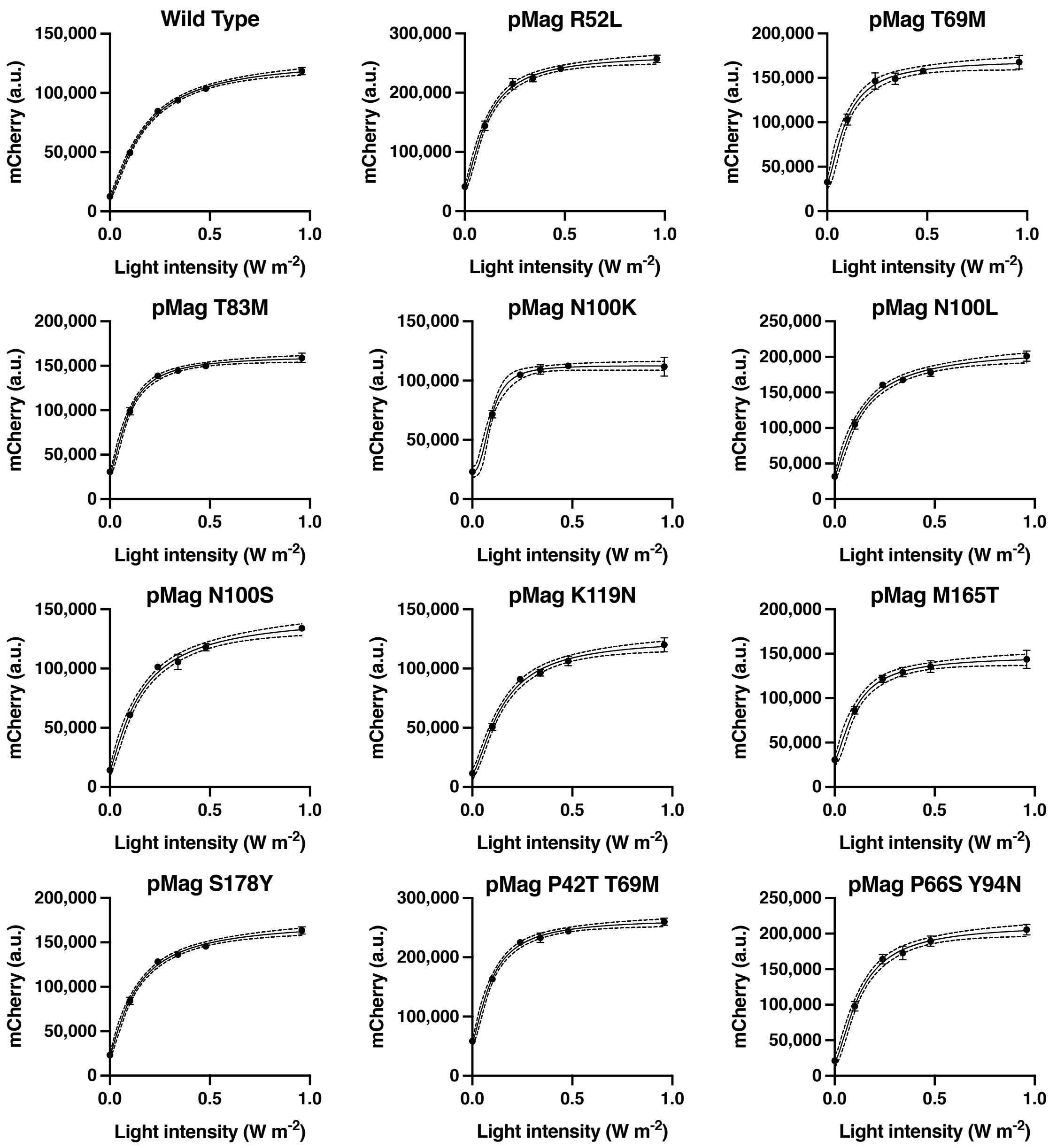


Supplementary Figure 8: Dose-response curves of mCherry fluorescence in response to varying light-intensity of Opto-T7RNAP*(563) and different pMag variants. Cultures were incubated for 5h at 30°C and endpoints measures through flow cytometry. Shown are mean and standard deviation of three biological replicates (n=3) and mathematical fits (solid lines) with lower and upper 95% profile-likelihood confidence limits (dashed lines).

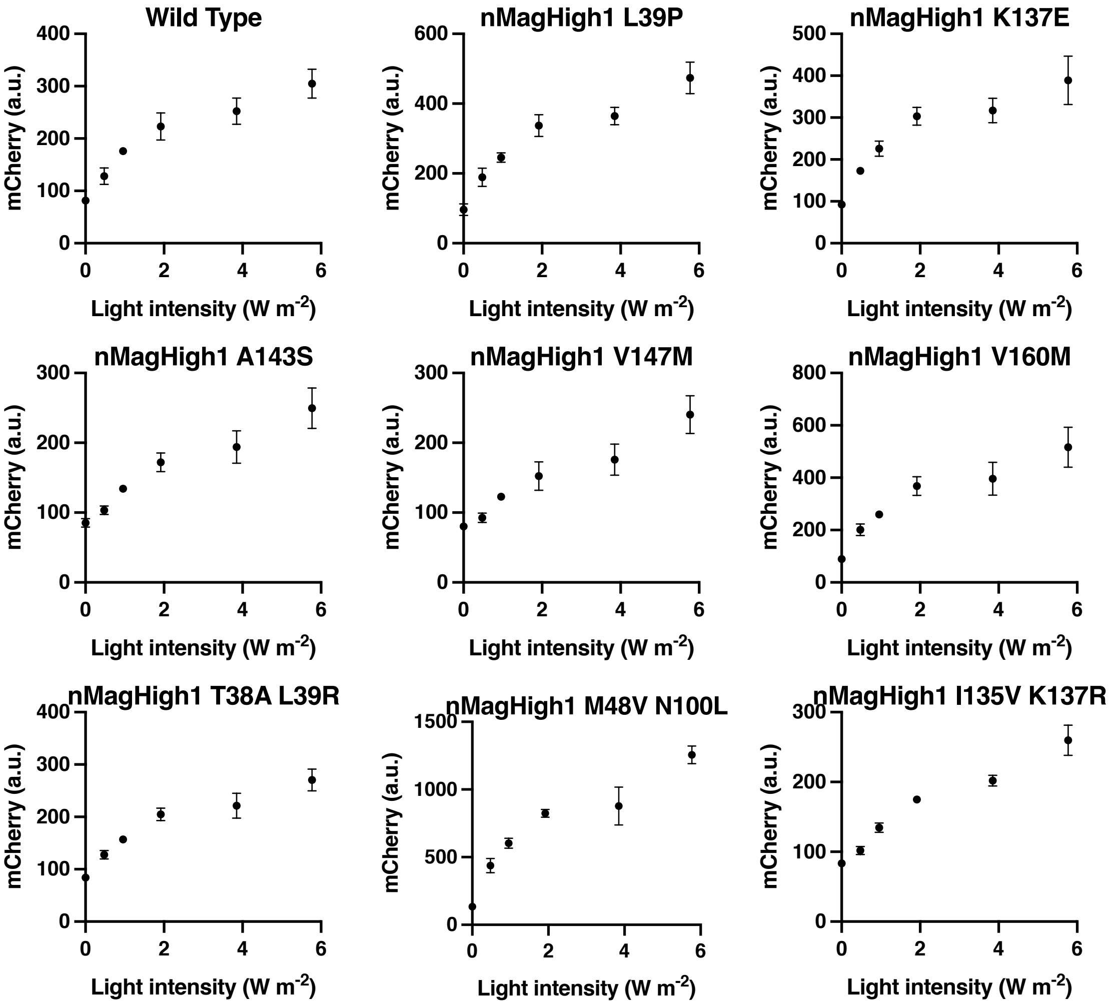


Supplementary Figure 9: Dose-response curves of mCherry fluorescence in response to varying light-intensity of Opto-T7RNAP*(563) and different nMagHigh1 variants. Cultures were incubated for 5h at 40°C and endpoints measured through flow cytometry. Shown are mean and standard deviation of three biological replicates (n=3).

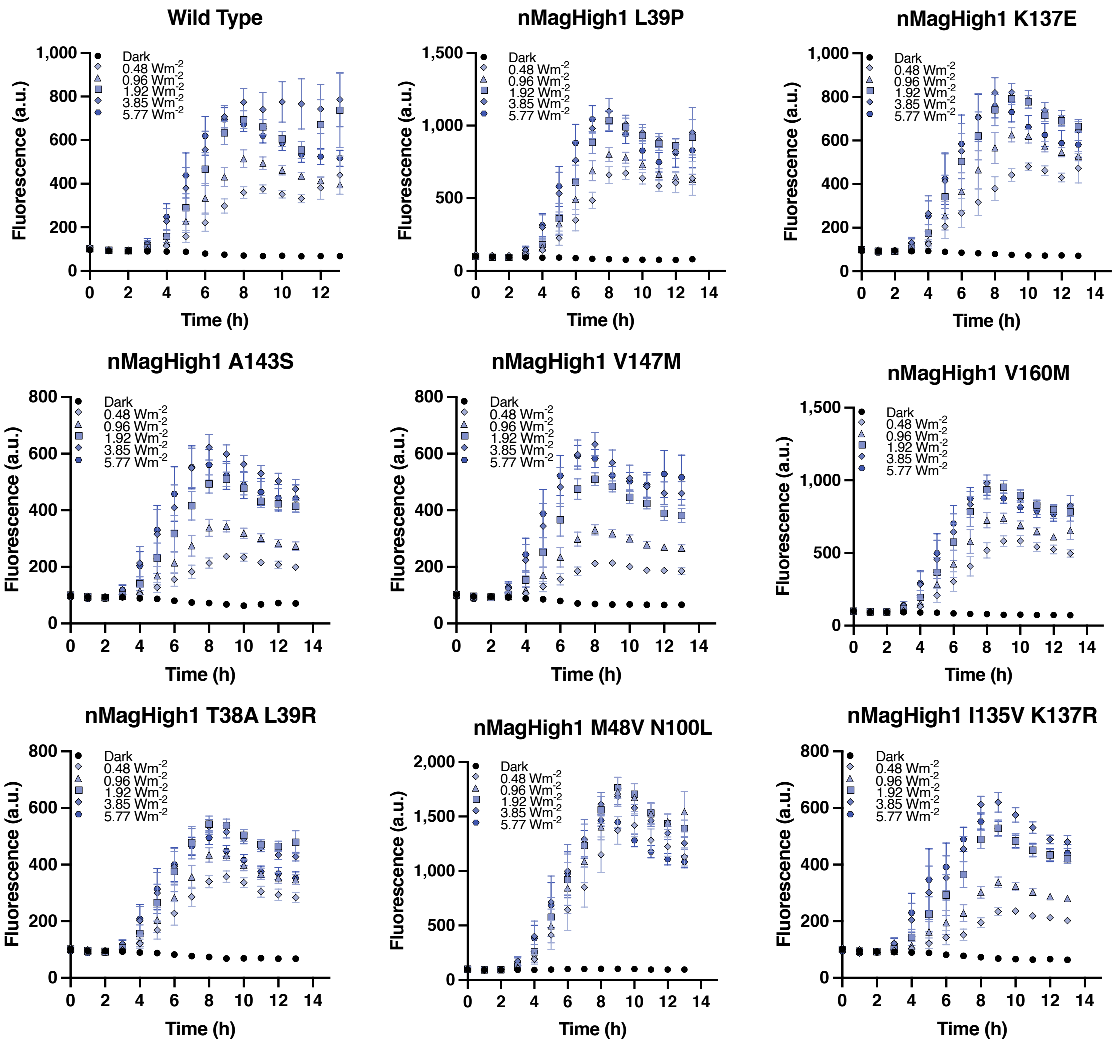


Supplementary Figure 10: Expression of mCherry fluorescence measured through spectrophotometry over time of Opto-T7RNAP*(563) and different nMagHigh1 variants in response to varying light intensities. Cultures were incubated at 37°C. Shown are mean and standard error of the mean fluorescence values of at least six (n=6-9) biological replicates.

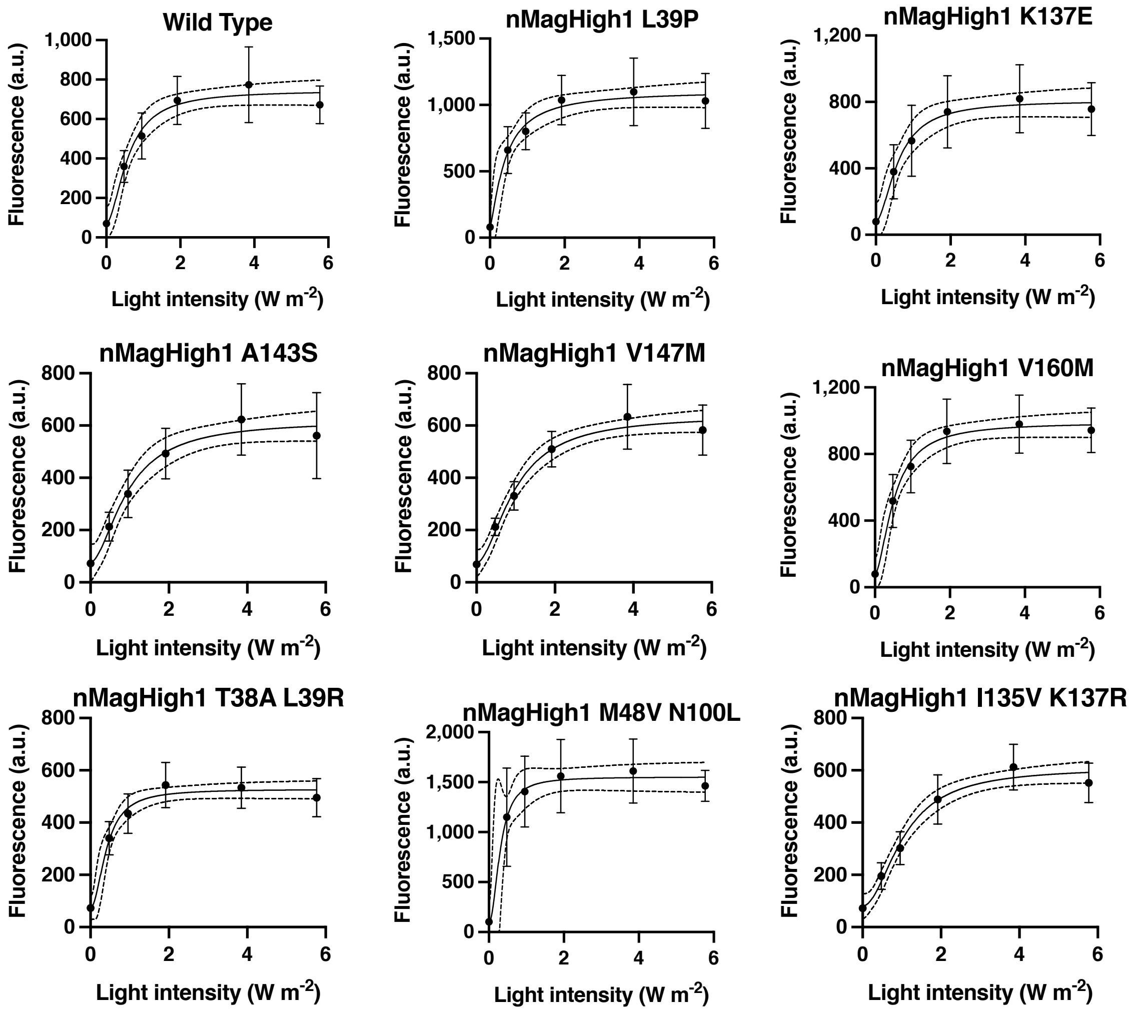


V

Supplementary Figure 11: Expression of mCherry fluorescence measured through spectrophotometry at timepoint 8h of Opto-T7RNAP*(563) and different nMagHigh1 variants in response to varying light intensities. Cultures were incubated at 37°C. Shown are mean and standard error of the mean fluorescence values at least six (n=6-9) biological replicates.

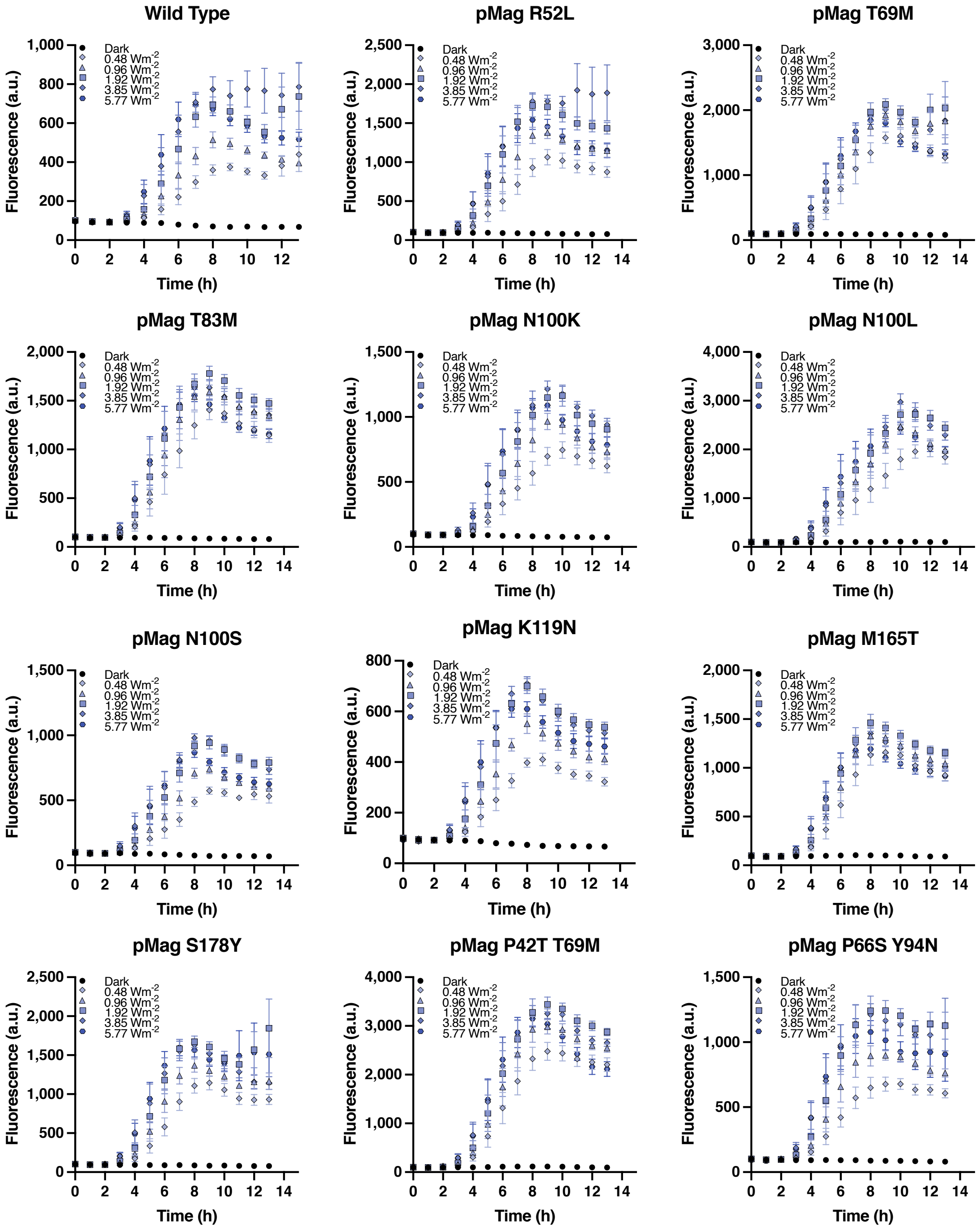


Supplementary Figure 12: Expression of mCherry fluorescence measured through spectrophotometry over time of Opto-T7RNAP*(563) and different pMag variants in response to varying light-intensities. Cultures were incubated at 37°C. Shown are mean and standard error of the mean fluorescence values of at least six (n=6-9) biological replicates.

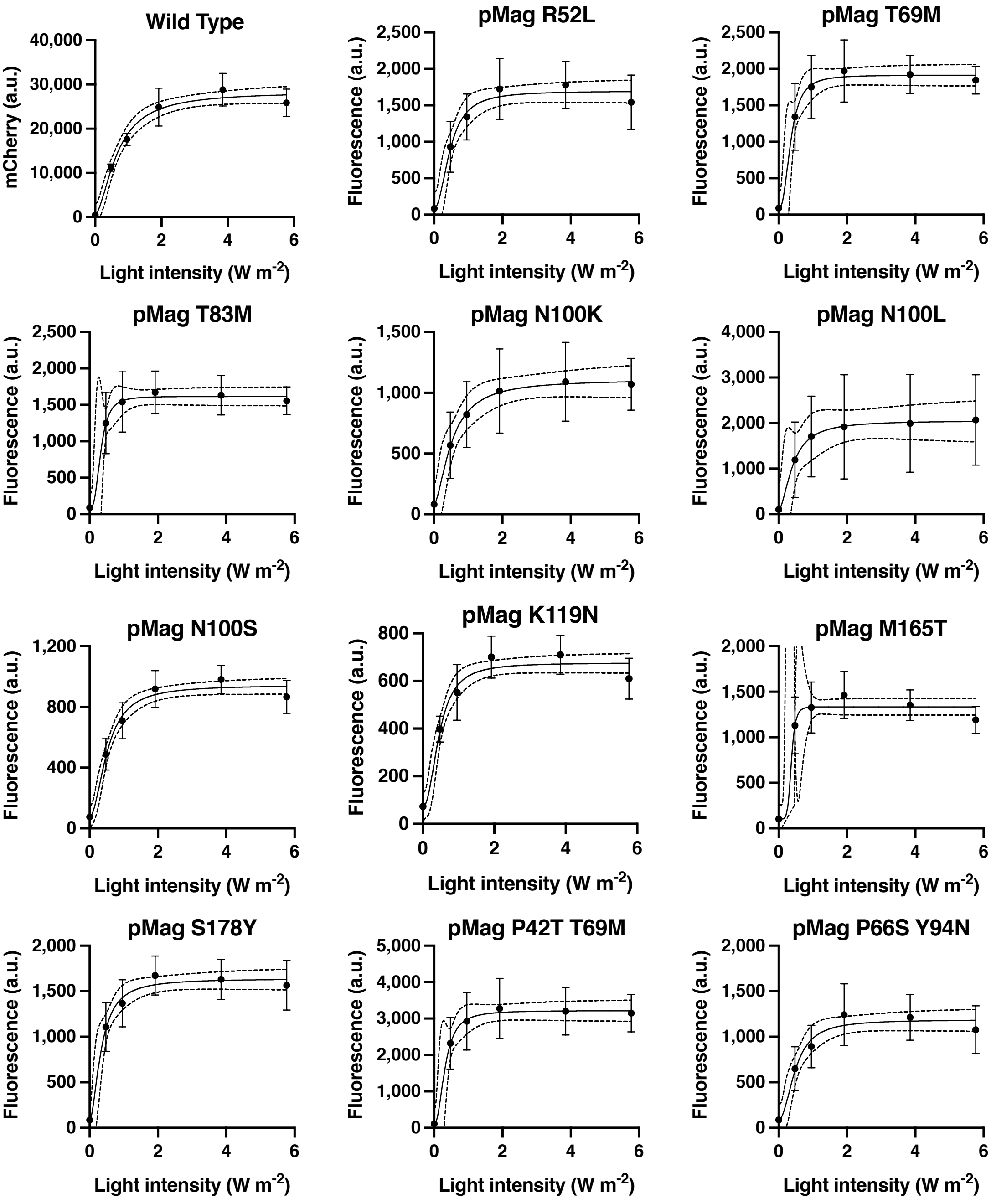


Supplementary Figure 13: Expression of mCherry fluorescence measured through spectrophotometry at timepoint 8h of Opto-T7RNAP*(563) and different nMagHigh1 variants in response to varying light-intensities. Cultures were incubated at 37°C. Shown are mean fluoresence values of of at least six (n=6-9) biological replicates.

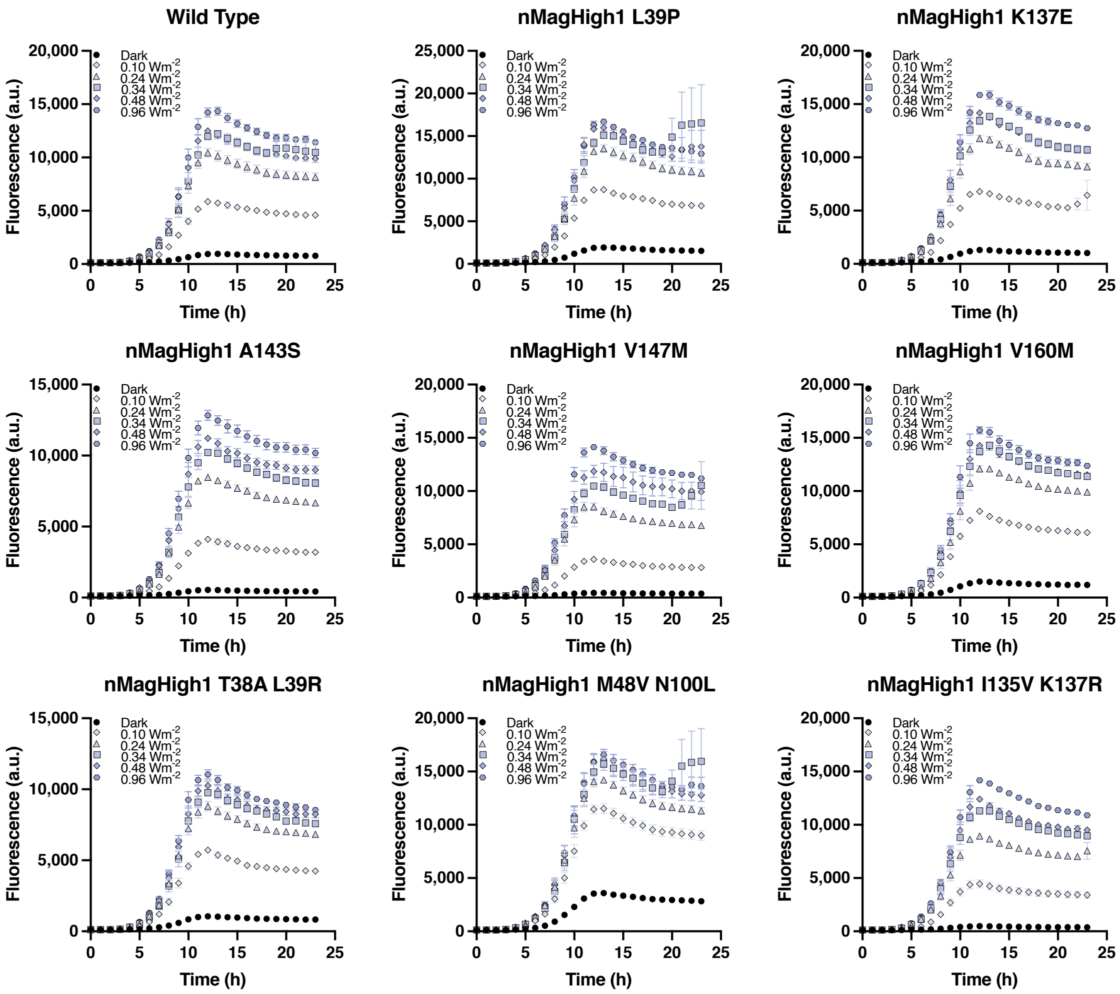


Supplementary Figure 14: Expression of mCherry fluorescence measured through spectrophotometry over time of Opto-T7RNAP*(563) and different nMagHigh1 variants in response to varying light-intensities. Cultures were incubated at 30°C. Shown are mean and standard error of the mean fluorescence values of at least six (n=6-9) biological replicates.

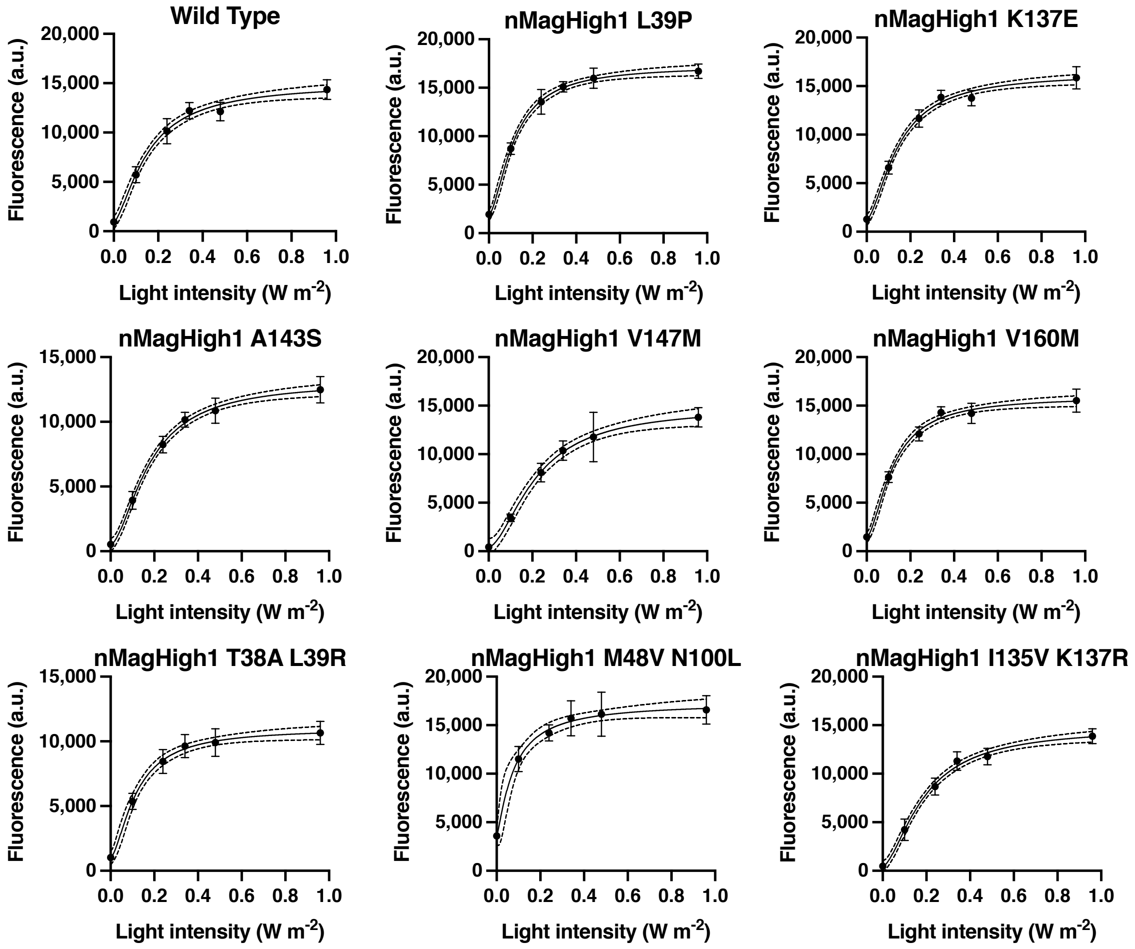


Supplementary Figure 15: Expression of mCherry fluorescence measured through spectrophotometry at timepoint 13h of Opto-T7RNAP*(563) and different nMagHigh1 variants in response to varying light-intensities. Cultures were incubated at 30°C. Shown are mean and standard error of the mean fluorescence values of at least six (n=6-9) biological replicates.

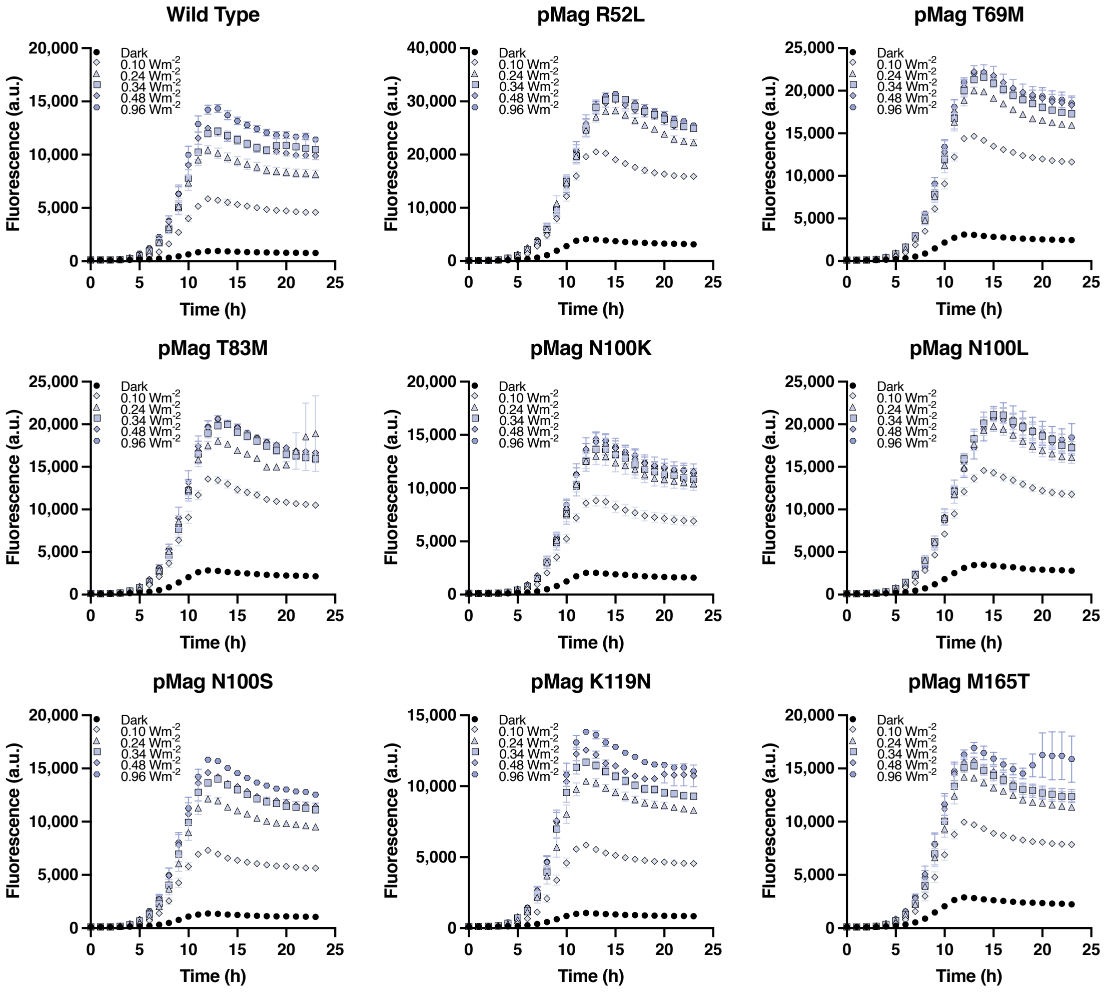


Supplementary Figure 16: Expression of mCherry fluorescence measured through spectrophotometry over time in response to varying light-intensity of Opto-T7RNAP*(563) and different pMag variants. Cultures were incubated at 30°C. Shown are mean and standard error of the mean fluorescence values of at least six (n=6-9) biological replicates.

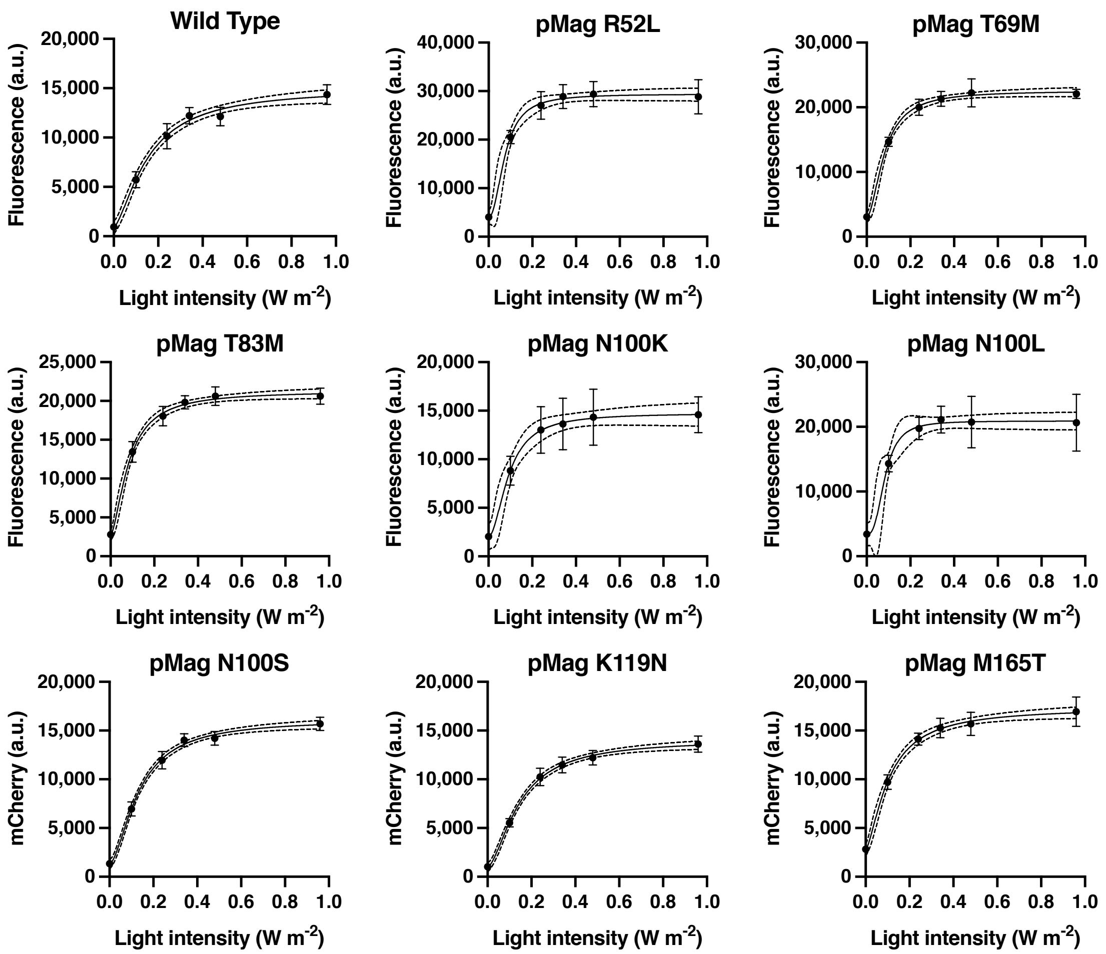


Supplementary Figure 17: Expression of mCherry fluorescence measured through spectrophotometry at timepoint 13h in response to varying light-intensity of Opto-T7RNAP*(563) and different pMag variants and at timepoint 15h for pMag N100L due to the later expression (see Supplementary Figure X30Dp). Cultures were incubated at 30°C. Shown are mean fluorescence values of at least six biological replicates (n=6-9).

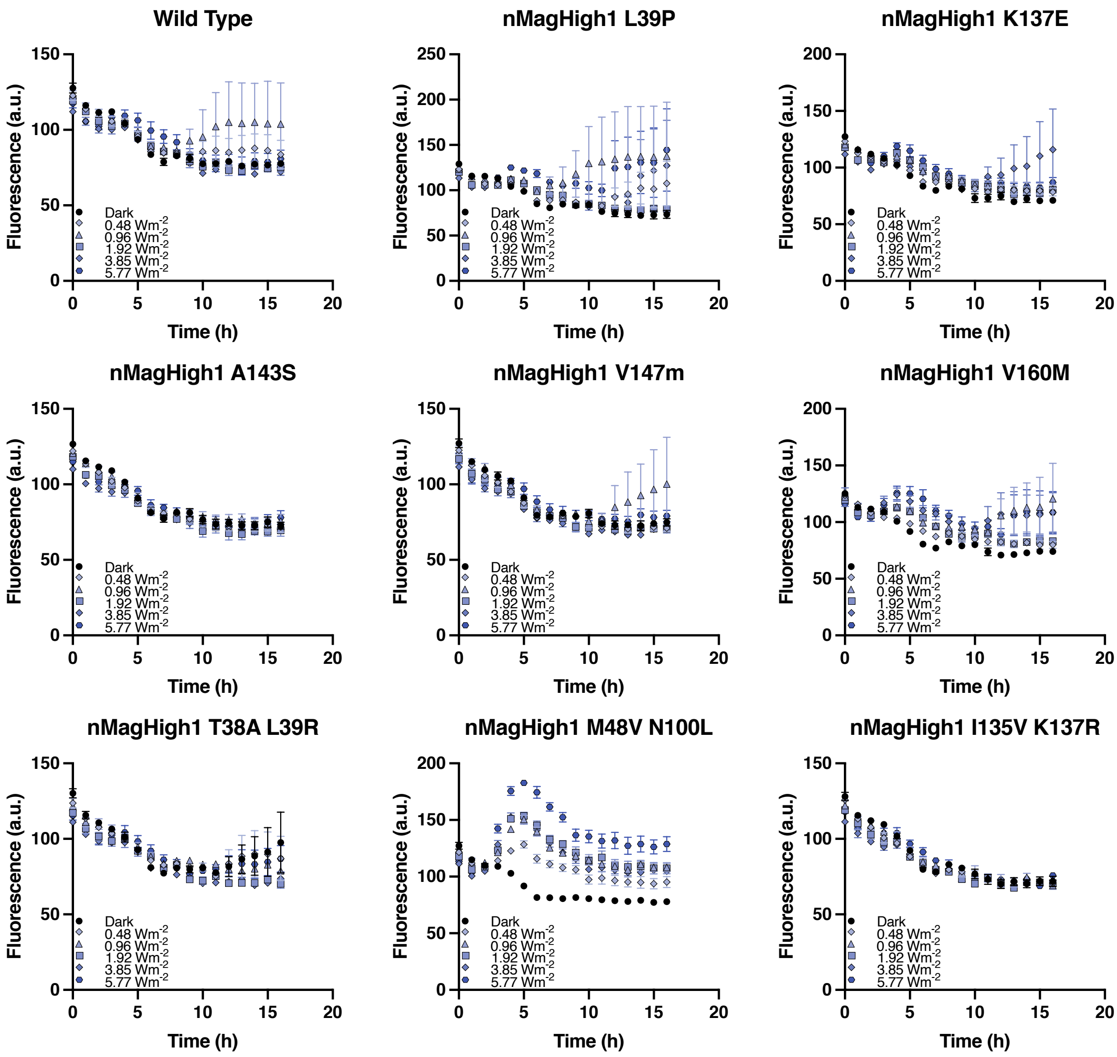


Supplementary Figure 18: Expression of mCherry fluorescence measured through spectrophotometry over time in response to varying light-intensity of Opto-T7RNAP*(563) and different nMagHigh1Mag variants. Cultures were incubated at 40°C. Shown are mean and standard error of the mean fluorescence values of six (n=6) biological replicates.

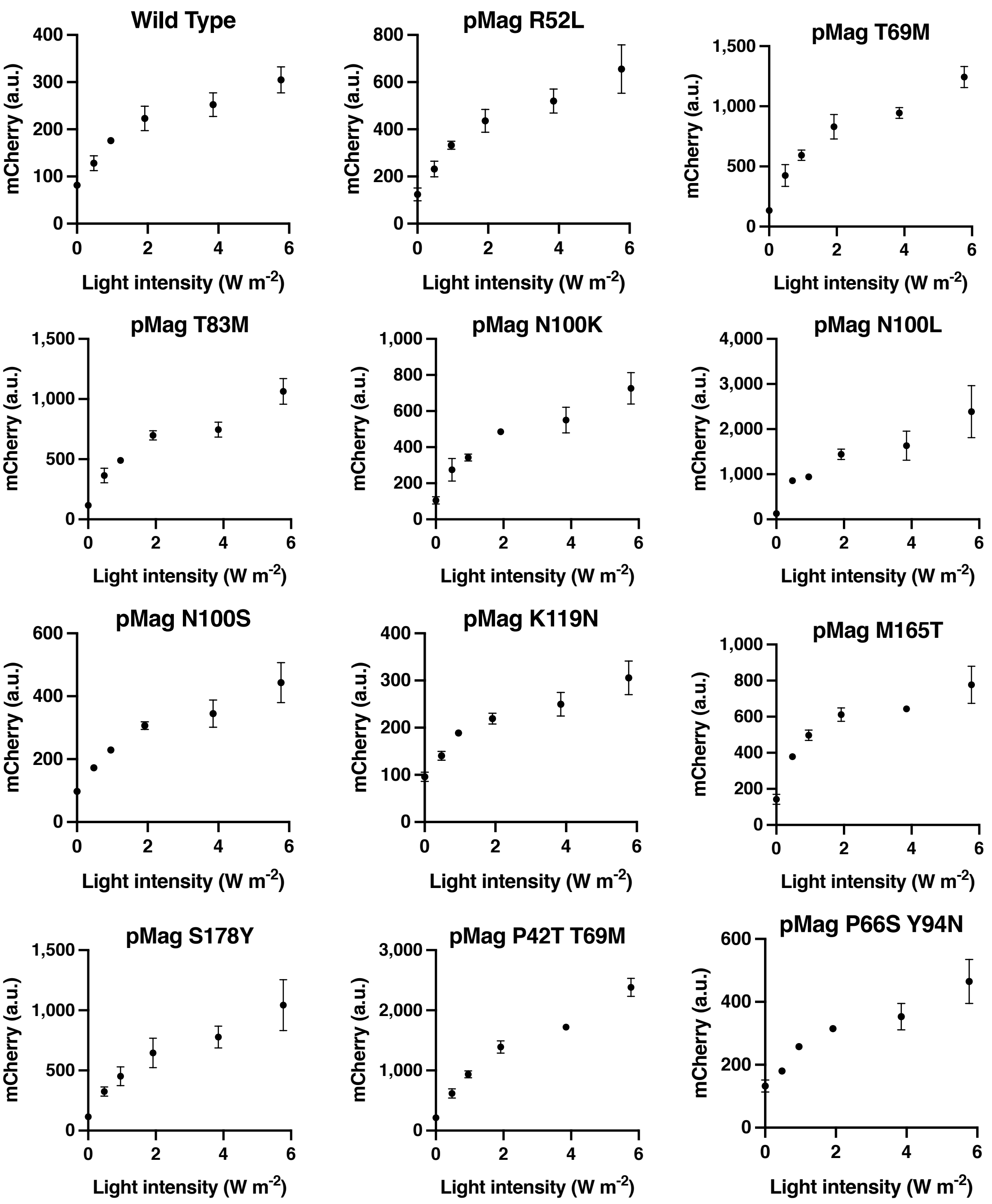


Supplementary Figure 19: Dose-response curves of mCherry fluorescence in response to varying light-intensity of Opto-T7RNAP*(563) and different pMag variants. Cultures were incubated for 5h at 40°C and endpoints measures through flow cytometry. Shown are mean and standard deviation of three biological replicates (n=3).

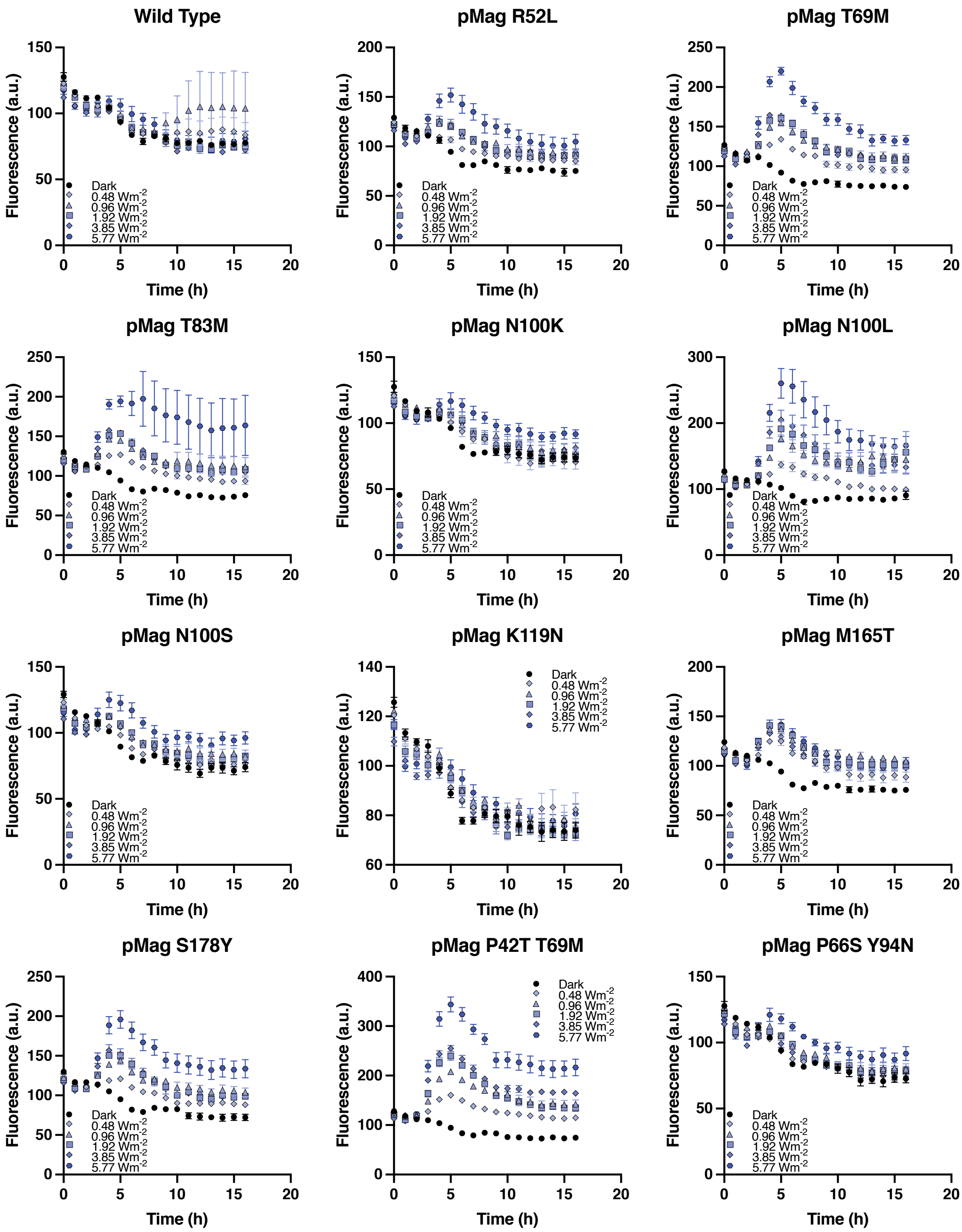


Supplementary Figure 20: Expression of mCherry fluorescence measured through spectrophotometry over time in response to varying light-intensity of Opto-T7RNAP*(563) and different pMag variants. Cultures were incubated at 40°C. Shown are mean and standard error of the mean fluorescence values of six (n=6) biological replicates.

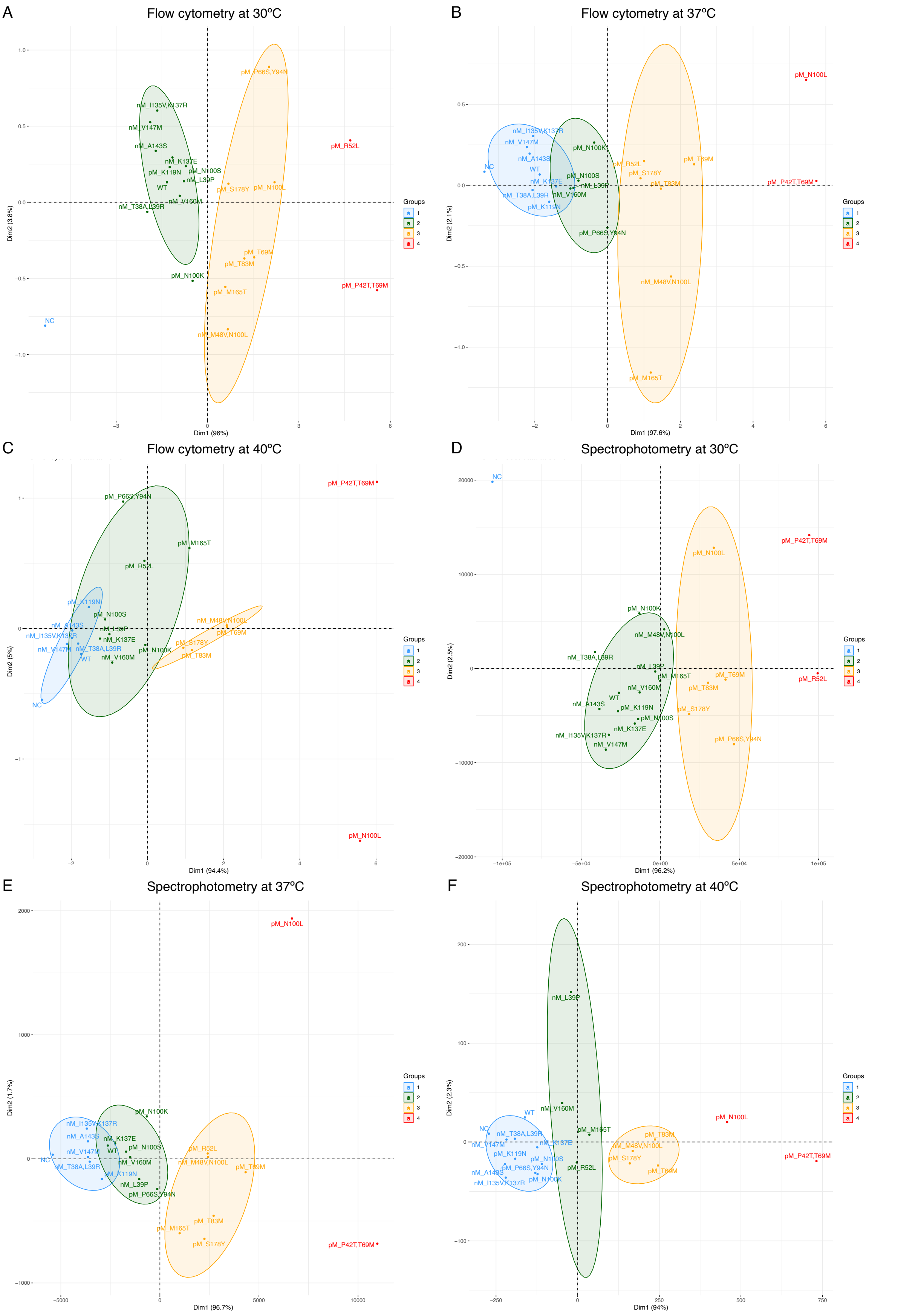


Supplementary Figure 21: PC analysis of the single-cell expression data (flow cytometry; A-C) as well as population data (spectrophotometry; D-F) at the indicated temperatures. Cultures were incubated for 5h and mCherry fluorescence at endpoints was measured by flow cytometry. X-axis shows that PC1 explains 96 %, 97.6% and 94.4% of the differences at 30, 37 and 40 ºC, respectively. Mutant variants are colored according to the clusters selected by hierarchical clustering (see Figure 3 E-F). Red: very high expression; Orange: high expression; Green: medium expression; Blue: low expression.

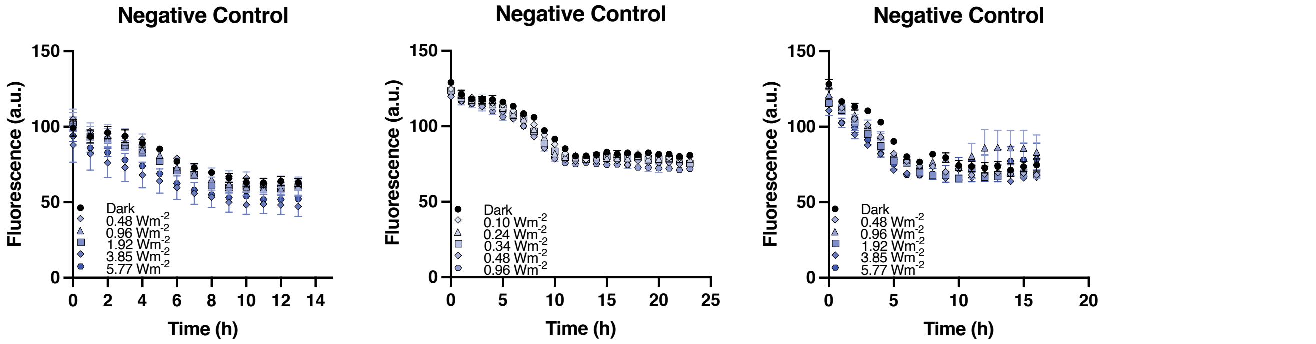


Supplementary Figure 22: Expression of mCherry fluorescence measured through spectrophotometry over time in response to varying light intensity of negative controls. Cultures were incubated at 37°C (left), 30°C (middle) and 40°C (right). Shown are mean and standard error of the mean fluorescence values of at least six (n=6-9) biological replicates.

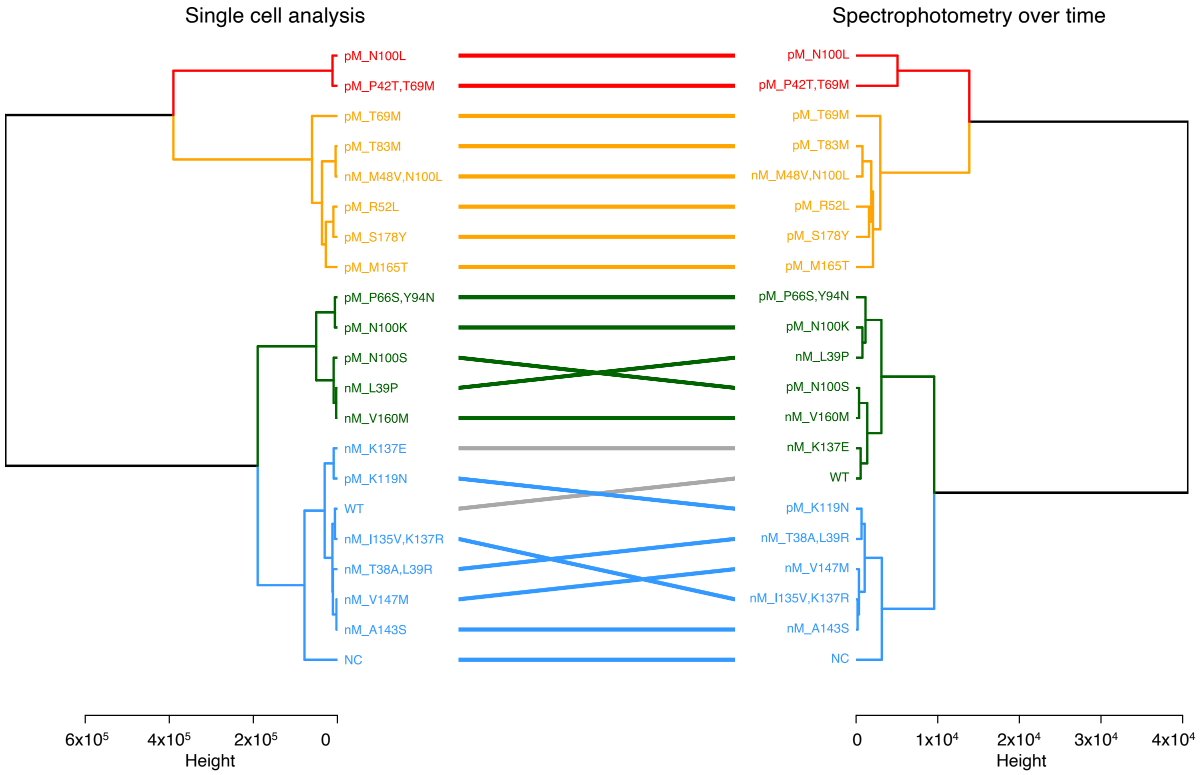


Supplementary Figure 23: Comparison of the dendrograms depicting the hierarchical clustering of variants according to the expression levels at 37ºC for the single cell data (left clustering) and spectrophotometry data (right clustering). Colored horizontal lines connect variants in the same expression clusters in both types of data. Grey horizontal lines connect variants located in different expression clusters in both types of data (i.e. WT and nMag K137E).

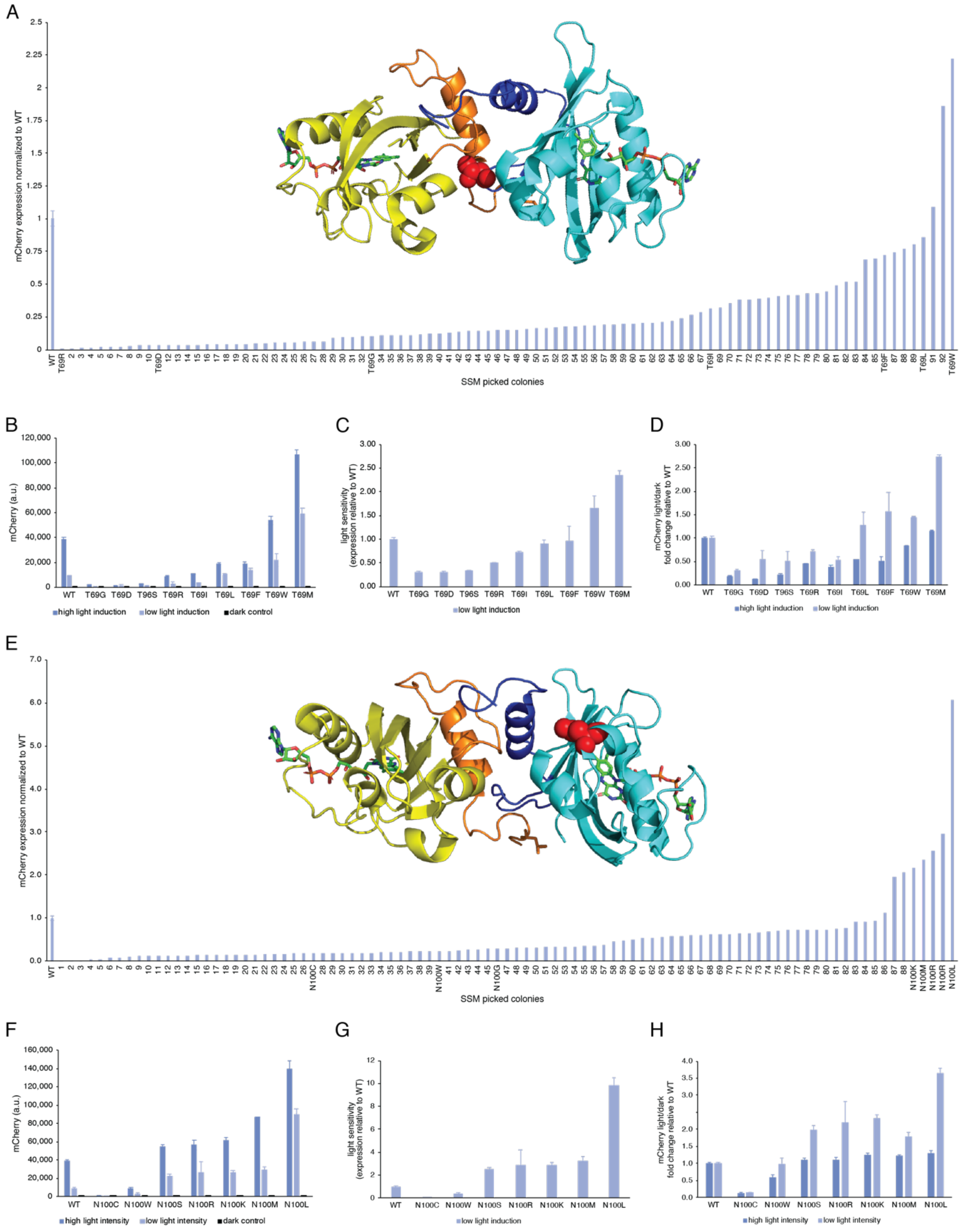


Supplementary Figure 24: Site Saturation Mutagenesis (SSM) of positions T69 and N100 in pMag. (A) mCherry expression level of 93 variants obtained through site saturation mutagenesis at position (A) T69 or (E) N100. Variants were selected to either capture individual variants throughout the range of expression levels as well as variants with increased expression compared to the WT were identified through sequencing and annotated. The inset shows the protein structure of the photosensor VVD in the light-induced dimeric state (PDB code: 3RH8[2]), with (A) amino acid residue T69 in the Ncap of one monomer highlighted as red spheres and in (E) with amino acid residue N100 at the surface of the PAS core in one monomer highlighted as red spheres. nMagHigh1 is shown in yellow and its Ncap in orange. pMag is colored sky blue and its Ncap in darker blue. mCherry expression levels obtained at either high-intensity light induction at 80 a.u. shown in light blue bars, low-intensity light induction at 20 a.u. shown in dark blue bars as well as grown in the dark shown in black, each at steady state after 5h growth in the respective condition for (B) selected T69 variants and (F) selected N100 variants. The light sensitivity of the same variants at low light induction is plotted relative to the WT for (C) T69 and (G) N100 variants to allow for direct comparison of the light sensitivity. Relative fold-change of dark-to-light expression level compared to the WT for (D) T69 and (H) N100 variants. (A, E) show normalized expression levels of individual samples relative to the mean mCherry expression value of the WT which consist of three biological triplicates. Diagrams show mean mCherry expression values and standard deviation of three biological replicates measured after 5h incubation time for (B, F). Relative light sensitivity is calculated from these values by normalizing the mean mCherry expression and standard deviation of variants to the mean mCherry expression of the WT (C, G). Relative fold changes were calculated as the ratio of light-induced to dark controls of the individual samples and the standard error of the ratio[3], which was again normalized to the fold change of the WT under the respective conditions (D, H).

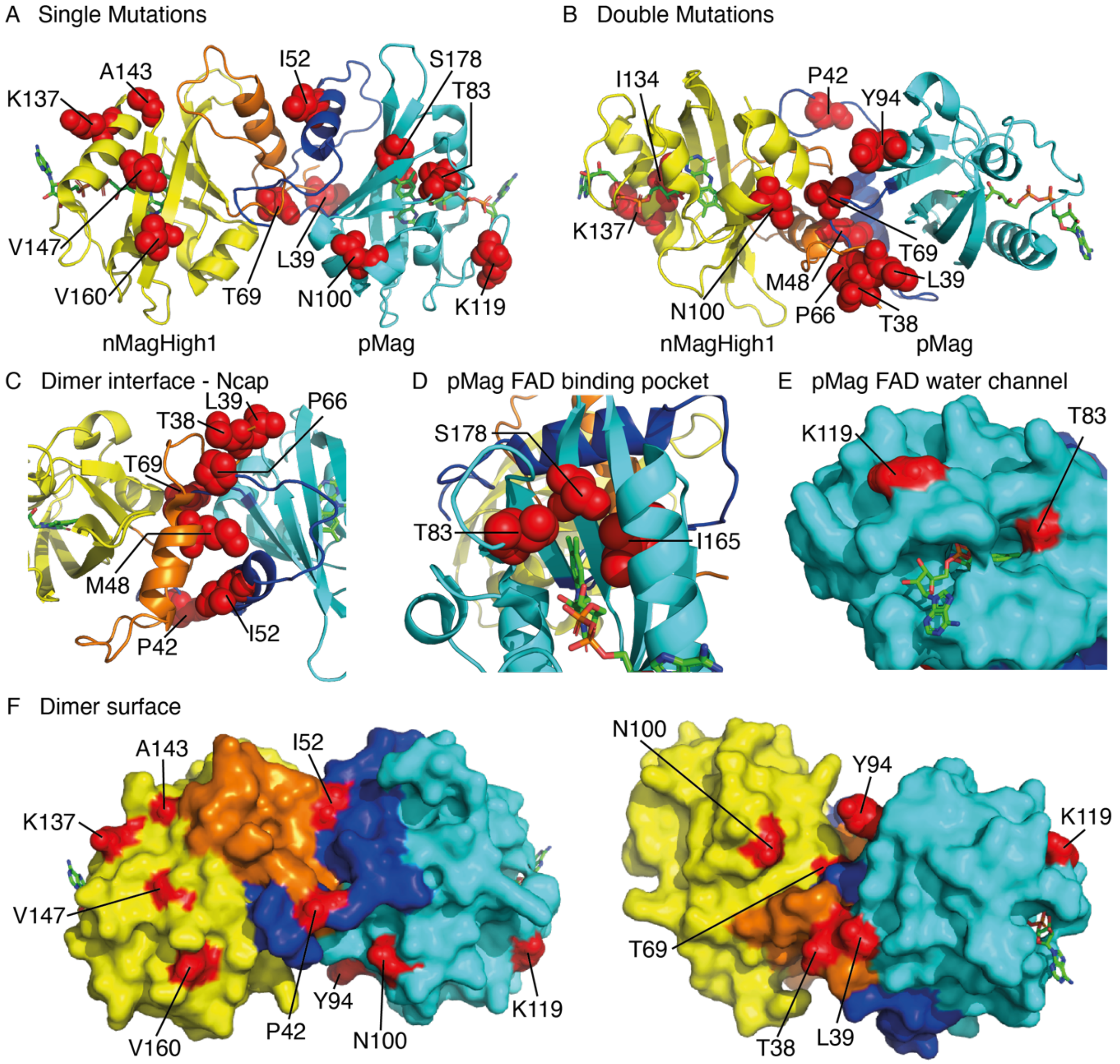


Supplementary Figure 25: Positions of identified mutations in the light-induced VVD homodimer PDB code: 3RH8[2]. All shown protein structures depict a light-induced VVD homodimer which serve as representation of Magnets that differ in 2 positions for pMag (I52R M55R) and nMagHigh1 (I52D M55G M135I M165I). nMagHigh1 is exemplarily shown in yellow and its Ncap in orange. pMag is colored cyan and its Ncap in blue. All shown amino acids are therefore the positions in the VVD photoregulator. The positions of identified mutations are shown in one letter codes next to the residues marked in red. Residue positions from identified (A) single mutations and found in (B) double mutants are highlighted as red spheres. (C) Mutations in the VVD Ncap: T38A, L39P/R, and P42T are located in the N-terminal "latch" (amino acid residues 37 – 44) that wraps around the domain, M48V and R52L are located in the interface of the dimer within the subsequent Aα helix, and P66S and T69M are located in the hinge region of the dimer interface. (C) Detail of flavin binding pocket with, T83, S178 and I165, for which T83M, M165T, and S178Y variants were identified. Residues are shown in red spheres. (E) Surface detail of the entrance of the water channel harboring the flavin chromophore with T83 residue and K119 residue located in the FAD loop shown in light blue. Both residues are shown in red. (F) Surface of the dimer protein with residues of identified variants shown in red

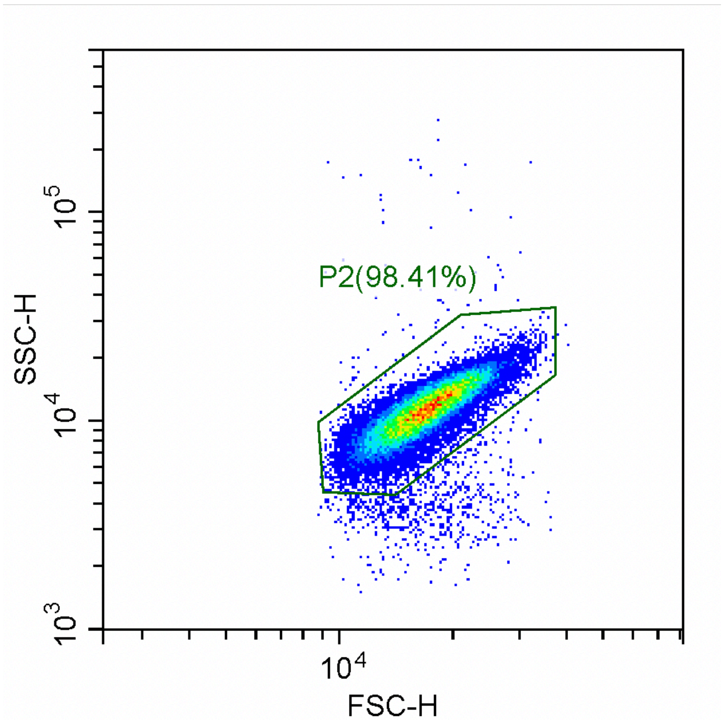


Supplementary Figure 26: Flow cytometry gating strategies. Gate used for the analysis of flow cytometric data of all samples. The SSC-H and FSC-H hexagon gate was drawn by eye on the AB360 control in the CytExpert v.2.1.0.92 and kept constant for all experiments using the same cell type. Shown is a screenshot of gates taken directly from the CytExpert software.

Supplementary Table 1: Identified variant mutations in the respective Magnet domain, summarized as discovered single or double mutations.

| nMagHigh1 | | pMag | |
|---|---|---|---|
| single | double | single | double |
| L39P | T38A, L39R | R52L | P42T, T69M |
| K137E | M48V, N100L | T69M | P66S, Y94N |
| A143S | I135V, K137R | T83M | |
| V147M | | N100K | |
| V160M | | N100L | |
| | | N100S | |
| | | K119N | |
| | | M165T | |
| | | S178Y | |

Supplementary Table 2: Parameters of dose-response curve fits of Opto-T7RNAP*(563) (Wild Type; WT) and nMagHigh1 variants, grown for 5h at 37°C and measured through flow cytometry. Fitting was performed as described in the Materials and Methods section.

| nMagHigh1 variants at 37°C | WT | L39P | K137E | A143S | V147M | V160M | T38A L39R | M48V N100L | I135V K137R |
|---|---|---|---|---|---|---|---|---|---|
| Best-fit values | | | | | | | | | |
| Basal expression (*b*) a.u. | 623 | 936 | 831 | 569 | 552 | 923 | 520 | 1845 | 547 |
| Maximal expression (*t*) a.u. | 28360 | 42942 | 36409 | 27030 | 26910 | 41987 | 21709 | 75667 | 31394 |
| Half-maximal intensity (*I50*) W cm$^{-2}$ | 0.67 | 0.53 | 0.66 | 0.96 | 1.08 | 0.56 | 0.45 | 0.33 | 1.06 |
| Hillslope (n) | 1.7 | 1.6 | 1.5 | 1.7 | 1.7 | 1.5 | 1.7 | 2.4 | 1.6 |
| Lower 95% conf. limit (profile likelihood) | | | | | | | | | |
| Basal expression (*b*) a.u. | -1816 | -2892 | -1329 | -1262 | -1454 | -1887 | -1267 | -3141 | -1454 |
| Maximal expression (*t*) a.u. | 26147 | 39599 | 34165 | 24979 | 24504 | 39324 | 20311 | 72507 | 28789 |
| Half-maximal intensity (*I50*) W cm$^{-2}$ | 0.53 | 0.40 | 0.55 | 0.79 | 0.88 | 0.46 | 0.34 | 0.22 | 0.87 |
| Hillslope (n) | 1.1 | 0.9 | 1.1 | 1.2 | 1.2 | 1.0 | 1.0 | 1.3 | 1.2 |
| Upper 95% conf. limit (profile likelihood) | | | | | | | | | |
| Basal expression (*b*) a.u. | 3065 | 4768 | 2994 | 2401 | 2554 | 3735 | 2309 | 6831 | 2547 |
| Maximal expression (*t*) a.u. | 32079 | 49067 | 39885 | 30386 | 31328 | 46215 | 23882 | 79543 | 36033 |
| Half-maximal intensity (*I50*) W cm$^{-2}$ | 0.86 | 0.70 | 0.80 | 1.21 | 1.45 | 0.68 | 0.56 | 0.42 | 1.39 |
| Hillslope (n) | 2.4 | 2.4 | 2.0 | 2.3 | 2.4 | 2.1 | 2.7 | 6.7 | 2.2 |
| Goodness of Fit | | | | | | | | | |
| Degrees of Freedom | 32 | 32 | 32 | 32 | 32 | 32 | 32 | 32 | 32 |
| R squared | 0.926 | 0.919 | 0.962 | 0.954 | 0.944 | 0.952 | 0.931 | 0.956 | 0.957 |
| Comparison to WT (in %) | | | | | | | | | |
| change in b | 100 | 150 | 133 | 91 | 89 | 148 | 84 | 296 | 88 |
| change in t | 100 | 151 | 128 | 95 | 95 | 148 | 77 | 267 | 111 |
| change in I50 | 100 | 80 | 98 | 143 | 162 | 83 | 68 | 50 | 158 |

Supplementary Table 3: Parameters of dose-response curve fits of Opto-T7RNAP*(563) (Wild Type; WT) and nMagHigh1 variants, grown for 8h at 37°C and measured through spectrophotometry. Fitting was performed as described in the Materials and Methods section.

| nMagHigh1 variants at 37°C | WT | L39P | K137E | A143S | V147M | V160M | T38A L39R | M48V N100L | I135V K137R |
|---|---|---|---|---|---|---|---|---|---|
| Best-fit values | | | | | | | | | |
| Basal expression (*b*) a.u. | 73 | 82 | 81 | 77 | 75 | 80 | 74 | 103 | 81 |
| Maximal expression (*t*) a.u. | 744.7 | 1107 | 808.3 | 618.5 | 638.6 | 989.6 | 527.8 | 1552 | 610.1 |
| Half-maximal intensity (*I50*) W cm$^{-2}$ | 0.59 | 0.41 | 0.61 | 0.95 | 0.99 | 0.52 | 0.42 | 0.31 | 1.05 |
| Hillslope (n) | 1.8 | 1.3 | 1.8 | 1.8 | 1.8 | 1.7 | 2.0 | 2.1 | 2.0 |
| Lower 95% conf. limit (profile likelihood) | | | | | | | | | |
| Basal expression (*b*) a.u. | -14.22 | -37.83 | -36.35 | 4.674 | 22.41 | -20.12 | 26.03 | -112 | 30.73 |
| Maximal expression (*t*) a.u. | 678.1 | 992.7 | 715.9 | 549.3 | 585.6 | 908.1 | 494.8 | 1418 | 560.7 |
| Half-maximal intensity (*I50*) W cm$^{-2}$ | 0.43 | 0.22 | 0.39 | 0.68 | 0.78 | 0.37 | 0.27 | 0.02 | 0.84 |
| Hillslope (n) | 0.9 | 0.3 | 0.6 | 0.9 | 1.2 | 0.8 | 1.0 | 0.1 | 1.3 |
| Upper 95% conf. limit (profile likelihood) | | | | | | | | | |
| Basal expression (*b*) a.u. | 159.9 | 201.5 | 198.4 | 148.1 | 126.8 | 180.9 | 122.5 | 318.2 | 130.7 |
| Maximal expression (*t*) a.u. | 873.3 | 2625 | 1142 | 798.7 | 734.2 | 1168 | 577.9 | n.d. | 692.8 |
| Half-maximal intensity (*I50*) W cm$^{-2}$ | 0.83 | 21.03 | 1.38 | 1.67 | 1.32 | 0.70 | 0.53 | n.d. | 1.36 |
| Hillslope (n) | 3.0 | 2.8 | 3.6 | 3.2 | 2.7 | 2.9 | 3.9 | n.d. | 3.0 |
| Goodness of Fit | | | | | | | | | |
| Degrees of Freedom | 47 | 46 | 50 | 50 | 50 | 50 | 49 | 50 | 49 |
| R squared | 0.802 | 0.812 | 0.702 | 0.782 | 0.882 | 0.830 | 0.847 | 0.737 | 0.885 |
| Comparison to WT (in %) | | | | | | | | | |
| change in b | 100 | 113 | 111 | 105 | 103 | 110 | 102 | 142 | 111 |
| change in t | 100 | 149 | 109 | 83 | 86 | 133 | 71 | 208 | 82 |
| change in I50 | 100 | 70 | 102 | 160 | 167 | 87 | 70 | 52 | 178 |

Supplementary Table 4: Parameters of dose-response curve fits of Opto-T7RNAP*(563) (Wild Type; WT) and pMag variants, grown for 5h at 37°C and measured through flow cytometry. Fitting was performed as described in the Materials and Methods section.

| pMag variants at 37°C | WT | R52L | T69M | T83M | N100K | N100L | N100S | K119N | M165T | S178Y | P42T T69M | P66S Y94N |
|---|---|---|---|---|---|---|---|---|---|---|---|---|
| Best-fit values | | | | | | | | | | | | |
| Basal expression (*b*) a.u. | 622.8 | 1497 | 1601 | 1444 | 902.5 | 2253 | 944.9 | 753.3 | 2042 | 1361 | 2629 | 1431 |
| Maximal expression (*t*) a.u. | 28360 | 78130 | 92569 | 76702 | 53898 | 153134 | 47240 | 29672 | 61087 | 71816 | 146536 | 56961 |
| Half-maximal intensity (*I50*) W cm$^{-2}$ | 0.67 | 0.53 | 0.34 | 0.32 | 0.48 | 0.39 | 0.59 | 0.52 | 0.34 | 0.43 | 0.35 | 0.55 |
| Hillslope (n) | 1.7 | 1.7 | 2.0 | 1.8 | 1.7 | 1.6 | 1.5 | 1.8 | 3.5 | 1.8 | 1.9 | 1.9 |
| Lower 95% conf. limit (profile likelihood) | | | | | | | | | | | | |
| Basal expression (*b*) a.u. | -1816 | -5194 | -6660 | -3208 | -2829 | -4621 | -1795 | -1672 | -3684 | -3558 | -5004 | -2954 |
| Maximal expression (*t*) a.u. | 26147 | 72687 | 87059 | 73412 | 50775 | 147061 | 44369 | 27772 | 57938 | 68056 | 141112 | 53515 |
| Half-maximal intensity (*I50*) W cm$^{-2}$ | 0.53 | 0.41 | 0.19 | 0.22 | 0.38 | 0.33 | 0.50 | 0.42 | 0.14 | 0.34 | 0.28 | 0.45 |
| Hillslope (n) | 1.1 | 1.1 | 0.9 | 1.1 | 1.1 | 1.1 | 1.1 | 1.2 | 1.2 | 1.2 | 1.3 | 1.3 |
| Upper 95% conf. limit (profile likelihood) | | | | | | | | | | | | |
| Basal expression (*b*) a.u. | 3065 | 8197 | 9864 | 6097 | 4636 | 9127 | 3686 | 3181 | 7767 | 6281 | 10263 | 5821 |
| Maximal expression (*t*) a.u. | 32079 | 86549 | 100844 | 81171 | 58519 | 161570 | 51891 | 32401 | 64583 | 76997 | 153253 | 61628 |
| Half-maximal intensity (*I50*) W cm$^{-2}$ | 0.86 | 0.66 | 0.43 | 0.39 | 0.58 | 0.45 | 0.72 | 0.64 | n.d. | 0.51 | 0.41 | 0.67 |
| Hillslope (n) | 2.4 | 2.6 | 4.7 | 3.1 | 2.3 | 2.1 | 2.0 | 2.7 | n.d. | 2.7 | 2.9 | 2.6 |
| Goodness of Fit | | | | | | | | | | | | |
| Degrees of Freedom | 32 | 32 | 32 | 32 | 32 | 32 | 32 | 32 | 32 | 32 | 32 | 32 |
| R squared | 0.926 | 0.927 | 0.922 | 0.962 | 0.950 | 0.978 | 0.963 | 0.933 | 0.916 | 0.952 | 0.972 | 0.941 |
| Comparison to WT (in %) | | | | | | | | | | | | |
| change in b | 100 | 240 | 257 | 232 | 145 | 362 | 152 | 121 | 328 | 219 | 422 | 230 |
| change in t | 100 | 275 | 326 | 270 | 190 | 540 | 167 | 105 | 215 | 253 | 517 | 201 |
| change in I50 | 100 | 79 | 50 | 47 | 71 | 59 | 89 | 78 | 50 | 64 | 52 | 83 |

Supplementary Table 5: Parameters of dose-response curve fits of Opto-T7RNAP*(563) (Wild Type; WT) and pMag variants, grown for 8h at 37°C and measured through spectrophotometry. Fitting was performed as described in the Materials and Methods section.

| pMag variants at 37°C | WT | R52L | T69M | T83M | N100K | N100L | N100S | K119N | M165T | S178Y | P42T T69M | P66S Y94N |
|---|---|---|---|---|---|---|---|---|---|---|---|---|
| Best-fit values | | | | | | | | | | | | |
| Basal expression (*b*) a.u. | 73 | 90 | 91 | 88 | 83 | 103 | 78 | 74 | 103 | 86 | 116 | 92 |
| Maximal expression (*t*) a.u. | 744.7 | 1697 | 1916 | 1616 | 1110 | 2055 | 943.3 | 676.3 | 1335 | 1638 | 3219 | 1187 |
| Half-maximal intensity (*I50*) W $cm^{-2}$ | 0.59 | 0.47 | 0.35 | 0.33 | 0.52 | 0.42 | 0.52 | 0.46 | 0.37 | 0.34 | 0.32 | 0.49 |
| Hillslope (n) | 1.8 | 2.1 | 2.5 | 3.0 | 1.6 | 1.8 | 2.0 | 2.2 | 6.2 | 1.8 | 2.3 | 2.1 |
| Lower 95% conf. limit (profile likelihood) | | | | | | | | | | | | |
| Basal expression (*b*) a.u. | -14.22 | -131.8 | -133.4 | -110.4 | -91.09 | -490.3 | 6.253 | 15.08 | -51.58 | -66.71 | -309.6 | -77.38 |
| Maximal expression (*t*) a.u. | 678.1 | 1547 | 1780 | 1502 | 970.5 | 1647 | 889.8 | 637.3 | 1256 | 1528 | 2956 | 1074 |
| Half-maximal intensity (*I50*) W $cm^{-2}$ | 0.43 | 0.30 | 0.13 | 0.02 | 0.27 | 0.00 | 0.42 | 0.35 | n.d. | 0.17 | 0.04 | 0.30 |
| Hillslope (n) | 0.9 | 0.9 | 0.7 | 0.3 | 0.3 | n.d. | 1.3 | 1.3 | 0.4 | 0.7 | 0.2 | 0.9 |
| Upper 95% conf. limit (profile likelihood) | | | | | | | | | | | | |
| Basal expression (*b*) a.u. | 159.9 | 311.9 | 315.4 | 287.2 | 256.5 | 695.4 | 149.8 | 133.3 | 258.2 | 239.4 | 541.6 | 261.5 |
| Maximal expression (*t*) a.u. | 873.3 | 1938 | 2168 | 2122 | 6311 | n.d. | 1013 | 723.3 | 1434 | 1869 | 6606 | 1382 |
| Half-maximal intensity (*I50*) W $cm^{-2}$ | 0.83 | 0.63 | 0.45 | n.d. | 1815.00 | n.d. | 0.63 | 0.56 | n.d. | 0.45 | 6.19 | 0.69 |
| Hillslope (n) | 3.0 | 4.8 | n.d. | n.d. | 4.3 | n.d. | 2.8 | 3.8 | n.d. | 4.2 | n.d. | 4.5 |
| Goodness of Fit | | | | | | | | | | | | |
| Degrees of Freedom | 47 | 49 | 50 | 49 | 50 | 49 | 50 | 50 | 50 | 48 | 50 | 50 |
| R squared | 0.802 | 0.769 | 0.806 | 0.791 | 0.673 | 0.395 | 0.901 | 0.868 | 0.803 | 0.865 | 0.767 | 0.725 |
| Comparison to WT (in %) | | | | | | | | | | | | |
| change in b | 100 | 124 | 125 | 122 | 114 | 141 | 107 | 102 | 142 | 119 | 160 | 126 |
| change in t | 100 | 228 | 257 | 217 | 149 | 276 | 127 | 91 | 179 | 220 | 432 | 159 |
| change in I50 | 100 | 79 | 60 | 55 | 88 | 71 | 88 | 78 | 63 | 58 | 55 | 83 |

Supplementary Table 6: Parameters of dose-response curve fits of Opto-T7RNAP*(563) (Wild Type; WT) and nMagHigh1 variants, grown for 5h at 30°C and measured through flow cytometry. Fitting was performed as described in the Materials and Methods section.

| nMagHigh1 variants at 30°C | WT | L39P | K137E | A143S | V147M | V160M | T38A L39R | M48V N100L | I135V K137R |
|---|---|---|---|---|---|---|---|---|---|
| Best-fit values | | | | | | | | | |
| Basal expression (*b*) a.u. | 12539 | 14867 | 11310 | 8456 | 5892 | 15556 | 9956 | 33974 | 5147 |
| Maximal expression (*t*) a.u. | 128601 | 141254 | 133081 | 117004 | 125381 | 129201 | 98245 | 140683 | 136077 |
| Half-maximal intensity (*I50*) W cm$^{-2}$ | 0.17 | 0.15 | 0.16 | 0.17 | 0.21 | 0.14 | 0.13 | 0.09 | 0.20 |
| Hillslope (n) | 1.3 | 1.2 | 1.4 | 1.7 | 1.6 | 1.5 | 1.4 | 1.6 | 1.3 |
| Lower 95% conf. limit (profile likelihood) | | | | | | | | | |
| Basal expression (*b*) a.u. | 9720 | 9218 | 6722 | 3627 | 456.1 | 9491 | 6771 | 28669 | -75.39 |
| Maximal expression (*t*) a.u. | 122353 | 129175 | 124277 | 109959 | 115637 | 119450 | 92783 | 134108 | 124188 |
| Half-maximal intensity (*I50*) W cm$^{-2}$ | 0.16 | 0.12 | 0.14 | 0.15 | 0.18 | 0.12 | 0.11 | 0.08 | 0.17 |
| Hillslope (n) | 1.1 | 0.9 | 1.1 | 1.4 | 1.3 | 1.0 | 1.1 | 1.1 | 1.0 |
| Upper 95% conf. limit (profile likelihood) | | | | | | | | | |
| Basal expression (*b*) a.u. | 15357 | 20514 | 15893 | 13280 | 11296 | 21618 | 13140 | 39279 | 10362 |
| Maximal expression (*t*) a.u. | 136930 | 164289 | 146345 | 126535 | 139879 | 145738 | 106163 | 151192 | 156207 |
| Half-maximal intensity (*I50*) W cm$^{-2}$ | 0.20 | 0.22 | 0.20 | 0.20 | 0.26 | 0.18 | 0.15 | 0.11 | 0.27 |
| Hillslope (n) | 1.5 | 1.6 | 1.7 | 2.1 | 2.0 | 2.0 | 1.8 | 2.3 | 1.7 |
| Goodness of Fit | | | | | | | | | |
| Degrees of Freedom | 14 | 14 | 14 | 14 | 14 | 14 | 14 | 14 | 14 |
| R squared | 0.997 | 0.989 | 0.993 | 0.991 | 0.990 | 0.986 | 0.994 | 0.989 | 0.991 |
| Comparison to WT (in %) | | | | | | | | | |
| change in b | 100 | 119 | 90 | 67 | 47 | 124 | 79 | 271 | 41 |
| change in t | 100 | 110 | 103 | 91 | 97 | 100 | 76 | 109 | 106 |
| change in I50 | 100 | 89 | 93 | 98 | 121 | 82 | 75 | 52 | 116 |

Supplementary Table 7: Parameters of dose-response curve fits of Opto-T7RNAP*(563) (Wild Type; WT) and nMagHigh1 variants, grown for 13h at 30°C and measured through spectrophotometry. Fitting was performed as described in the Materials and Methods section.

| nMagHigh1 variants at 30°C | WT | L39P | K137E | A143S | V147M | V160M | T38A L39R | M48V N100L | I135V K137R |
|---|---|---|---|---|---|---|---|---|---|
| Best-fit values | | | | | | | | | |
| Basal expression (*b*) a.u. | 944.2 | 1939 | 1275 | 524.7 | 441.7 | 1486 | 1033 | 3591 | 476.9 |
| Maximal expression (*t*) a.u. | 14995 | 17252 | 16329 | 13107 | 14849 | 15913 | 10982 | 17311 | 14767 |
| Half-maximal intensity (*I50*) W $cm^{-2}$ | 0.15 | 0.12 | 0.14 | 0.18 | 0.22 | 0.12 | 0.12 | 0.08 | 0.19 |
| Hillslope (n) | 1.5 | 1.6 | 1.6 | 1.7 | 1.7 | 1.7 | 1.6 | 1.2 | 1.6 |
| Lower 95% conf. limit (profile likelihood) | | | | | | | | | |
| Basal expression (*b*) a.u. | 277 | 1370 | 704.4 | 32.33 | -456.5 | 941 | 488.6 | 2543 | -112.5 |
| Maximal expression (*t*) a.u. | 13863 | 16476 | 15492 | 12358 | 13365 | 15128 | 10250 | 15919 | 13805 |
| Half-maximal intensity (*I50*) W $cm^{-2}$ | 0.13 | 0.10 | 0.13 | 0.16 | 0.18 | 0.11 | 0.10 | 0.05 | 0.17 |
| Hillslope (n) | 1.1 | 1.3 | 1.3 | 1.4 | 1.3 | 1.3 | 1.1 | 0.5 | 1.3 |
| Upper 95% conf. limit (profile likelihood) | | | | | | | | | |
| Basal expression (*b*) a.u. | 1611 | 2508 | 1846 | 1016 | 1335 | 2031 | 1578 | 4639 | 1066 |
| Maximal expression (*t*) a.u. | 16756 | 18271 | 17442 | 14094 | 17268 | 16966 | 12156 | 22409 | 16105 |
| Half-maximal intensity (*I50*) W $cm^{-2}$ | 0.19 | 0.13 | 0.17 | 0.21 | 0.28 | 0.14 | 0.14 | 0.15 | 0.22 |
| Hillslope (n) | 1.9 | 2.0 | 2.0 | 2.0 | 2.3 | 2.1 | 2.2 | 2.3 | 2.0 |
| Goodness of Fit | | | | | | | | | |
| Degrees of Freedom | 40 | 41 | 49 | 50 | 46 | 44 | 46 | 47 | 49 |
| R squared | 0.965 | 0.980 | 0.974 | 0.972 | 0.935 | 0.977 | 0.951 | 0.907 | 0.968 |
| Comparison to WT (in %) | | | | | | | | | |
| change in b | 100 | 205 | 135 | 56 | 47 | 157 | 109 | 380 | 51 |
| change in t | 100 | 115 | 109 | 87 | 99 | 106 | 73 | 115 | 98 |
| change in I50 | 100 | 75 | 94 | 117 | 143 | 78 | 77 | 51 | 124 |

Supplementary Table 8: Parameters of dose-response curve fits of Opto-T7RNAP*(563) (Wild Type; WT) and pMag variants, grown for 5h at 30°C and measured through flow cytometry. Fitting was performed as described in the Materials and Methods section.

| pMag variants at 30°C | WT | R52L | T69M | T83M | N100K | N100L | N100S | K119N | M165T | S178Y | P42T T69M | P66S Y94N |
|---|---|---|---|---|---|---|---|---|---|---|---|---|
| Best-fit values | | | | | | | | | | | | |
| Basal expression (*b*) a.u. | 12539 | 41431 | 32604 | 30733 | 23084 | 31754 | 14001 | 11358 | 30641 | 22991 | 58643 | 21119 |
| Maximal expression (*t*) a.u. | 128601 | 264567 | 170188 | 161146 | 112736 | 212277 | 145632 | 125617 | 147397 | 174306 | 265693 | 212713 |
| Half-maximal intensity (*I50*) W $cm^{-2}$ | 0.17 | 0.11 | 0.10 | 0.09 | 0.09 | 0.13 | 0.16 | 0.15 | 0.11 | 0.13 | 0.10 | 0.13 |
| Hillslope (n) | 1.3 | 1.5 | 1.5 | 1.6 | 2.5 | 1.3 | 1.2 | 1.5 | 1.5 | 1.2 | 1.4 | 1.5 |
| Lower 95% conf. limit (profile likelihood) | | | | | | | | | | | | |
| Basal expression (*b*) a.u. | 9720 | 33344 | 24785 | 26645 | 18332 | 24292 | 8821 | 6529 | 23695 | 18660 | 51504 | 12309 |
| Maximal expression (*t*) a.u. | 122353 | 252321 | 159804 | 155571 | 108973 | 197244 | 134022 | 117411 | 138046 | 164868 | 254858 | 199673 |
| Half-maximal intensity (*I50*) W $cm^{-2}$ | 0.16 | 0.10 | 0.08 | 0.08 | 0.08 | 0.11 | 0.13 | 0.13 | 0.09 | 0.12 | 0.09 | 0.11 |
| Hillslope (n) | 1.1 | 1.1 | 0.9 | 1.2 | 1.8 | 0.9 | 0.9 | 1.1 | 1.0 | 1.0 | 1.1 | 1.2 |
| Upper 95% conf. limit (profile likelihood) | | | | | | | | | | | | |
| Basal expression (*b*) a.u. | 15357 | 49517 | 40422 | 34820 | 27836 | 39213 | 19178 | 16182 | 37587 | 27321 | 65782 | 29925 |
| Maximal expression (*t*) a.u. | 136930 | 281954 | 189266 | 168629 | 117505 | 239148 | 166200 | 137897 | 164315 | 188336 | 281420 | 232156 |
| Half-maximal intensity (*I50*) W $cm^{-2}$ | 0.20 | 0.13 | 0.12 | 0.10 | 0.10 | 0.17 | 0.21 | 0.18 | 0.13 | 0.16 | 0.11 | 0.15 |
| Hillslope (n) | 1.5 | 1.9 | 2.3 | 2.0 | 4.7 | 1.7 | 1.6 | 1.9 | 2.2 | 1.5 | 1.8 | 2.0 |
| Goodness of Fit | | | | | | | | | | | | |
| Degrees of Freedom | 14 | 14 | 14 | 14 | 14 | 14 | 14 | 14 | 14 | 14 | 14 | 14 |
| R squared | 0.997 | 0.994 | 0.986 | 0.996 | 0.989 | 0.991 | 0.992 | 0.991 | 0.984 | 0.996 | 0.995 | 0.990 |
| Comparison to WT (in %) | | | | | | | | | | | | |
| change in b | 100 | 330 | 260 | 245 | 184 | 253 | 112 | 91 | 244 | 183 | 468 | 168 |
| change in t | 100 | 206 | 132 | 125 | 88 | 165 | 113 | 98 | 115 | 136 | 207 | 165 |
| change in I50 | 100 | 63 | 55 | 54 | 54 | 75 | 90 | 87 | 61 | 77 | 56 | 74 |

Supplementary Table 9: Parameters of dose-response curve fits of Opto-T7RNAP*(563) (Wild Type; WT) and nMagHigh1 variants, grown for 15h for N100L and 13h for all other variants at 30°C and measured through spectrophotometry. Fitting was performed as described in the Materials and Methods section.

| pMag variants at 30°C | WT | R52L | T69M | T83M | N100K | N100L | N100S | K119N | M165T | S178Y | P42T T69M | P66S Y94N |
|---|---|---|---|---|---|---|---|---|---|---|---|---|
| Best-fit values | | | | | | | | | | | | |
| Basal expression (*b*) a.u. | 944.2 | 4055 | 3075 | 2797 | 2049 | 3412 | 1338 | 1023 | 2821 | 2298 | 6849 | 2596 |
| Maximal expression (*t*) a.u. | 14995 | 29401 | 22549 | 21261 | 14750 | 20904 | 16109 | 14050 | 17345 | 20888 | 28763 | 25232 |
| Half-maximal intensity (*I50*) W $cm^{-2}$ | 0.15 | 0.08 | 0.08 | 0.08 | 0.09 | 0.08 | 0.13 | 0.15 | 0.11 | 0.11 | 0.04 | 0.12 |
| Hillslope (n) | 1.5 | 2.2 | 1.9 | 1.6 | 1.9 | 2.8 | 1.7 | 1.6 | 1.5 | 1.5 | 1.2 | 1.9 |
| Lower 95% conf. limit (profile likelihood) | | | | | | | | | | | | |
| Basal expression (*b*) a.u. | 277 | 2434 | 2279 | 2082 | 669.5 | 1628 | 863.7 | 559 | 2176 | 1657 | 5164 | 1800 |
| Maximal expression (*t*) a.u. | 13863 | 28111 | 21756 | 20447 | 13511 | 19754 | 15470 | 13376 | 16433 | 19981 | 26814 | 24357 |
| Half-maximal intensity (*I50*) W $cm^{-2}$ | 0.13 | 0.06 | 0.07 | 0.07 | 0.06 | 0.06 | 0.12 | 0.13 | 0.09 | 0.09 | 0.01 | 0.11 |
| Hillslope (n) | 1.1 | 1.2 | 1.3 | 1.2 | 0.8 | 1.2 | 1.4 | 1.4 | 1.1 | 1.2 | 0.1 | 1.6 |
| Upper 95% conf. limit (profile likelihood) | | | | | | | | | | | | |
| Basal expression (*b*) a.u. | 1611 | 5677 | 3870 | 3511 | 3428 | 5196 | 1812 | 1486 | 3465 | 2939 | 8535 | 3392 |
| Maximal expression (*t*) a.u. | 16756 | 31442 | 23617 | 22384 | 17719 | 22931 | 16900 | 14924 | 18754 | 22122 | n.d. | 26284 |
| Half-maximal intensity (*I50*) W $cm^{-2}$ | 0.19 | n.d. | 0.09 | 0.09 | 0.13 | 0.10 | 0.15 | 0.16 | 0.13 | 0.12 | n.d. | 0.13 |
| Hillslope (n) | 1.9 | n.d. | 2.6 | 2.2 | n.d. | n.d. | 2.0 | 2.0 | 2.0 | 1.9 | n.d. | 2.3 |
| Goodness of Fit | | | | | | | | | | | | |
| Degrees of Freedom | 40 | 49 | 50 | 50 | 49 | 50 | 50 | 49 | 49 | 50 | 50 | 50 |
| R squared | 0.965 | 0.939 | 0.973 | 0.975 | 0.838 | 0.860 | 0.982 | 0.978 | 0.965 | 0.979 | 0.906 | 0.980 |
| Comparison to WT (in %) | | | | | | | | | | | | |
| change in b | 100 | 429 | 326 | 296 | 217 | 361 | 142 | 108 | 299 | 243 | 725 | 275 |
| change in t | 100 | 196 | 150 | 142 | 98 | 139 | 107 | 94 | 116 | 139 | 192 | 168 |
| change in I50 | 100 | 49 | 53 | 54 | 61 | 54 | 87 | 94 | 69 | 69 | 26 | 75 |

Supplementary Table 10: Fluorescence (mCherry) expression of WT and nMagHigh1 variants at 40°C. Shown is the mean fluorescence of six biological replicates (n=3) and percentage of fluorescence from the different variants in comparison with the WT.

| 40°C | WT | L39P | | K137E | | A143S | | V147M | | V160M | | T38A L39R | | M48V N100L | | I135V K137R | |
|---|---|---|---|---|---|---|---|---|---|---|---|---|---|---|---|---|---|
| light intensity (W cm-2) | Fluo AVG (a.u.) | Fluo AVG (a.u.) | % | Fluo AVG (a.u.) | % | Fluo AVG (a.u.) | % | Fluo AVG (a.u.) | % | Fluo AVG (a.u.) | % | Fluo AVG (a.u.) | % | Fluo AVG (a.u.) | % | Fluo AVG (a.u.) | % |
| 0 | 82 | 96 | 118 | 92 | 113 | 85 | 105 | 80 | 98 | 90 | 110 | 84 | 103 | 135 | 165 | 83 | 102 |
| 0.48 | 128 | 189 | 148 | 173 | 135 | 103 | 81 | 93 | 72 | 201 | 157 | 128 | 100 | 438 | 342 | 102 | 80 |
| 0.96 | 176 | 246 | 140 | 226 | 128 | 134 | 76 | 123 | 70 | 260 | 147 | 157 | 89 | 604 | 343 | 135 | 77 |
| 1.92 | 223 | 337 | 151 | 303 | 136 | 172 | 77 | 152 | 68 | 368 | 165 | 205 | 92 | 824 | 369 | 175 | 78 |
| 3.85 | 252 | 364 | 144 | 317 | 126 | 194 | 77 | 176 | 70 | 396 | 157 | 221 | 88 | 878 | 348 | 202 | 80 |
| 5.77 | 305 | 474 | 155 | 389 | 127 | 250 | 82 | 240 | 79 | 517 | 169 | 270 | 89 | 1256 | 412 | 260 | 85 |

Supplementary Table 11: Fluorescence (mCherry) expression of WT and pMag variants at 40°C. Shown is the mean fluorescence of six biological replicates (n=3) and percentage of fluorescence from the different variants in comparison with the WT.

| 40°C | WT | R52L | | T69M | | T83M | | N100K | | N100L | | N100S | | K119N | | M165T | |
|---|---|---|---|---|---|---|---|---|---|---|---|---|---|---|---|---|---|
| light intensity (W cm-2) | Fluo AVG (a.u.) | Fluo AVG (a.u.) | % | Fluo AVG (a.u.) | % | Fluo AVG (a.u.) | % | Fluo AVG (a.u.) | % | Fluo AVG (a.u.) | % | Fluo AVG (a.u.) | % | Fluo AVG (a.u.) | % | Fluo AVG (a.u.) | % |
| 0 | 82 | 124 | 153 | 134 | 164 | 118 | 144 | 105 | 129 | 129 | 159 | 98 | 120 | 96 | 118 | 143 | 175 |
| 0.48 | 128 | 232 | 181 | 426 | 332 | 365 | 285 | 275 | 215 | 856 | 668 | 172 | 135 | 141 | 110 | 378 | 295 |
| 0.96 | 176 | 333 | 189 | 593 | 337 | 491 | 279 | 343 | 195 | 944 | 536 | 229 | 130 | 189 | 107 | 498 | 283 |
| 1.92 | 223 | 436 | 195 | 830 | 372 | 699 | 313 | 485 | 218 | 1442 | 647 | 306 | 137 | 219 | 98 | 612 | 274 |
| 3.85 | 252 | 520 | 206 | 944 | 374 | 746 | 296 | 550 | 218 | 1634 | 648 | 345 | 137 | 250 | 99 | 643 | 255 |
| 5.77 | 305 | 655 | 215 | 1243 | 407 | 1065 | 349 | 726 | 238 | 2386 | 782 | 444 | 145 | 306 | 100 | 777 | 255 |

Supplementary Table 12: Fluorescence (mCherry) expression of WT and pMag variants at 40°C. Shown is the mean fluorescence of six biological replicates (n=3) and percentage of fluorescence from the different variants in comparison with the WT.

| 40°C | WT | S178Y | | P42T T69M | | P66S Y94N | |
|---|---|---|---|---|---|---|---|
| light intensity (W cm-2) | Fluo AVG (a.u.) | Fluo AVG (a.u.) | % | Fluo AVG (a.u.) | % | Fluo AVG (a.u.) | % |
| 0 | 82 | 115 | 141 | 213 | 261 | 132 | 162 |
| 0.48 | 128 | 326 | 254 | 619 | 484 | 180 | 141 |
| 0.96 | 176 | 453 | 257 | 937 | 532 | 258 | 146 |
| 1.92 | 223 | 646 | 290 | 1389 | 623 | 315 | 141 |
| 3.85 | 252 | 777 | 308 | 1720 | 682 | 353 | 140 |
| 5.77 | 305 | 1043 | 342 | 2381 | 781 | 465 | 152 |

Supplementary Table 13: List of primers and plasmids used in this study

| Primer/plasmid name | Sequence (5'-3') |
|---|---|
| oAB507 | AGACTCGAGGGTACCTTATTTGTACAGTTCATCCA |
| oAB707 | CTAACTTACATTAATTGCGTTGCGCTCTAGATTATTTGTATAGTTCATCCATGCCATGTG |
| oAB708 | GGGGACTGTTGGGCGCCATCTCCTTGCATGCACTAGTGTTCTTTCCTGCGTTATCCCCTG |
| oAB819 | AGAGGATCCGAAGAATACGGGATAGAACTGAATGGTTTCAAAAGGCGAAGAAGACAAC |
| oAB734 | GGTAGGTGGTCGCGCGGTTAACTTGCTTCCTGGCGGTTCTGGAGGT |
| oAB736 | GTCTTTCGACTGAGCCTTTCGTTTTATTTGATGCCTCCTAGGTTA |
| oAB810 | AGAAGGAGGTCATACCCGTTTTTTTGGAAGGAGGTAAATTAATTA |
| oAB744 | GACTTTCTTAGCAACAATCCCGTAGATGTCCTGAACGGTTTCACTACCTCCAGAACCGCC |
| oAB589 | TAACCTAGGAGGCATCAAATAAAACG |
| oAB446 | ACCTCCAGAACCGCCAGGAAGCAAGTTAACCGCGCG |
| oAB448 | GGCGGTTCTGGAGGTAGTGAAACCGTTCAGGACATCTACGG |
| oAB809 | TAATTAATTTACCTCCTTCCAAAAAAACGGGTATG |
| pAB150 | as described in Baumschlager et al., 2017 |
| pAB50 | as described in Baumschlager et al., 2017 |
| pAB50-11k | GTCGTTTGGTATGGCTTCATTCAGCTCCGGTTCCCAACGATCAAGGCGAGTTACATGATCCCCC<br>ATGTTGTGCAAAAAAGCGGTTAGCTCCTTCGGTCCTCCGATCGTTGTCAGAAGTAAGTTGGCCG<br>CAGTGTTATCACTCATGGTTATGGCAGCACTGCATAATTCTCTTACTGTCATGCCATCCGTAAGA<br>TGCTTTTCTGTGACTGGTGAGTACTCAACCAAGTCATTCTGAGAATAGTGTATGCGGCGACCGA<br>GTTGCTCTTGCCCGGCGTCAATACGGGATAATACCGCGCCACATAGCAGAACTTTAAAAGTGCT<br>CATCATTGGAAAACGTTCTTCGGGGCGAAAACTCTCAAGGATCTTACCGCTGTTGAGATCCAGT<br>TCGATGTAACCCACTCGTGCACCCAACTGATCTTCAGCATCTTTTACTTTCACCAGCGTTTCTGG<br>GTGAGCAAAAACAGGAAGGCAAAATGCCGCAAAAAAGGGAATAAGGGCGACACGGAAATGTT<br>GAATACTCATACTCTTCCTTTTTCAATCATGATTGAAGCATTTATCAGGGTTATTGTCTCATGAG<br>CGGATACATATTTGAATGTATTTAGAAAAATAAACAAATAGGTCATGACCAAAATCCCTTAACG<br>TGAGTTTTCGTTCCACTGAGCGTCAGACCCCGTAGAAAAGATCAAAGGATCTTCTTGAGATCCT<br>TTTTTTCTGCGCGTAATCTGCTGCTTGCAAACAAAAAAACCACCGCTACCAGCGGTGGTTTGTTT<br>GCCGGATCAAGAGCTACCAACTCTTTTTCCGAAGGTAACTGGCTTCAGCAGAGCGCAGATACCA<br>AATACTGTCCTTCTAGTGTAGCCGTAGTTAGGCCACCACTTCAAGAACTCTGTAGCACCGCCTA<br>CATACCTCGCTCTGCTAATCCTGTTACCAGTGGCTGCTGCCAGTGGCGATAAGTCGTGTCTTACC<br>GGGTTGGACTCAAGACGATAGTTACCGGATAAGGCGCAGCGGTCGGGCTGAACGGGGGGTTCG<br>TGCACACAGCCCAGCTTGGAGCGAACGACCTACACCGAACTGAGATACCTACAGCGTGAGCTA<br>TGAGAAAGCGCCACGCTTCCCGAAGGGAGAAAGGCGGACAGGTATCCGGTAAGCGGCAGGGT<br>CGGAACAGGAGAGCGCACGAGGGAGCTTCCAGGGGGAAACGCCTGGTATCTTTATAGTCCTGT<br>CGGGTTTCGCCACCTCTGACTTGAGCGTCGATTTTTGTGATGCTCGTCAGGGGGGCGGAGCCTA<br>TGGAAAAACGCCAGCAACGCGGCCTTTTTACGGTTCCTGGCCTTTTGCTGGCCTTTTGCTCACAT<br>GTTCTTTCCTGCGTTATCCCCTGATTCTGTGGATAACCGTATTACCGCCTTTGAGTGAGCTGATA<br>CCGCTCGCCGCAGCCGAACGACCGAGCGCAGCGAGTCAGTGAGCGAGGAAGCGGAAGAGCGC<br>CTGATGCGGTATTTTCTCCTTACACATCTGTGCGGTATTTCACACCGCATATATGGTGCACTCTC<br>AGTACAATCTGCTCTGATGCCGCATAGTTAAGCCAGTATACACTCCGCTATCGCTACGTGACTG<br>GGTCATGGCTGCGCCCCGACACCCGCCAACACCCGCTGACGCGCCCTGACGGGCTTGTCTGCTC<br>CCGGCATCCGCTTACAGACAAGCTGTGACCGTCTCCGGGAGCTGCATGTGTCAGAGGTTTTCAC<br>CGTCATCACCGAAACGCGCGAGGCAGCTGCGGTAAAGCTCATCAGCGTGGTCGTGAAGCGATT<br>CACAGATGTCTGCCTGTTCATCCGCGTCCAGCTCGTTGAGTTTCTCCAGAAGCGTTAATGTCTGG<br>CTTCTGATAAAGCGGGCCATGTTAAGGGCGGTTTTTTCCTGTTTGGTCACTGATGCCTCCGTGTA<br>AGGGGGATTTCTGTTCATGGGGGTAATGATACCGATGAAACGAGAGAGGATGCTCACGATACG<br>GGTTACTGATGATGAACATGCCCGGTTACTGGAACGTTGTGAGGGTAAACAACTGGCGGTATG<br>GATGCGGCGGGACCAGAGAAAAATCACTCAGGGTCAATGCCAGCGCTTCGTTAATACAGATGT<br>AGGTGTTCCACAGGGTAGCCAGCAGCATCCTGCGATGCAGATCCGGAACATAATGGTGCAGGG<br>CGCTGACTTCCGCGTTTCCAGACTTTACGAAACACGGAAACCGAAGACCATTCATATTGTTGCT<br>CAGGTCGCAGACGTTTTGCAGCAGCAGTCGCTTCACGTTCGCTCGCGTATCGGTGATTCATTCTG<br>CTAACCAGTAAGGCAACCCCGCCAGCCTAGCCGGGTCCTCAACGACAGGAGCACGATCATGCT<br>AGTCATGCCCCGCGCCCACCGGAAGGAGCTGACTGGGTTGAAGGCTCTCAAGGGCATCGGTCG<br>AGATCCCGGTGCCTAATGAGTGAGCTAACTTACATTAATTGCGTTGCGCTCTAGATTATTTGTAT<br>AGTTCATCCATGCCATGTGTAATCCCAGCAGCTGTTACAAACTCAAGAAGGACCATGTGGTCTC<br>TCTTTTCGTTGGGATCTTTCGAAAGGGCAGATTGTGTGGACAGGTAATGGTTGTCTGGTAAAAG<br>GACAGGGCCATCGCCAATTGGAGTATTTTGTTGATAATGGTCTGCTAGTTGAACGCTTCCATCTT<br>CAATGTTGTGTCTAATTTTGAAGTTAACTTTGATTCCATTCTTTTGTTTGTCTGCCATGATGTATA<br>CATTGTGTGAGTTATAGTTGTATTCCAATTTGTGTCCAAGAATGTTTCCATCTTCTTTAAAATCA<br>ATACCTTTTAACTCGATTCTATTAACAAGGGTATCACCTTCAAACTTGACTTCAGCACGTGTCTT<br>GTAGTTCCCGTCATCTTTGAAAAATATAGTTCTTTCCTGTACATAACCTTCGGGCATGGCACTCT |

| | |
|---|---|
| | TGAAAAAGTCATGCTGTTTCATATGATCTGGGTATCTCGCAAAGCATTGAACACCATAACCGAA<br>AGTAGTGACAAGTGTTGGCCATGGAACAGGTAGTTTTCCAGTAGTGCAAATAAATTTAAGGGTA<br>AGTTTTCCGTATGTTGCATCACCTTCACCCTCTCCACTGACAGAAAATTTGTGCCCATTAACATC<br>ACCATCTAATTCAACAAGAATTGGGACAACTCCAGTGAAAAGTTCTTCTCCTTTACGCATAGAT<br>CTTTTCCTTATTACGCCTGCTGGCGAAAGGGGGATGTGCTGCAAGGCGATTAAGTTGGGTAACG<br>CCCGGGTTTTACCAGTCACGACGTTGTAAAACGACGGCCAGTGAATCGTGTCAGTGGTGTATCA<br>TTATAGGGAGTTATTCCGGCCTGACAAGAGGAATCAGGGGATAACGCAGGAAAGAACACTAGT<br>GCATGCAAGGAGATGGCGCCCAACAGTCCCCCGGCCACGGGGCCTGCCACCATACCCACGCCG<br>AAACAAGCGCTCATGAGCCCGAAGTGGCGAGCCCGATCTTCCCCATCGGTGATGTCGGCGATAT<br>AGGCGCCAGCAACCGCACCTGTGGCGCCGGTGATGCCGGCCACGATGCGTCCGGCGTAGCCTA<br>GGTAATACGACTCACTATAGGGAGAGGATCCGAAGAATACGGGATAGAACTGAATGGTTTCAA<br>AAGGCGAAGAAGACAACATGGCGATTATCAAGGAATTTATGCGTTTCAAGGTCCACATGGAAG<br>GCAGCGTCAATGGTCACGAATTTGAAATTGAAGGCGAAGGTGAAGGCCGTCCGTATGAAGGCA<br>CCCAGACGGCAAAACTGAAGGTCACCAAAGGCGGTCCGCTGCCGTTTGCTTGGGATATTCTGTC<br>ACCGCAATTCATGTATGGTTCGAAAGCGTACGTTAAGCATCCGGCCGATATCCCGGACTATCTG<br>AAACTGTCCTTTCCGGAAGGCTTCAAATGGGAACGTGTTATGAACTTCGAAGATGGCGGTGTGG<br>TTACCGTCACGCAGGATAGCTCTCTGCAAGACGGTGAATTTATTTATAAAGTGAAGCTGCGCGG<br>CACCAATTTCCCGAGCGATGGTCCGGTTATGCAGAAAAAGACGATGGGCTGGGAAGCGAGTTC<br>CGAACGTATGTACCCGGAAGACGGTGCCCTGAAAGGCGAAATCAAGCAGCGCCTGAAACTGAA<br>GGATGGCGGTCACTATGACGCAGAAGTGAAAACCACGTACAAGGCTAAAAAGCCGGTCCAACT<br>GCCGGGTGCATACAACGTGAACATCAAGCTGGATATCACCAGCCATAACGAAGACTATACGAT<br>CGTTGAACAGTACGAACGTGCAGAAGGCCGCCACTCTACCGGCGGTATGGATGAACTGTACAA<br>ATAAGGTACCCTCGAGTCTGGTAAAGAAACCGCTGCTGCGAAATTTGAACGCCAGCACATGGA<br>CTCGTCTACTAGTCGCAGCTTAATTAACCTAAACTGCTGCCACCGCTGAGCAATAACTAGCATA<br>ACCCCTTGGGGCCTCTAAACGGGTCTTGAGGGGTTTTTTGCTAGCGAAAGGAGGAGTCGACTAT<br>ATCCGGATTGGCGAATGGGACGCGCCCTGTAGCGGCGCATTAAGCGCGGCGGGTGTGGTGGTT<br>ACGCGCAGCGTGACCGCTACACTTGCCAGCGCCCTAGCGCCCGCTCCTTTCGCTTTCTTCCCTTC<br>CTTTCTCGCCACGTTCGCCGGCTTTCCCCGTCAAGCTCTAAATCGGGGGCTCCCTTTAGGGTTCC<br>GATTTAGTGCTTTACGGCACCTCGACCCCAAAAAACTTGATTAGGGTGATGGTTCACGTAGTGG<br>GCCATCGCCCTGATAGACGGTTTTTCGCCCTTTGACGTTGGAGTCCACGTTCTTTAATAGTGGAC<br>TCTTGTTCCAAACTGGAACAACACTCAACCCTATCTCGGTCTATTCTTTTGATTTATAAGGGATT<br>TTGCCGATTTCGGCCTATTGGTTAAAAAATGAGCTGATTTAACAAAAATTTAACGCGAATTTTA<br>ACAAAATATTAACGTTTACAATTTCTGGCGGCACGATGGCATGAGATTATCAAAAAGGATCTTC<br>ACCTAGATCCTTTTAAATTAAAAATGAAGTTTTAAATCAATCTAAAGTATATATGAGTAAACTT<br>GGTCTGACAGTTACCAATGCTTAATCAGTGAGGCACCTATCTCAGCGATCTGTCTATTTCGTTCA<br>TCCATAGTTGCCTGACTCCCCGTCGTGTAGATAACTACGATACGGGAGGGCTTACCATCTGGCC<br>CCAGTGCTGCAATGATACCGCGAGACCCACGCTCACCGGCTCCAGATTTATCAGCAATAAACCA<br>GCCAGCCGGAAGGGCCGAGCGCAGAAGTGGTCCTGCAACTTTATCCGCCTCCATCCAGTCTATT<br>AATTGTTGCCGGGAAGCTAGAGTAAGTAGTTCGCCAGTTAATAGTTTGCGCAACGTTGTTGCCA<br>TTGCTACAGGCATCGTGGTGTCACGCTC |
